# Ref(2)P/p62 protein pleomorphism confers tissue-specific function

**DOI:** 10.64898/2026.01.10.696886

**Authors:** Yan Yan, Kai Zhang, Gabrielle Pycior, Pablo Adrian Guillen-Poza, Ruben Hervas, Paulo Leal, Kexi Yi, Zulin Yu, Ying Zhang, Mansoor Hussain, Laurence Florens, Jay Unruh, Kausik Si

## Abstract

Protein structure dictates function and proteins often perform the same function in different cell or tissue-type. Here, we report that in *Drosophila melanogaster*, the broadly expressed protein Ref(2)P, an orthologue of mammalian p62/SQSTM1, performs mutually exclusive functions across tissues – autophagy in midgut and mitochondria-associated proteostasis in muscle. These functional differences arise from tissue-specific secondary and tertiary structures adopted by the same Ref(2)P polypeptide, rather than protein abundance, isoforms, post-translational modification, or binding-partner availability. The tissue-specific structures are cell autonomous and resistant to environmental perturbation. Deletion of a short intrinsically disordered region converts muscle Ref(2)P to a midgut-like structural and functional state. However, this forced switch causes muscle dysfunction, linking tissue-specific physiology to alternative structural states of the same polypeptide. We posit that *Drosophila* Ref(2)P exemplifies a mechanism for tissue-specificity, protein pleomorphism, in which the intrinsic structural plasticity of a protein and the physicochemical/molecular environment of a tissue enable a broadly expressed protein to adopt tissue-specific structure and function.

## MAIN

Each tissue acquires a functional identity and maintains it even as conditions change. Tissue-specific functions are primarily attributed to differentially expressed proteins and proteins that are broadly expressed are generally assumed to carry out the same function in every tissue ^1^. However, tissues of distinct developmental origin and function share a large part of the proteome and whether these shared proteins contribute to tissue-specific biology and if so, how, remain unexplored.

Structure endows proteins with their biochemical properties and biological functions. Most proteins adopt a specific structure to perform a stereotyped function regardless of cell or tissue types. However, emerging evidence suggests the inherent properties of polypeptides can allow a single protein to adopt distinct structure and functions. These structural differences could be extreme, adopting different secondary structures such as metamorphic proteins ^2^ and fold- switching proteins ^3^ or different tertiary or quaternary organization of secondary structures ^4,5^. However, less is known about whether these multiple structural states exist in the same cell or in a mutually exclusive manner, and more importantly, whether such states contribute to tissue-identity or tissue-specific function.

Cellular proteostasis is maintained by multiple, partially independent degradation pathways, including ubiquitin–proteasome degradation, autophagy–lysosomal degradation, and mitochondrial import–associated degradation pathways ^6,7^. Across these systems, substrate specificity often relies on two recurring steps: (i) cargo marking, frequently through ubiquitination, and (ii) cargo handoff/delivery, mediated by ubiquitin-binding adaptors or receptors that couple marked cargo to the appropriate degradation machinery ^8,9^. Notably, the number of such adaptors varies widely across species, yet these core degradation pathways are conserved ^10–12^. This raises the question how ubiquitinated substrates are partitioned among distinct degradation pathways.

Ref(2)P, the *Drosophila* orthologue of mammalian p62/SQSTM1, is a broadly expressed, multidomain and multifunctional adaptor that binds ubiquitinated cargos and delivers them to degradation machinery. Ref(2)P domain organization is conserved from insects to mammals: an N-terminal Phox and Bem1 (PB1) domain that mediates self-polymerization ^13^, a ZZ-type zinc finger ^14^, a short Atg8-interacting motif (AIM; the LC3-interacting region, LIR, in mammals) ^15^ and a C-terminal ubiquitin-associated (UBA) domain ^16^. Mammalian p62/SQSTM1 tethers ubiquitinated substrates to Atg8/LC3 on the phagophore ^17,18^ and PB1-driven polymerization concentrate cargo into micrometer-scale condensates ^19,20^. *Drosophila* Ref(2)P was initially identified through its requirement for sigma rhabdovirus multiplication and male fertility ^21^ and its function is best characterized in tissues with high autophagic activity, particularly the midgut, which undergoes a robust, developmentally programmed autophagic destruction ^22,23,24,25,26^. In addition to its autophagic function, Ref(2)P/p62 also scaffolds signaling complexes that control the Keap1–Nrf2 oxidative-stress response ^27^ and nutrient sensing by mTORC1 ^28^. Despite its central role in quality control pathway, it remains unclear how these multiple functions of Ref(2)P are coordinated in various tissues.

In this study, we find that Ref(2)P adopts distinct tissue-specific structures that enable a single adaptor to route ubiquitinated cargo to mutually exclusive and tissue-specific degradation pathway: autophagy in the midgut but mitochondrial import–associated degradation in muscle. Our finding implies that tissue environment can drive a protein to adopt tissue-specific structures and consequently restrict the protein to one of the available functions that is tied to the physiology of the tissue.

## *Drosophila* Ref(2)P protein displays distinct localization and binding activities across different tissues

To determine how Ref(2)P engages proteostatic processes across tissues, we examined the localization of Ref(2)P in cell lines and in adult fly tissues, using antibodies against untagged endogenous Ref(2)P and anti-Flag antibodies against Ref(2)P-3xFlag knock-in allele (Extended Data Fig. 1 and 2). In every cell type examined, Ref(2)P formed puncta that colocalized with ubiquitin (Fig. 1a-1d and Extended Data Fig. 1). In cell lines Ref(2)P colocalized with autophagic markers consistent with its well characterized role in autophagy (Extended Data Fig. 1j-1l). Surprisingly, in adult fly tissue, ubiquitin/ Ref(2)P double-positive puncta localized to different cellular compartments in a mutually exclusive manner: canonical autophagic structures in the adult midgut (Fig. 1b and Extended Data Fig. 1c, 1h), mitochondria in the indirect flight muscle (Fig. 1c and Extended Data Fig. 1b, 1g), and cytosolic granules devoid of autophagic or mitochondrial markers in oenocytes, the fly liver-like tissue (Fig. 1d and Extended Data Fig. 1f, 1i). Analysis of multiple animals confirmed that tissue-specific localization of Ref(2)P is a general phenomenon (Fig. 1e).

**Fig. 1:**
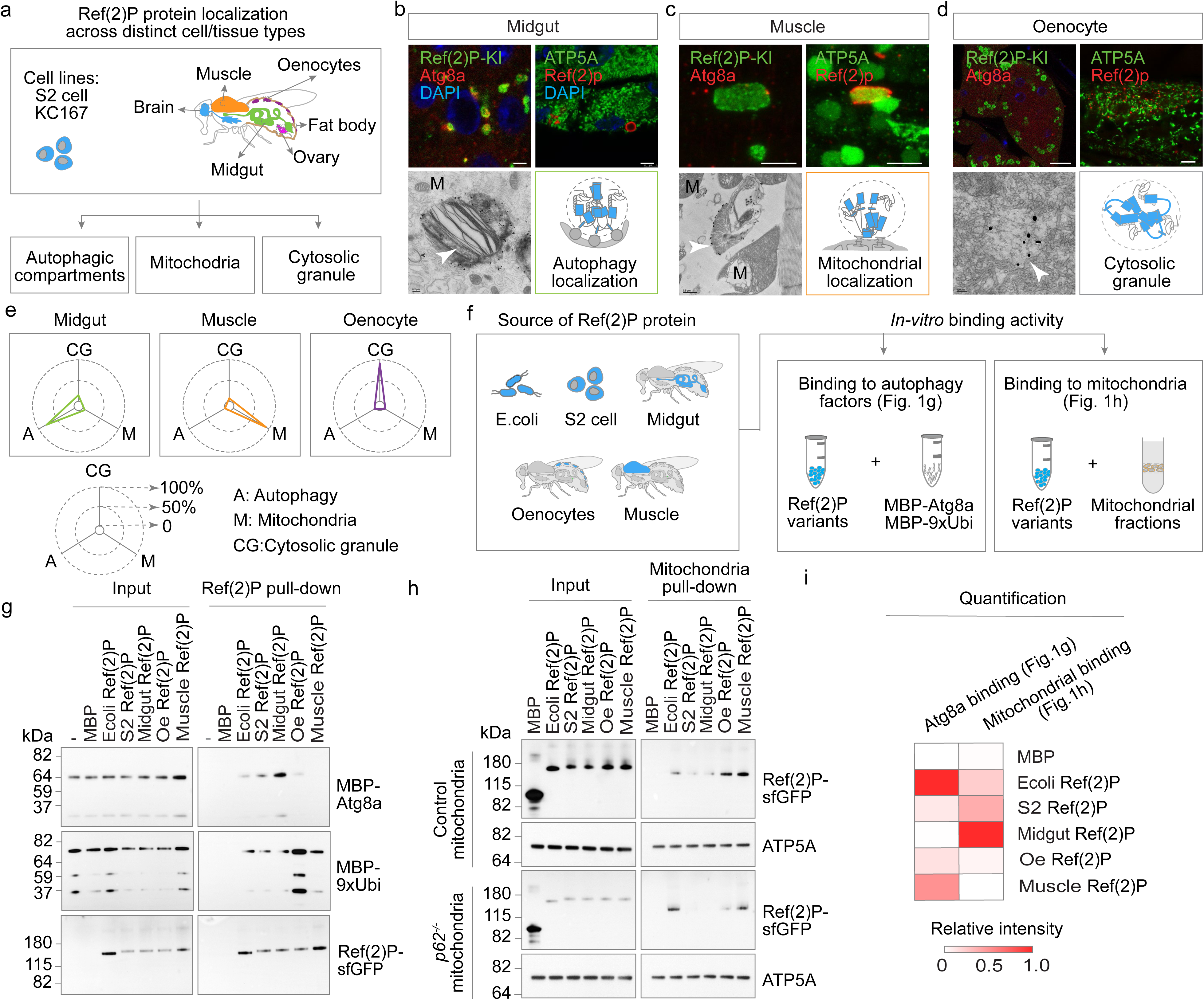
*Drosophila* Ref(2)P protein displays distinct localization and binding activities across different tissues. **a**, Summary of Ref(2)P localization surveyed across *Drosophila* cell lines (S2, KC167) and adult tissues (brain, muscle, midgut, ovary, fat body and oenocytes). **b–d**, Localization of Ref(2)P in adult midgut (b), indirect flight muscle (c) and oenocytes (d). Top left, immunofluorescence of Ref(2)P-3xFlag knock-in flies stained with anti-Flag (Ref(2)P-KI, green) and anti-Atg8a (red) to label autophagic compartments; DAPI (blue). Top right, wild-type flies stained with anti-ATP5A (green) to label mitochondria and anti-Ref(2)P (red). Bottom left, silver-enhanced immuno- electron microscopy of wild-type tissue stained with anti-Ref(2)P; arrowheads, Ref(2)P signal; M, mitochondria. Bottom right, schematic of the compartment occupied by Ref(2)P in that tissue; the same depiction is used throughout the paper. Scale bars 5 µm. **e**, Radar plots quantifying the distribution of Ref(2)P/ubiquitin puncta between autophagic compartments (A), mitochondria (M) and cytosolic granules (CG) in midgut, muscle and oenocytes of 1-week-old flies under normal conditions. Values are the mean percentage of colocalization; n ≥ 9 flies per tissue. Autophagic compartments were marked with anti-Atg8a and mitochondria with anti-ATP5A. **f**, Strategy for testing the in-vitro binding of Ref(2)P protein. Ref(2)P-sfGFP-Flag was expressed in, and purified from, five sources — *E. coli*, S2 cells, midgut, oenocytes and muscle — and assayed *in vitro* for binding to autophagy-pathway factors (g) and to isolated mitochondria (h). **g**, Binding of purified Ref(2)P to recombinant MBP-Atg8a and MBP-9xUbi. Proteins were mixed, recovered with a GFP nanobody and analyzed by western blotting with the indicated antibodies. A no-protein reaction and MBP alone served as controls; 2% of the input was loaded. **h**, Binding of purified Ref(2)P to mitochondria isolated from control (*w1118*) or Ref(2)P-null (Ref(2)P^-/-^)flies. Mitochondria-bound Ref(2)P was assessed by western blotting; anti-ATP5A detects the mitochondrial fraction and anti-GFP detects MBP-GFP (control) and Ref(2)P-sfGFP. 2% of the input was loaded. **i**, Quantification of g and h. Heat map of relative binding intensity (each row scaled to its strongest signal) for Atg8a binding (top) and mitochondrial binding (bottom) across the five Ref(2)P sources.

What determines this tissue-specific distribution? We reasoned that this could be achieved in multiple, but not mutually exclusive, ways. First, it could be due to tissue-specific variants of Ref(2)P protein. However, expression of the same Ref(2)P coding sequence (Ref(2)P-WT-sfGFP) recapitulates the tissue-specific localization and rescues the tissue-specific function of Ref(2)P in recruiting ubiquitin to specific compartments under Ref(2)P RNAi background (Extended Data Fig. 5e-5h). Second, it could be due to the differential expression of either Ref(2)P protein itself or Ref(2)P cofactors. To this end, we measured Ref(2)P protein abundance by western blotting and the abundance of cofactors by tissue-specific RNA sequencing and quantitative proteomics (Extended Data Fig. 3). Autophagy and mitochondrial factors were expressed at comparable levels in both tissues, and Ref(2)P level was slightly higher in midgut compared to muscle (Extended Data Fig. 3b-3c). Raising Ref(2)P protein level in muscle, either by overexpression (Extended Data Fig. 5g) or by heat shock (Fig. 2g and Extended Data Fig. 6), did not move it to autophagic structures, suggesting Ref(2)P protein abundance is not correlated with its differential localization.

**Fig. 2:**
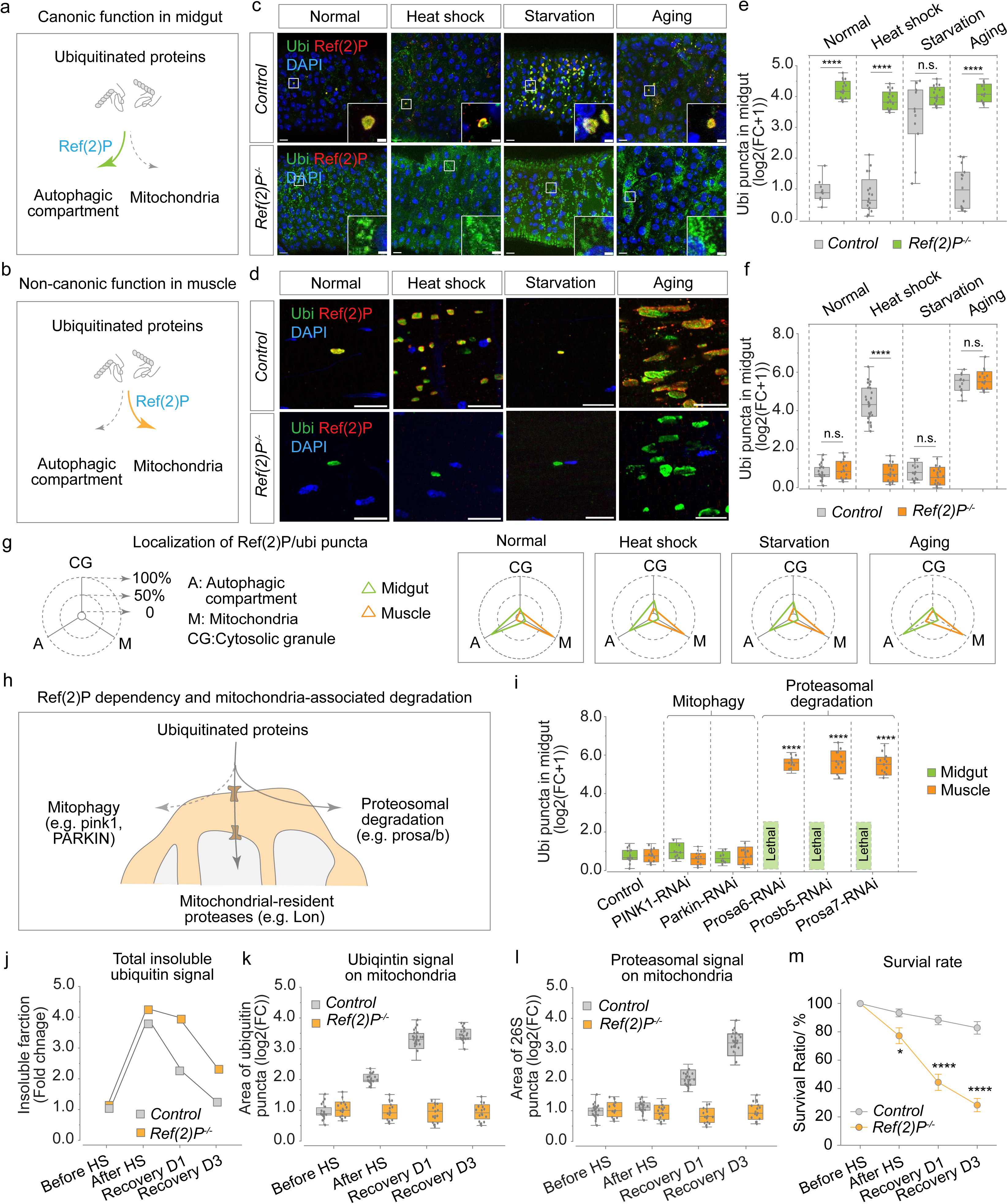
Ref(2)P mediates autophagic degradation in midgut and mitochondria-associated degradation in muscle. **a**, **b**, Schematic of the two mutually exclusive routes tested: the canonical function in midgut, in which Ref(2)P delivers ubiquitinated proteins to autophagic compartments (a), and the non- canonical function in muscle, in which Ref(2)P delivers them to mitochondria (b). **c**, **d**, Immunofluorescence of midgut (c) and indirect flight muscle (d) from control and Ref(2)P-null (Ref(2)P^−/−^) flies under normal conditions, after heat shock, after 48 h starvation and at 6 weeks of age. Ubiquitin (green), Ref(2)P (red), DAPI (blue). Scale bars 10 µm; Insets, magnified views of the boxed regions, Scale bars 2 µm. **e, f,** Quantification of ubiquitin puncta in midgut (e) and muscle (f) from c and d, expressed as log2(fold change + 1). n ≥ 8 flies per condition; two-tailed unpaired t-test; *P < 0.05, **P < 0.01, ***P < 0.001, ****P < 0.0001; n.s., not significant. **g**, Radar plots of the distribution of Ref(2)P/ubiquitin puncta between autophagic compartments (A), mitochondria (M) and cytosolic granules (CG) in midgut and muscle under normal, heat-shock, starvation and aged conditions. **h**, The three routes by which mitochondria contribute to protein degradation that were tested genetically: mitophagy (pink1, PARKIN), and proteasomal degradation at the mitochondrial surface (prosa/prosb subunits). **i**, Quantification of ubiquitin puncta (log2(fold change + 1)) in midgut (green) and muscle (orange) upon tissue-specific knockdown of the genes in h. n ≥ 6 flies per condition; statistics as in e. Corresponding images are in Extended Data Fig. 7a-7b. **j**, Total detergent-insoluble ubiquitin signal (fold change relative to before heat shock) in muscle of control and Ref(2)P^−/−^ flies before heat shock, immediately after heat shock, and after 1 and 3 days of recovery. Source blots are in Extended Data Fig. 7d- 7e. **k**, **l**, Area of ubiquitin puncta (k) and of 26S proteasome puncta (l) on muscle mitochondria over the same time course (log2 fold change). Corresponding images are in Extended Data Fig. 7f-7g. **m,** Survival of control and Ref(2)P-null flies subjected to repeated heat shock, scored over the same time course. Data are mean ± s.d. from three independent cohorts; two-tailed unpaired t-test; *P < 0.05, ****P < 0.0001.

**Fig. 3:**
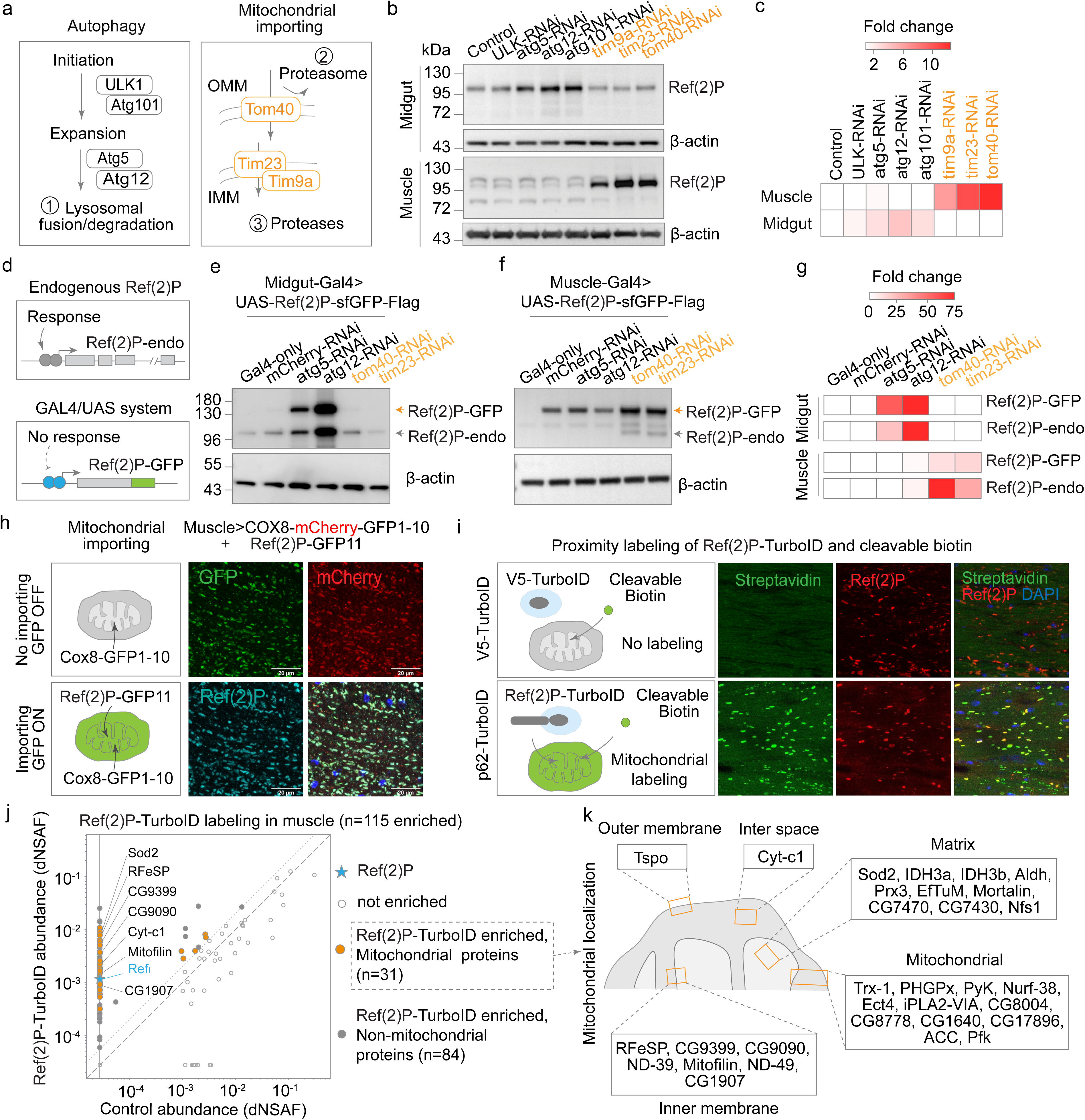
Ref(2)P turnover is differentially regulated by autophagy in the midgut and the mitochondrial import machinery in muscle. **a**, The two degradation routes. Left, autophagy: initiation (ULK1/Atg1, Atg101), phagophore expansion (Atg5, Atg12) and (1) lysosomal fusion and degradation. Right, mitochondrial import: the outer-membrane translocase (Tom40) and the inner-membrane translocases (Tim23, Tim9a), feeding (2) proteasomal and (3) mitochondrial protease-dependent degradation. **b**, Western blot of endogenous Ref(2)P in midgut and muscle of flies expressing the indicated RNAi transgenes under tissue-specific GAL4 control. β-actin, loading control. **c**, Heat map of Ref(2)P fold change quantified from b. Knockdown of autophagy genes raises Ref(2)P only in the midgut; knockdown of mitochondrial import factors raises Ref(2)P only in muscle. **d**, Rationale for separating protein turnover from transcriptional feedback. Endogenous Ref(2)P carries its own regulatory sequences and can respond transcriptionally, whereas the GAL4/UAS-driven Ref(2)P-sfGFP transgene lacks them; a coordinate response of the transgene therefore reports altered turnover. **e-f**, Western blot of tissue lysates from Midgut-GAL4>UAS-Ref(2)P-sfGFP-Flag (e) and Muscle- GAL4>UAS-Ref(2)P-sfGFP-Flag (f) flies carrying the indicated RNAi transgenes. GAL4 alone and mCherry-RNAi are negative controls. Transgenic (Ref(2)P-GFP) and endogenous (Ref(2)P-endo) protein are resolved by size; β-actin, loading control. **g**, Heat map of fold change for Ref(2)P-GFP and Ref(2)P-endo quantified from **e-f**. The transgene and the endogenous protein respond in the same tissue-restricted pattern. **h**, Mitochondrial split-GFP reporter. Muscle cells co-expressed a mitochondria-targeted COX8-mCherry-GFP1-10 together with Ref(2)P tagged with GFP11. Left, schematic: GFP fluorescence is reconstituted only if GFP11-tagged Ref(2)P reaches the compartment that contains GFP1-10. Right, GFP, mCherry, Ref(2)P and merged images. Scale bars as indicated. **i**, Proximity labelling of the Ref(2)P environment in muscle. Ref(2)P was fused to TurboID and biotinylated proteins were captured using a cleavable biotin reagent. Images show streptavidin (green), Ref(2)P (red) and DAPI (blue) in control and Ref(2)P-TurboID muscle. The labelling chemistry and streptavidin-capture quality control are shown in Extended Data Fig. 8c- 8d. **j**, Abundance of proteins recovered from Ref(2)P-TurboID muscle versus control (distributed normalized spectral abundance factor, dNSAF). Star, the Ref(2)P bait; open circles, not enriched; orange, enriched proteins annotated as mitochondrial (n = 31); grey, enriched non-mitochondrial proteins (n = 84). Selected mitochondrial hits are labelled. **k**, Submitochondrial assignment of the 31 enriched mitochondrial proteins: outer membrane (OMM, n = 1), intermembrane space (n = 1), inner membrane (IMM, n = 7), matrix (n = 10) and mitochondrial proteins without a defined submitochondrial annotation (n = 12).

**Fig. 4:**
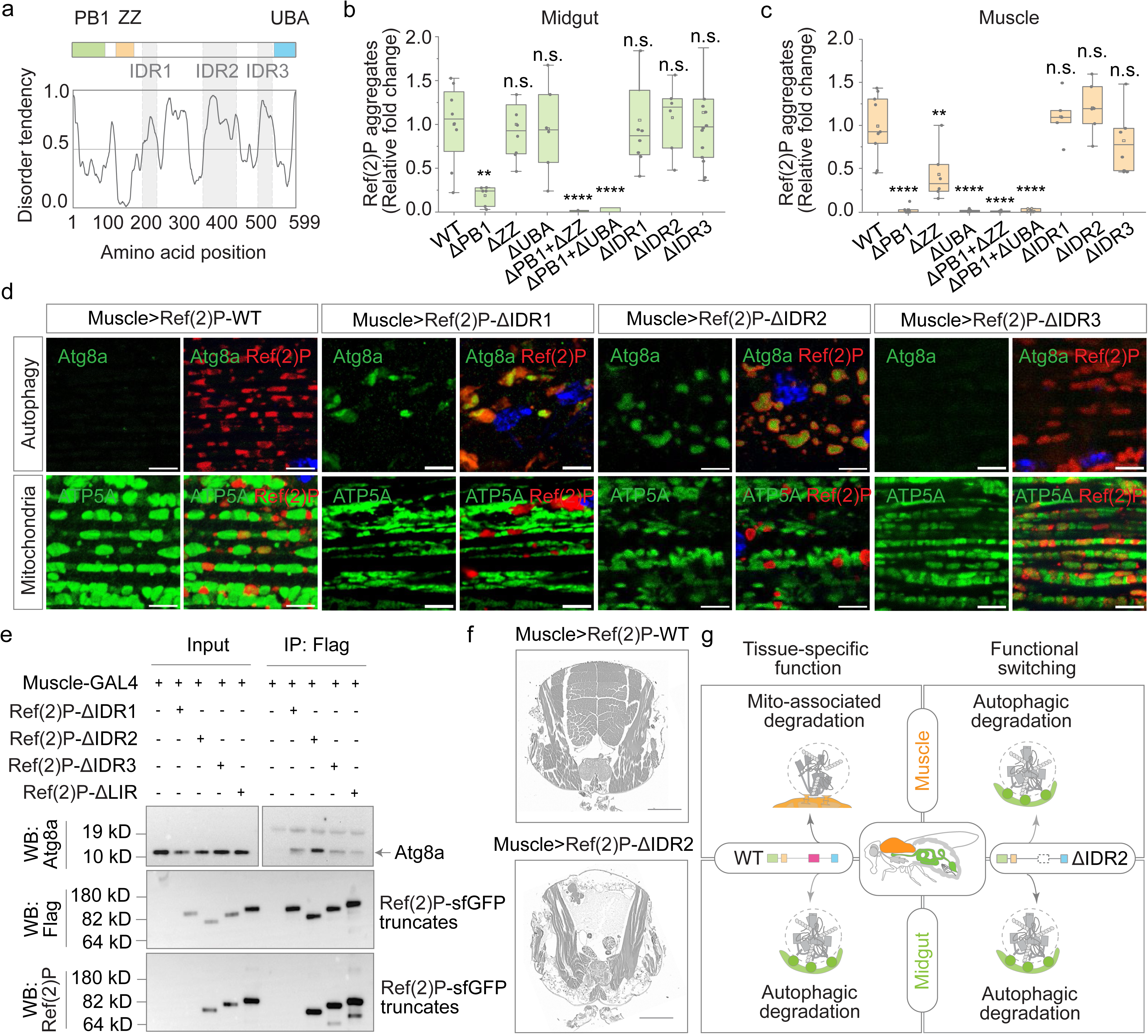
Deletion of a specific IDR switches the tissue-specific function of Ref(2)P. **a**, Domain organization and disorder tendency of *Drosophila* Ref(2)P. The PB1 (green), ZZ (orange) and UBA (blue) domains are indicated; IDR1 (amino acids 192-225), IDR2 (360-449) and IDR3 (503-542) are shaded grey. **b**, **c**, Quantification of Ref(2)P aggregates (relative fold change) in midgut (b) and muscle (c) of flies expressing wild-type or truncated Ref(2)P. n ≥ 6 flies per condition; two-tailed unpaired t-test using the mean value of each animal; **P < 0.01,** **P < 0.0001; n.s., not significant. Corresponding images are in Extended Data Fig. 9a. **d**, Immunofluorescence of muscle expressing Ref(2)P-WT, Ref(2)P-ΔIDR1, Ref(2)P-ΔIDR2 or Ref(2)P-ΔIDR3. Top, Atg8a (green) marks autophagic compartments; bottom, ATP5A (green) marks mitochondria; Ref(2)P (red, anti-Flag). Scale bars 10 µm. **e,** Co-immunoprecipitation from dissected muscle lysates of flies expressing Ref(2)P-ΔIDR1, Ref(2)P-ΔIDR2, Ref(2)P-ΔIDR3 or Ref(2)P-ΔLIR. Anti-Flag was used for immunoprecipitation and anti-Atg8a for western blotting; 2% of the input was loaded. Anti- Flag and anti-Ref(2)P identify the expressed Ref(2)P-sfGFP variants. **f**, Sections of 1-week-old flight muscle from flies expressing Ref(2)P-WT or Ref(2)P-ΔIDR2 under Act88F-GAL4 control. Scale bars, 200 µm. Sections for all IDR deletions are in Extended Data Fig. 9c. **g**, Schematic of the tissue-specific functions of Ref(2)P-WT and of the functional switch caused by deletion of IDR2.

**Fig. 5:**
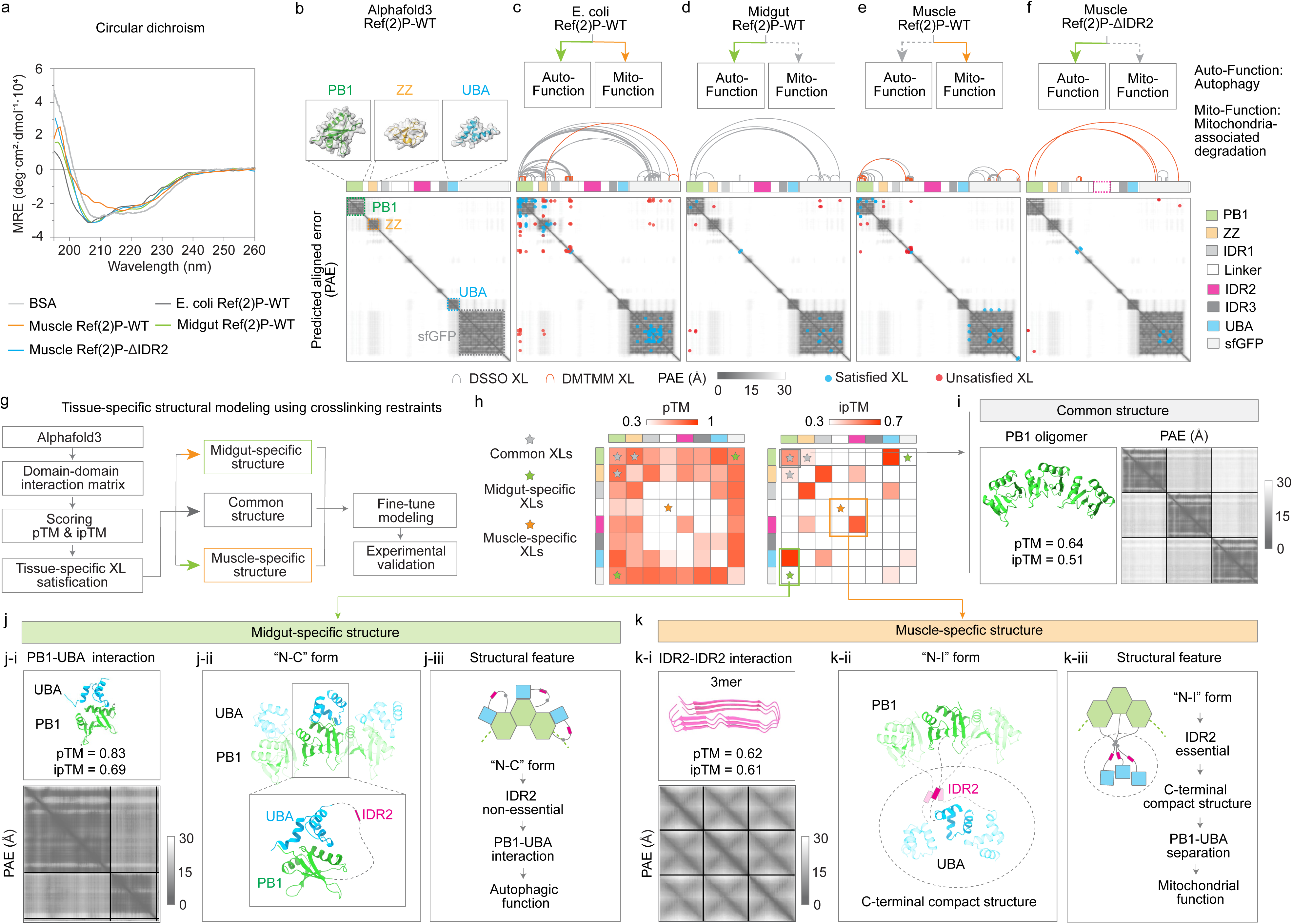
Tissue-specific structural states of Ref(2)P. **a**, Circular dichroism spectra (190-260 nm) of BSA, of Ref(2)P purified from *E. coli*, midgut and muscle, and of muscle Ref(2)P-ΔIDR2. **b**, AlphaFold3 prediction of Ref(2)P-sfGFP with the PB1, ZZ and UBA domains highlighted; predicted aligned error (PAE) scale 0-30 Å. **c-f**, Crosslinking patterns of Ref(2)P-sfGFP purified from *E. coli* (c), midgut (d) and muscle (e), and of muscle Ref(2)P-ΔIDR2 (f). Top, the activities retained by each protein (Auto-Function, autophagy; Mito- Function, mitochondria-associated degradation). Middle, crosslink arc diagrams: DSSO (grey) and DMTMM (orange). Bottom, PAE map annotated with the crosslinked pairs; satisfied crosslinks (blue) have a Cα-Cα Euclidean distance below 35 Å, unsatisfied crosslinks (red) above 35 Å. PAE scale 0-30 Å. **g**, Workflow for tissue-specific structural modelling using crosslinking restraints. **h**, Domain-domain interaction matrices from AlphaFold3 predictions (pTM scale 0.3-1.0; ipTM scale 0.3-0.7). Domains are indicated as the same colors as **c-f**; Stars mark the positions of crosslinked pairs: grey, common; green, midgut-specific; orange, muscle-specific. **i**, Common structural element of midgut and muscle Ref(2)P: the PB1 oligomer. PAE scale 0-30 Å. **j**, Midgut-specific structure. **j-i**, AlphaFold3 prediction of the PB1-UBA interaction. **j-ii**, The PB1-UBA interaction placed in a Ref(2)P trimer, defining the "N-C" form. **j-iii**, Structural features of midgut Ref(2)P: the "N-C" form, in which PB1 engages UBA and IDR2 is dispensable, supports autophagic function. **k**, Muscle-specific structure. **k-i**, AlphaFold3 prediction of the IDR2-IDR2 interaction (trimer). **k-ii**, The IDR2-IDR2 interaction placed in a Ref(2)P trimer, defining the "N-I" form and the resulting C- terminal compaction. **k-iii**, Structural features of muscle Ref(2)P: the "N-I" form, in which IDR2 is essential, compacts the C terminus and separates PB1 from UBA, supports mitochondrial function.

These observations suggested there are either tissue-specific factors (such as pH, ionic concentration, oxidative state, or mitochondrial abundance) that prevent Ref(2)P protein from binding to either autophagic factors or mitochondria, or there is something inherently different about the Ref(2)P protein itself in different tissues. To distinguish between these possibilities, we removed the tissue context and measured biochemical activity of purified Ref(2)P-sfGFP from five different sources: *E. coli*, S2 cells, midgut, oenocytes and muscle (Fig. 1f and Extended Data Fig. 4d-4f, see Methods). Purified protein was tested for binding to recombinant autophagic factor Atg8a, poly-ubiquitin (Fig. 1g) and mitochondria isolated from adult flies under the same biophysical condition, using similar amount of protein (Fig. 1h and Extended Data Fig. 4b-4c). The purified Ref(2)P protein preserved the specificity of tissue of origin. Midgut-derived Ref(2)P bound Atg8a far more efficiently than muscle-derived Ref(2)P, whereas muscle-derived Ref(2)P bound mitochondria more efficiently than midgut-derived Ref(2)P (Fig. 1g-1i). Ref(2)P purified from S2 cells behaved like midgut Ref(2)P, consistent with localization to autophagic structure in cell lines (Fig. 1a and Extended Data Fig. 1l). Oenocyte-derived Ref(2)P bound poly-ubiquitin but lowly to Atg8a (Fig. 1g), consistent with its localization to non-autophagic, ubiquitin-positive cytosolic granules. Notably, recombinant Ref(2)P purified from *E. coli* bound to both Atg8a and mitochondria (Fig. 1g-1i), indicating that the polypeptide is intrinsically capable producing both states. These biochemical differences are not an *in vitro* artifact substantially more Atg8a co-immunoprecipitated with Ref(2)P-sfGFP from midgut than from muscle or oenocytes (Extended Data Fig. 4a). Together, these results indicate that the same Ref(2)P protein localizes to different subcellular compartments due to some inherent difference.

## Ref(2)P mediates autophagic degradation in midgut and mitochondria-associated degradation in muscle

We sought to understand the functional significance of Ref(2)P’s differential localization, if any, and focused on two tissues, midgut and muscle, where Ref(2)P localizes to well characterized compartments, autophagosomes and mitochondria. (Fig. 2a,b). Several observations indicate that Ref(2)P performs autophagic function in gut, but not in muscle. Consistent with higher autophagic activity of gut, Atg8a-GFP-positive autophagic structures were readily detected under basal conditions (Extended Data Fig. 7c). In Ref(2)P-null mutants (Ref(2)P^-/-^) (Extended Data Fig. 5a-c) and tissue-specific knock down of Ref(2)P (Extended Data Fig. 5e-h), in midgut cells, ubiquitinated signal became dispersed instead of a typical punctate appearance (Fig. 2c,e). Loss of Ref(2)P also caused pronounced lipid-droplet accumulation only in the midgut (Extended Data Fig. 5d), a phenotype associated with defective lipophagy ^29^. We next examined whether stress altered the localization or function of midgut Ref(2)P. Neither starvation for two days, which activates autophagy ^30,31^, nor ageing (6 weeks old flies), which increases mitochondrial dysfunction ^32^, changed Ref(2)P localization (Fig. 2g and Extended Data Fig. 6). Thus, under both normal physiological conditions and stress, Ref(2)P localizes to autophagic structure in midgut, consistent with its known role in autophagic degradation in midgut ^17^.

What is the function of mitochondria localized Ref(2)P in muscle? Under normal physiological conditions in muscle, loss of Ref(2)P did not produce the dispersed-ubiquitin phenotype, unlike in the midgut (Fig. 2d,f). Under stress conditions, such as heat shock, mitochondria are known to participate in the degradation of both cytosolic and mitochondrial proteins ^33^. Following repeated heat shock (Extended Data Fig. 7d), ubiquitin and 26S proteasome signals accumulated on muscle mitochondria over 1-3 days of recovery (Fig. 2k,l and Extended Data Fig. 7f,g). In Ref(2)P-null muscle, accumulation of ubiquitin and proteasomal components on mitochondria was significantly reduced (From WT ∼ 42-fold to null mutant ∼1.8-fold, Fig. 2j-l), and the clearance of detergent-insoluble ubiquitinated proteins was also delayed (Fig. 2j-l and Extended Data Fig. 7e- g). This Ref(2)P-null defect had an organismal consequence, as only approximately 28% of Ref(2)P-null flies survived repeated heat shock, compared to ∼82% of control flies (Fig. 2m). Other physiological perturbations also produced distinct responses of Ref(2)P in muscle. Starvation did not alter the mitochondrial ubiquitin signal, whereas ageing increased ubiquitin accumulation on mitochondria (Fig. 2d,f). Nevertheless, in muscle of both starved and aged flies, Ref(2)P localized to mitochondria, unlike midgut where it localizes to autophagic structure (Fig. 2g and Extended Data Fig. 6).

Previous studies suggested two major processes mediates mitochondria-dependent ubiquitinated protein degradation: canonical mitophagy, which in most cases requires PINK1 ^34^, PARKIN ^35^ and autophagic flux ^36^, or mitochondria-associated protein degradation, which involves the mitochondrial import machinery ^33^, mitochondrial proteases ^37^ and/or cytosolic proteasome ^38^ (Fig. 2h and Extended Data Fig. 7a). We first compared the consequences of loss of PINK1 and PARKIN to that of loss of Ref(2)P in muscle. Neither loss of Ref(2)P nor the loss of PINK1 and PARKIN resulted in the accumulation of damaged, ubiquitinated mitochondria expected from defective mitophagy (Fig. 2i and Extended Data Fig. 7a,b). In contrast, depletion of the proteasome subunits Prosα6, Prosβ5 and Prosα7 markedly increased ubiquitin puncta in muscle (Fig. 2i and Extended Data Fig. 7b). Moreover, basal autophagic activity was nearly undetectable in muscle (Extended Data Fig. 7c). These results suggest in fly muscle Ref(2)P is involved in mitochondria-associated degradation rather than canonical mitophagy. Together, these findings suggest that Ref(2)P preferentially engages autophagic degradation in the midgut and mitochondria-associated proteostasis in muscle. The functions of Ref(2)P in muscle and gut are tissue-autonomous and likely linked to the tissue identity but not tissue state.

## Ref(2)P turnover is differentially regulated in the midgut and in muscle

Mammalian p62/SQSTM1 is both a regulator and substrate of autophagy ^17,18,39^. The differential function of *Drosophila* Ref(2)P in midgut and muscle prompted us to compare whether Ref(2)P itself is differentially regulated in these two tissues (Fig. 3a). Knockdown of autophagy genes involved in initiation of autophagy (ULK1/Atg1, Atg101) or at phagophore expansion (Atg5, Atg12) increased Ref(2)P in the midgut but not in muscle (Fig. 3b-3c and Extended Data Fig. 8a). Conversely, knockdown of the mitochondrial outer-membrane translocase Tom40 or of the inner-membrane translocases Tim23 and Tim9a increased Ref(2)P in muscle but not in the midgut (Fig. 3b-3c and Extended Data Fig. 8a). An increase in steady state Ref(2)P could arise from slower turnover or compensatory changes, such as transcriptional feedback. To this end, in addition to endogenous Ref(2)P, we also analyzed the level of GAL4/UAS-driven Ref(2)P-sfGFP transgene. The GAL4/UAS transgene-expression system lacks the endogenous regulatory sequences through which transcriptional feedback would normally act (Fig. 3d). The transgenic and endogenous proteins responded in the same tissue-restricted pattern: autophagy-gene knockdown increased the level of both endogenous and transgenic Ref(2)P in the midgut, and mitochondrial import- factor knockdown increased both in muscle (Fig. 3e-3g), suggesting the increase in Ref(2)P is likely due to reduced turnover.

Ref(2)P accumulation in fly muscle could be a secondary consequence of perturbation of mitochondrial import. If Ref(2)P is directly involved in a mitochondria-import dependent process, then in wild type condition, Ref(2)P should be present inside the mitochondria. Two independent lines of evidence suggest Ref(2)P translocates inside of the muscle mitochondria. First, we used a mitochondrial split-GFP reporter, where an inner mitochondrial membrane protein, cox-8, is fused to one half of GFP (COX8-mCherry-GFP1-10) and the other half of GFP is tagged to Ref(2)P (Ref(2)P-GFP11). We observed reconstitution of GFP fluorescence on mitochondria (Fig. 3h), suggesting Ref(2)P is present in the inner mitochondrial compartment. Second, we used TurboID ^40^ to biotinylate proteins within ∼10nm of Ref(2)P. We expressed Ref(2)P-fused TurboID in fly muscle and identified the biotinylated proteins by mass spectrometry (Fig. 3i and Extended Data Fig. 8c-8d). Of the 115 significantly enriched proteins, 31 were annotated as mitochondrial proteins (Fig. 3j). Out of these 31 proteins 17 proteins were inner-mitochondrial and matrix proteins (Fig. 3k). Taken together these observations suggest Ref(2)P itself is turned over by autophagy in midgut and mitochondria-import dependent degradation in muscle.

## Deletion of a single IDR switches the tissue-specific function of Ref(2)P

To begin to understand how the same Ref(2)P protein performs mutually exclusive tissue-specific functions, we asked which regions of the protein are required for its localization and functional differences. We generated transgenic flies expressing domain-deleted Ref(2)P variants and assessed tissue-specific ubiquitin- and Ref(2)P-positive puncta (Fig. 4a-4c and Extended Data Fig. 9a). Self-assembly of Ref(2)P, assessed from its punctate appearance, is important for its function ^41,42^. Deletion of the N-terminal PB1 domain prevented puncta formation in both midgut and muscle, whereas deletion of the C-terminal UBA domain prevented puncta formation only in muscle (Fig. 4b-4c and Extended Data Fig. 9a). Deletion of the ZZ domain, reported to bind RNA ^14^, did not change puncta formation but disrupted the localization of Ref(2)P to mitochondria in muscle (Extended Data Fig. 9a). Deletion of multiple domains, such as PB1 plus ZZ or PB1 plus UBA, abolished puncta formation in both tissues (Fig. 4b-4c). These observations indicate that the PB1 domain has a common role, whereas the UBA and ZZ domains have different roles in the self-assembly and localization of Ref(2)P in midgut and muscle.

In addition to its folded domains, Ref(2)P contains intrinsically disordered regions (IDRs). IDRs can serve multiple functions such as protein assembly, adopting a folded state upon binding a partner, or topological organization of already folded domains ^43–45^. Structure-prediction programs indicate that Ref(2)P has three IDRs: IDR1 (amino acids 192-225), IDR2 (360-449) and IDR3 (503-542) (Fig. 4a). We generated transgenic flies expressing Ref(2)P variants lacking one of the IDRs (Ref(2)PΔIDR) and examined puncta formation and localization (Fig. 4d). Deletion of any of the IDRs had no effect on Ref(2)P localization or function in the midgut (Extended Data Fig. 9b). In muscle, however, deletion of IDR1 or IDR2, but not IDR3, relocalized Ref(2)P to autophagic structures (Fig. 4d). To distinguish whether IDR deletion causes misfolding and consequent delivery to autophagic structures or genuinely switches the function of Ref(2)P towards autophagy, we tested the binding of the IDR-deleted variants to Atg8a. Although all three Ref(2)PΔIDR proteins were expressed equivalently in muscle, only deletion of IDR2 increased the Ref(2)P-Atg8a interaction (Fig. 4e), indicating that loss of IDR2 switches the intrinsic mitochondrial function of Ref(2)P to an autophagic one. Moreover, Ref(2)PΔIDR2 caused loss of muscle fibers (Fig. 4f and Extended Data Fig. 9c), indicating that forcing Ref(2)P from its mitochondrial to its autophagic function disrupts muscle homeostasis. Deletion of IDR1 also caused loss of muscle fibers, but without enhancing the Ref(2)P-Atg8a interaction (Fig. 4e and Extended Data Fig. 9c), consistent with likely misfolding and loss of Ref(2)P function rather than a switch in function. Together, these observations identify IDR2 as an important determinant of the tissue-specific localization and function of Ref(2)P (Fig. 4g).

## IDR2-dependent functional switch is not due to targeting, post-translational modification, or oligomeric state

How does IDR2 confer muscle-specific function? One possibility is that IDR2 itself contains a mitochondrial targeting signal. However, when expressed just by itself (Gal4:UAS-IDR2-sfGFP-Flag or Gal4:UAS-IDR2-cGFP), IDR2 did not localize to mitochondria (Extended Data Fig. 9d), and showed a diffuse signal in both muscle and gut (Extended Data Fig. 9e), suggesting IDR2 does not contain a mitochondria-targeting signal.

A second possibility is tissue-specific post-translational modification (PTM) within IDR2 or modification outside that work in conjunction with IDR2. Since the purified Ref(2)P retains tissue-specific function and recombinant Ref(2)P displays both functions, we searched for PTMs on purified Ref(2)Ps. We did not observe any difference in electrophoretic mobility of gut, muscle or recombinant Ref(2)P generally expected of PTMs (Fig. 1g-1h and Extended Data Fig. 4e-4f). To determine rare PTMs, if any, we analyzed purified Ref(2)P using mass spectrometry. Any causal PTM should be shared between the recombinant Ref(2)P and one of the tissue-derived Ref(2)Ps. Moreover, the interaction between Ref(2)P and Atg8 or mitochondria is stoichiometric (Fig. 1g- 1h) and therefore, any causal PTM should be present in substantial fraction of the protein. We first established that our mass spectrometry pipeline could detect low abundant PTMs (See methods). Under this experimental condition, no phosphorylated or acetylated peptides were detected from midgut Ref(2)P. For muscle Ref(2)P, only ∼0.1% of all detected peptides carried a known modification (Extended Data Fig. 10a-10d) and only one shared phosphorylation at S194 (in 0.07% of detected peptides) was detected between muscle and recombinant Ref(2)P (Extended Data Fig. 10a, 10c). Furthermore, gut Ref(2)P and muscle Ref(2)PΔIDR2 both have autophagic function, but only Ref(2)PΔIDR2 carried any detectable modifications (∼0.6% of detected peptides; Extended Data Fig. 10d). Therefore, the tissue-specific function is unlikely to arise from a stable and prevalent known post-translational modification.

A third possibility is difference in oligomeric state of Ref(2)P. Indeed "morpheeins" achieve distinct functions by forming distinct oligomers dictated by conformational change in the monomer ^46^. However, oligomeric state of purified Ref(2)P determined by size-exclusion chromatography (SEC) and negative-stain transmission electron microscopy (Extended Data Fig. 10e-10h) showed no obvious difference in Ref(2)P oligomeric state between tissues. We concluded that IDR2 is important for the tissue-specific functional difference for some other reason.

## Ref(2)P from muscle and midgut differs in both secondary and tertiary structure

The differential requirement for individual folded domains (Fig. 4a-4c and Extended Data Fig. 9a) and the ability of IDR2 to switch function prompted us to ask whether the functional difference reflects an inherent structural difference and/or a distinct topological organization of the folded domains. Limited proteolysis and thermal-shift assays have been used to assess such differences ^47,48^. Limited digestion of midgut and muscle Ref(2)P with papain, which cleaves after basic and aromatic residues, produced cleavage products of different sizes, indicating that the surface accessibility of these residues differs between the two tissues (Extended Data Fig. 11a-11b). Thermal-shift assays measure denaturation and consequent aggregation ^49^, behavior that is linked to the inherent structure of a protein. Both in isolation and in tissue lysate, midgut Ref(2)P aggregated at a lower temperature than muscle Ref(2)P (∼40 °C versus ∼55 °C; Extended Data Fig. 11c). These low-resolution assays suggested that Ref(2)P adopts different structures in muscle and gut. To obtain higher-resolution information we first attempted cryogenic electron microscopy. Beyond the PB1 domain, we could not obtain a high-resolution structure of rest of the protein, most likely because of the disordered regions; similar difficulties have been reported for human p62 ^19,50,51^. We therefore analyzed Ref(2)P from different tissues by circular dichroism, which reports secondary-structure content. The far-UV CD spectrum of midgut Ref(2)P showed more random coil and disorder, whereas muscle Ref(2)P showed higher α-helix and β-sheet content (Fig. 5a). Notably, muscle Ref(2)P-ΔIDR2, which behaves like midgut Ref(2)P, also had reduced α-helix and β-sheet content (Fig. 5a).

The difference in secondary structure raised the possibility of a difference in tertiary structure. We used crosslinking mass spectrometry (CL-MS) together with structural modelling to assess the difference in tertiary structure ^52–55^. Crosslinks report spatial distances between specific amino acids in three dimensions ^56,57^ and can be used as distance restraints to model interactions between domains and regions ^58^. We applied two crosslinkers to Ref(2)P-sfGFP isolated from midgut, muscle and bacteria (Fig. 5b): the MS-cleavable crosslinker disuccinimidyl sulfoxide (DSSO) ^59^, with a ∼10 Å spacer arm that crosslinks lysine pairs within a Cα-Cα distance of ∼35 Å; and the non-cleavable, zero-length crosslinker 4-(4,6-dimethoxy-1,3,5-triazin-2-yl)-4-methyl- morpholinium chloride (DMTMM), which crosslinks lysine to aspartate/glutamate within a Cα-Cα distance of ∼15 Å ^60^ (Extended Data Fig. 12a-12c). We found both common and tissue-specific crosslinked pairs. Midgut Ref(2)P showed unique crosslinks between the N-terminal PB1 domain and the C-terminal sfGFP (Fig. 5d), whereas muscle Ref(2)P had unique crosslinks between PB1 and the middle inter-domain region (Fig. 5e). We refer to the midgut pattern as the "N-C" form and the muscle pattern as the "N-I" form. Muscle Ref(2)PΔIDR2, which function like midgut Ref(2)P, also had a midgut specific "N-C" crosslinking pattern (Fig. 5f). Bacterially derived Ref(2)P, which retains both activities, showed crosslink pattern of both forms (Fig. 5c).

We next asked whether these unique crosslinks genuinely report inherent structural differences of Ref(2)P monomer. Because Ref(2)P forms homo-oligomers, a detected pair could join two residues of the same polypeptide (intra-chain) or two different polypeptides (inter-chain). To look for differences in Ref(2)P monomer (Extended Data Fig. 12e-12f, see Methods), we separated monomeric and oligomeric fractions of crosslinked samples by SDS-PAGE, excised the corresponding bands and performed in-gel digestion and mass spectrometry. In parallel we performed an in-solution reaction which allows monitoring of low-intensity crosslinked precursors (Extended Data Fig. 12g-12j). Twenty-two of the 42 unique pairs found in in-solution digests were also found in monomers isolated from gel (Extended Data Fig. 12g-12j). A few crosslink pairs were found in oligomeric fractions that were unique to muscle Ref(2)P (Extended Data Fig. 12i). We further confirmed that the unique crosslinks were not artefacts of insufficient precursor fragmentation, and for structural inference only considered reproducible crosslinks (Extended Data Fig. 12d, 12p). Importantly, the unique crosslinks assigned to the "N-C" form are intra-chain crosslinks reporting the structure of midgut Ref(2)P monomer. The muscle-specific "N-I" crosslinks are between equivalent residues within the same linker region (amino acids 244– 315, immediately before IDR2). Since a residue cannot be crosslinked to itself, such homotypic pairs report inter-molecular interaction in the "N-I" form (Extended Data Fig. 12k-12o). Taken together these results suggest that differential crosslink pattern between midgut and muscle Ref(2)P is indeed due to inherent structural differences in Ref(2)P monomer.

## Tissue-specific three-dimensional models of midgut and muscle Ref(2)P

To gain insight into the nature of the three-dimensional structural differences that contributed to the crosslink differences, we generated structural models of midgut and muscle Ref(2)P using the common and tissue-specific crosslinks (Fig. 5g). We first compared four modelling approaches based on crosslink satisfaction and prediction confidence (pTM and ipTM)and found an AlphaFold3-based workflow to be the most reliable (Extended Data Fig. 13a-13i, please see Methods for details). Only the folded domains of fly, mouse and human p62 were confidently predicted. The intervening regions remained unresolved (Extended Data Fig. 14a-14c). We then mapped the crosslink data onto the domain-domain interaction matrix and found that both midgut- and muscle-specific pairs fell on or adjacent to high-confidence interaction nodes (Fig. 5h and Extended Data Fig. 15a-15b) and focused on the regions within those nodes to refine the models (Fig. 5h).

Crosslink-guided modelling yielded three insights. First, a PB1 oligomer is a structural element common to midgut and muscle Ref(2)P (Fig. 5i and Extended Data Fig. 14d-14f). Second, a PB1- UBA interaction is specific to midgut Ref(2)P (Fig. 5j and Extended Data Fig. 15c-15e). Third, in muscle the crosslink between PB1 and the middle region is best explained by IDR2 forming an intermolecular assembly that compacts the C-terminal half of the protein (Fig. 5k and Extended Data Fig. 15f-15h).

## Experimental validation of the tissue-specific structural states

To test the validity of the predicted structural models we tested the key elements of the models: the common PB1-PB1 oligomer, the midgut-specific PB1-UBA interaction, and the muscle-specific IDR2 assembly (Extended Data Fig. 16a). Based on the predicted model (Fig. 5i and Extended Data Fig. 14d-14e), we mutated residues that are in the interface of PB1-oligomer (K7A, D58A, D60A and D62A; Fig. 6a). Double K7A, D60A mutation reduced Ref(2)P self-assembly similar to PB1 domain deletion mutant (Fig. 6b and Extended Data Fig. 16b-16e), and fluorescence recovery after photobleaching showed that the mutants formed more labile assemblies compared to wild type Ref(2)P (Extended Data Fig. 16f).

**Fig. 6:**
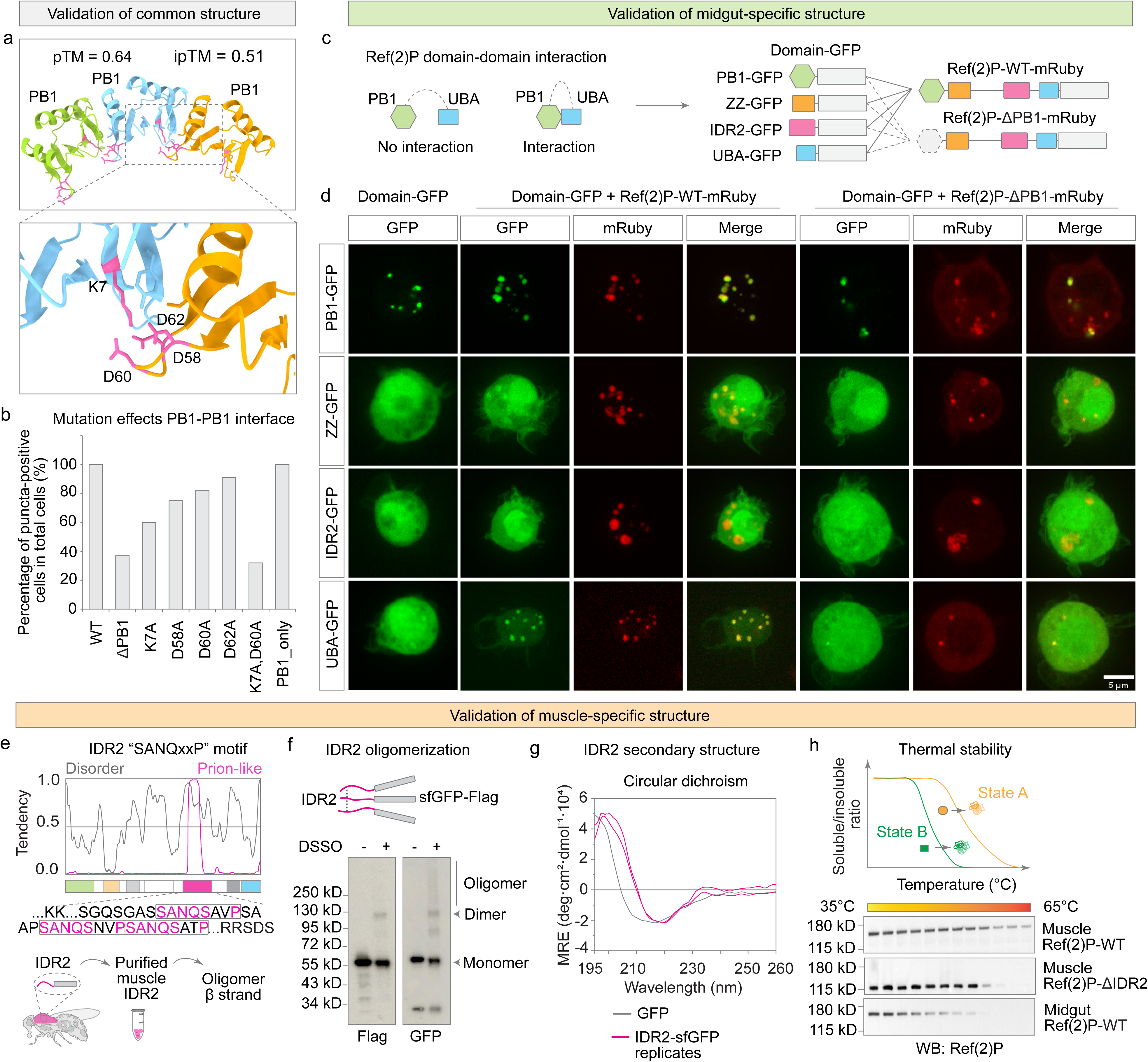
Experimental validation of the tissue-specific structural states of Ref(2)P. **a**, Validation of the common structure. AlphaFold3 model of the PB1 trimer (pTM 0.64, ipTM 0.51), with the predicted interface enlarged below: K7 of one chain contacts D58, D60 and D62 of the neighboring chain. **b**, Percentage of puncta-positive cells among all fluorescent S2 cells expressing wild-type Ref(2)P, Ref(2)P-ΔPB1, the indicated PB1 interface mutants, or the PB1 domain alone. Representative images, the mutant PB1 trimer model and FRAP analysis are in Extended Data Fig. 16b-16f. **c**, Design of the domain-trapping assay used to test the predicted intramolecular interactions in S2 cells, where Ref(2)P behaves as it does in the midgut. A domain- GFP fusion is co-expressed with Ref(2)P-WT-mRuby or Ref(2)P-ΔPB1-mRuby; recruitment of the domain into Ref(2)P-WT puncta but not into Ref(2)P-ΔPB1 puncta indicates an interaction with PB1. **d**, Fluorescence images of S2 cells expressing PB1-GFP, ZZ-GFP, IDR2-GFP or UBA-GFP alone (left) or together with Ref(2)P-WT-mRuby (middle) or Ref(2)P-ΔPB1-mRuby (right). ZZ and UBA, but not IDR2, are recruited into Ref(2)P-WT puncta. Scale bar, 5 µm. **e**, Validation of the muscle- specific structure. Top, disorder (grey) and prion-like (magenta) tendency of Ref(2)P, identifying three copies of a "SANQxxP" motif within IDR2; the motif sequences are shown below. Bottom, scheme for purifying the IDR2 fragment from muscle to test self-assembly. **f**, DSSO crosslinking of IDR2-sfGFP-Flag purified from muscle, analyzed by western blotting for Flag and GFP. Expression and purification are shown in Extended Data Fig. 16g-16h. **g**, Circular dichroism spectra of GFP and of IDR2-sfGFP purified from muscle. **h**, Thermal stability of Ref(2)P. Top, schematic of the assay: the soluble/insoluble ratio is followed across a temperature gradient, with state B less stable than state A. Bottom, western blots of muscle Ref(2)P-WT, muscle Ref(2)P-ΔIDR2 and midgut Ref(2)P-WT across a 35-65 °C gradient. Deletion of IDR2 converts muscle Ref(2)P to the lower-stability behavior of midgut Ref(2)P.

To assess the predicted PB1-UBA interaction, we used S2 cells, since in S2 cells and midgut Ref(2)P is involved in autophagy (Fig. 1a, 1i and Extended Data Fig. 1l). Individual domains fused to GFP were co-expressed with either Ref(2)P-WT-mRuby or Ref(2)P-ΔPB1-mRuby; recruitment into Ref(2)P-WT puncta but not into Ref(2)P-ΔPB1 puncta indicates a PB1-dependent interaction (Fig. 6c). Expressed alone, neither ZZ-GFP nor UBA-GFP formed puncta. Co-expressed with Ref(2)P- WT, but not with Ref(2)P-ΔPB1, both were recruited into puncta, indicating that ZZ and UBA either contact PB1 directly or join existing PB1-dependent assemblies (Fig. 6d). IDR2-GFP, in contrast, was not recruited by either Ref(2)P-WT or Ref(2)P-ΔPB1 (Fig. 6d), even though both proteins contain IDR2, implying that this interaction requires the muscle environment.

We then tested the muscle-specific prediction directly. The IDR2 domain contains three copies of a "SANQxxP" motif that can form cross-β rich self-assembly (Fig. 6e). To test whether IDR2 is indeed self-assembled in muscle, we expressed IDR2-sfGFP driven by muscle GAL4 and purified the IDR2-sfGFP fragment from muscle (Extended Data Fig. 16g-16h). Crosslinking with DSSO revealed dimer- and oligomers of IDR2-sfGFP (Fig. 6f and Extended Data Fig. 16i). Circular dichroism of muscle-purified IDR2-sfGFP showed a spectrum consistent with β-sheet (Fig. 6g). Finally, cross-β rich self-assembly confers stability. If IDR2 indeed from cross-β rich self-assembly in muscle, then its removal should reduce muscle Ref(2)P stability. Indeed, muscle Ref(2)P-ΔIDR2 showed lower thermal stability (Muscle Ref(2)P-WT: >65 °C; Muscle Ref(2)P-ΔIDR2 ∼ 54 °C; and Midgut Ref(2)P-WT: ∼ 45 °C) and lower β-sheet content similar to midgut Ref(2)P (Fig. 5a, Fig. 6h).

These observations suggest in midgut Ref(2)P "N-C" form PB1 interacts with UBA and IDR2 is dispensable. Whereas muscle Ref(2)P adopts the "N-I" form, in which IDR2 is essential and forms a β-sheet enriched homo-assembly (Fig. 5j-iii, 5k-iii and Extended Data Fig. 16a). Can these structural differences account for the functional differences between midgut and muscle Ref(2)P (Fig. 1g, 1i and Fig. 7a). In mammals, p62 binds to LC3, mammalian Atg8a orthologue, through the LC3-interacting region or LIR (or AIM in *Drosophila*), which is comprised of an aromatic residue followed by two variable residues and an aliphatic residue ^15,61^. In Ref(2)P, a putative Atg8a interacting motif (AIM), W454-x-x-I457, is located immediately C-terminal to IDR2 (Fig. 7b). An AlphaFold3 model of the Atg8a–AIM complex superimposes on the human LC3–LIR structure with an interface RMSD of 0.89 Å (Fig. 7c and Extended Data Fig. 17a-17c), indicating that Ref(2)P W454-x-x-I457 is a bona fide AIM. The location of Ref(2)P AIM suggests in muscle, where IDR2 oligomerizes, the AIM should be sequestered within the compacted C-terminal region and thereby interfere with Atg8a. Indeed, when *in vitro* translated Ref(2)P is incubated with recombinant Atg8a, in presence or absence of IDR2 purified from muscle (Fig. 7e), muscle IDR2 significantly reduced Ref(2)P-Atg8a interaction (Fig. 7f-7g). Taken together these results suggest that in muscle, IDR2-oligomerization and compaction of c-terminal end occlude the Atg8- interacting motif and prevent localization to autophagy.

**Fig. 7:**
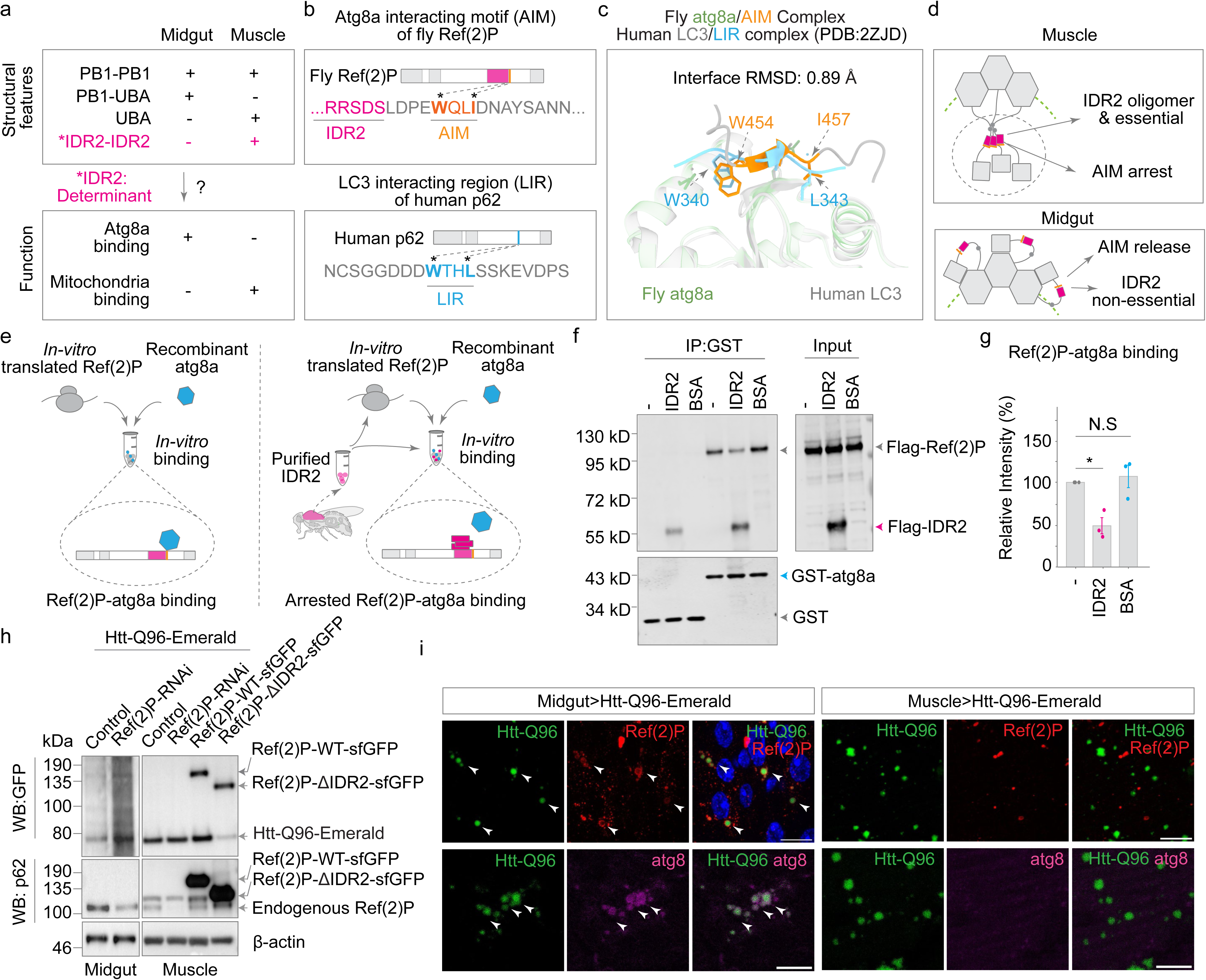
IDR2-dependent structural organization directs Ref(2)P-mediated protein clearance. **a**, Summary of the structural features and activities assigned to midgut and muscle Ref(2)P, identifying the IDR2-IDR2 interaction as the determinant to be explained. **b**, The Atg8a- interacting motif (AIM) of *Drosophila* Ref(2)P (W454-x-x-I457) lies immediately C-terminal to IDR2, in the position equivalent to the LC3-interacting region (LIR) of human p62. Asterisks mark the two anchor residues. **c**, Superposition of the AlphaFold3 model of the fly Atg8a-AIM complex on the human LC3-LIR complex (PDB 2ZJD). Interface RMSD 0.89 Å; the W and I/L anchor residues are labelled. The full comparison is in Extended Data Fig. 17a-17c. **d**, Model arising from a-c. In muscle, IDR2 oligomerizes and is essential for the structure, sequestering the adjacent AIM (AIM arrest); in the midgut, IDR2 is dispensable and the AIM is available for Atg8a binding (AIM release). **e**, Design of the *in vitro* competition experiment. In vitro-translated Ref(2)P was incubated with recombinant Atg8a in the absence (left) or presence (right) of IDR2 purified from muscle. **f**, GST pull-down from e. GST-Atg8a or GST alone was used for capture; anti-Flag detects Ref(2)P and the IDR2 fragment. Input, 2%. **g**, Quantification of Ref(2)P(2)P-Atg8a binding from f. Mean ± s.d. of n = 3 independent experiments; two-tailed unpaired t-test; *P < 0.05. Added IDR2 reduces Ref(2)P-Atg8a binding. **h**, Western blot of tissue lysates from flies co-expressing Htt-Q96- Emerald with Ref(2)P RNAi, with Ref(2)P-WT-sfGFP or with Ref(2)P-ΔIDR2-sfGFP, in midgut (left) and muscle (right). Anti-GFP detects both Htt-Q96-Emerald and the Ref(2)P transgenes; anti- Ref(2)P detects endogenous and transgenic Ref(2)P; β-actin, loading control. **i**, Immunofluorescence of midgut (left) and muscle (right) expressing Htt-Q96-Emerald (green) stained with anti-Ref(2)P (red, top) or anti-Atg8a (magenta, bottom). Arrowheads mark colocalization. Scale bars 10 µm

## Tissue-specific function of Ref(2)P protein results in tissue-specific degradation of aggregation- prone proteins

What are the biological consequences, if any, of tissue-specific deployment of autophagy and mitochondria-associated protein degradation? Many cytosolic aggregation-prone proteins, including Huntingtin (Htt) polyQ proteins ^62,63^, can be degraded through different quality-control pathways depending on the cellular context ^33,18,64,65^. We therefore expressed aggregation-prone Htt-Q96 in muscle or midgut. Htt-Q96 formed aggregates in both tissues but colocalized with Ref(2)P and Atg8a only in the midgut (Fig. 7i). Consistent with this localization, Ref(2)P knockdown increased Htt-Q96 protein approximately threefold in the midgut but had no detectable effect in muscle (Fig. 7h, first four lanes), indicating that Ref(2)P-dependent autophagy mediates Htt-Q96 clearance in the midgut. Expression of Ref(2)PΔIDR2, which promotes autophagic engagement in muscle also reduced Htt-Q96 protein approximately fivefold in muscle (Fig. 7h, last two lanes).

Is this specific to polyQ, or does it reflect a broader phenomenon? Ectopic expression of aggregation-prone amyloid-beta 42 (Aβ*42*) or of a variant of human superoxide dismutase (hSOD1-A4V) caused a marked accumulation of Ref(2)P aggregates in muscle — approximately 17-fold and 8-fold, respectively — but not in the midgut (Extended Data Fig. 17d, 17f). Conversely, poly-GA promoted Ref(2)P aggregation in both muscle (∼6.5-fold) and gut (∼5-fold) (Extended Data Fig. 17d, 17f), and a graded polyQ series (Htt-Q25 to Htt-Q96) engaged Ref(2)P in muscle at lower repeat lengths than in the midgut (Extended Data Fig. 17e-17f). Expression of *Aβ42* and of poly-GR in muscle was also more detrimental to the organism than expression in the gut (Extended Data Fig. 17d). Taken together, these data indicate that protein aggregation can have distinct fates in different tissues, and that this difference is due in part to the tissue-specific, Ref(2)P-dependent engagement of different protein quality control systems.

## Discussion

Although all cells in an organism share the same protein-coding genes, proteome content and function diverge across cell types. The cell type specific proteome emerges from quantitative (how much of a protein is produced) or qualitative differences (which isoform or modified form is made due to alternative splicing/editing, non-AUG codon ^66^ or reading through stop codons ^67,68^ or post-translational modifications). Here, we report an unexplored mechanism that does not rely on changes in protein amount or amino acid sequence. We find that *Drosophila* p62, Ref(2)P performs two mutually exclusive, tissue-autonomous functions because the same polypeptide adopts distinct, tissue-specific three-dimensional structures. What causes Ref(2)P to adopt the tissue-specific structure and ensuing function remains unclear. Either the physiological differences between muscle and midgut lead to the differential role of Ref(2)P or the differences in the Ref(2)P function contribute to the physiological differences.

## Protein pleomorphism as a plausible mechanism of tissue-specificity of protein multifunctionality

The possibility that some proteins deviate from one of the central tenets of biology “one polypeptide-one structure” is neither new nor unprecedented ^69,70^. There are documented examples of proteins that switch between stable folded states with different secondary structures, distinct core architectures, or newly exposed surfaces ^4,71,72,73^. We have recently observed that adaptation of yeast to different environmental conditions results in same polypeptide attaining distinct structural and functional state in context-specific manner ^74^. The behavior of Ref(2)P is reminiscent of another protein mitotic arrest deficiency 2 (Mad2) ^75,76^. In Mad2, part of protein remains unchanged, whereas 60-200 amino-acid segment undergoes large structural change to create closed (active) and open (inactive) states ^76^. In Ref(2)P, the N-terminal PB1 likely remains similar between muscle and gut, but the topological organization between the N-terminal PB1 domain and C-terminal UBA domain, as well as secondary structure of intervening regions, differs significantly. In muscle, the IDR2 cross-β structure positions the C-terminal domains away from the N-terminal PB1. This rearrangement likely sequesters the binding interface or motifs for autophagy factors, such as Atg8a-interacting motifs (AIMs), and thereby abolishing the autophagy function.

While the properties of Ref(2)P are reminiscent of other instances of “one polypeptide-many structures” such as metamorphic proteins ^77,78^ and fold-switching proteins ^4^, they differ from these proteins in specific ways. First, protein metamorphosis and fold-switching allow a protein to adopt multiple reversible or irreversible structural states within the same cell, while Ref(2)P requires different tissue environments to adopt multiple, mutually exclusive, structural states. Second, while metamorphic and fold-switching proteins typically switch between conformations in response to specific stimuli to achieve functional changes ^71,72^, switching among Ref(2)P conformations leads to tissue dysfunction. This suggests that the existence of multiple structural states of Ref(2)P protein has evolved as an inherent mechanism to ensure tissue-specific physiology.

We coined the term “pleomorphic protein” to capture the phenomenon that broadly expressed multifunctional proteins can utilize tissue- or cell type–specific environments to adopt a particular structural state and execute one function among multiple potential functions. Recent advances in proteomics have revealed that 3-5 % of the human proteome is highly multifunctional ^79–81^, with many broadly expressed proteins forming distinct protein complexes in different contexts. Protein pleomorphism represents a mechanism to create multiple functions from a single protein without altering its amino acid sequence. In the future, it will be important to determine the breadth of protein pleomorphism and how a pleomorphic protein adopts tissue-specific structural states. To date, about a hundred proteins have been classified as fold- switching proteins based on Protein Data Bank (PDB) structures ^4,82^ and AlphaFold2 prediction ^83^. However, for only a few, such as KaiB in cyanobacterium ^72,84^, RfaH in *E. coli* ^71^ and the chemokine lymphotactin in humans ^73^, has it been well documented that an alternate structure confers new functions. Whether metamorphic and fold-switching proteins also show tissue selectivity in deploying multiple structures and functions remains an intriguing question for future research.

Our finding also suggests tissue-specific proteostasis can be shaped by structural specialization of a shared proteostatic regulator, in addition to tissue-restricted expression of proteostasis regulators ^85^. It is documented that the basal activities of major degradation pathways differ across tissues ^86–88^. However, the extent of these differences and underlying principles governing biased usage of various pathways remains poorly defined. An important next step is to test whether analogous pathway bias is conserved in mammals and whether it contributes to tissue- specific proteome composition and vulnerability.

A limitation of this study is the resolution of our structural evidence. While our data clearly supports that Ref(2)P adopts distinct structural states in midgut versus muscle—reflected in differences in secondary-structure content and overall tertiary organization, we do not yet have atomic level understanding of the structural differences between tissues. This limitation partly rooted in a technical challenge of structural analysis of small proteins with intrinsically disordered region. A second limitation is we do not know how Ref(2)P engages mitochondria. Our genetic, reporter and proximity-labelling data suggest a relationship between Ref(2)P, its cargo and the mitochondrial import machinery. However, neither demonstrates translocation through TOM and TIM. We therefore cannot yet distinguish whether Ref(2)P is imported and degraded within the organelle or is held at the surface while its cargo is translocated. IDR2 is necessary for mitochondrial association but is not itself a targeting sequence, so both the surface created by the muscle-specific state and the mitochondrial component it recognizes remain to be identified.

## Acknowledgement

We thank A. Sánchez Alvarado, R. Krumlauf, J. Workman, and L. Li for their constructive comments on the manuscript. We thank G. Smith for proof reading. We thank J. Chen for in-gel digestion and crosslinking mass spectrometry. We thank B. Redwine for providing GFP nanobody plasmid. This work was supported by grants from Stowers Institute for Medical Research to K. Si.

## Contributions

A. Y. Yan, K. Zhang and K. Si conceptualized the project. Y. Yan and K. Zhang were responsible for most data acquisition, analysis, and visualization; Y. Zhang helped in acquisition of mass spectrometry data, K. Yi performed the electronic microscopy experiments, P. Leal helped generating transgenic lines. G. Pycior provided technical support in biochemical experiments; Z. Yu performed CLEM experiments and supported imaging analysis; P. Guillen-Poza and R. Hervas helped in protein purification and cryoEM; L. Florens helped in mass spectrometry data analysis; J. Unruh helped developing code for structural modeling. Y. Yan and K. Zhang initiated the manuscript draft, and K. Si and Y. Yan revised the manuscript; K. Si supervised the work and acquired funding.

## Methods

### Drosophila stocks

We obtained Act88F-GAL4 (#38459 and #38461), cad-GAL4 (#3042), Desat1-GAL4 (#65405 and #65406), elav-GAL4 (#458), D42-GAL4 (#8816), UAS-Luc (#35788), UAS-mCherry^RNAi^ (#35785) UAS-Ref(2)P^RNAi^ (#36111 and #33978), UAS-ULK^RNAi^ (#26731), UAS-atg5^RNAi^ (#27551), UAS-atg12^RNAi^ (#27552), UAS-atg101^RNAi^ (#34360), UAS-tim9a^RNAi^ (#55235), UAS-tim23^RNAi^ (#77421), UAS-tom40^RNAi^ (#26005), UAS-pink1^RNAi^ (#31170) and UAS-park^RNAi^ (#37509), UAS-hSOD.A4V (#33607), UAS-Aβ42 (64216), UAS-polyGR100 (#58696), UAS-polyGA100 (#58697), UAS-htt-Q25 (#58360), UAS-htt-Q46 (#56777), UAS-htt-Q72 (#58361), UAS-htt-Q96 (#56774) lines from Bloomington Drosophila Stock Center at Indiana University. The following stocks were generated in this study: Ref(2)P^-/-^ knockout, Ref(2)P-Flag-knock-in, Ref(2)P-APEX2-Flag-knock-in, UAS- Ref(2)P-WT-sfGFP, UAS-Ref(2)P△PB1-sfGFP, UAS-Ref(2)P△ZZ-sfGFP, UAS-Ref(2)P△LIR-sfGFP, UAS-Ref(2)P△UBA-sfGFP, UAS-Ref(2)P△(PB1+ZZ)-sfGFP, UAS-Ref(2)P△(PB1+UBA)-sfGFP, UAS- Ref(2)P△IDR1-sfGFP, UAS-Ref(2)P△IDR2-sfGFP, UAS-Ref(2)P△IDR3-sfGFP, UAS-IDR2-sfGFP-Flag, UAS-Ref(2)P-TurboID, UAS-Ref(2)P-GFP11, UAS-NGFP-Ref(2)P-CGFP, UAS-Cox8-GFP1-10 and UAS-LucDM. All flies were raised at 25 °C on standard medium on a 12-hour light/dark cycle unless otherwise indicated. For oenocyte expression (Driven by Desat1-GAL4 & tubP-GAL80[ts] (#65406)), flies were developed at 18 °C and further induced at 29 °C for 3-days before experiments. All genetic experiments were conducted with female animals. A complete list of fly genotypes used in this study is presented in the Supplementary Table. Transgenic flies were generated in house or using the transgenic service of the company Bestgene.

### CRISPR–Cas9-mediated genomic editing

To generate Ref(2)P null mutant, the gRNAs (gRNA1 sequence “AAAATCACCTACCAGGGCGC” and gRNA2 sequence “TATTGCGTCGCGAAATCGAG”) were cloned into pU6.2 and pU6.2B vectors (Addgene) respectively. The U6-gRNA from U6.2 was excised with EcoRI/NotI and cloned into the same sites of pU6.2B to create a single vector expressing both gRNAs. The gRNAs were injected into *nos-cas9* (NIG-FLY#CAS0001) flies. Mutants were screened and confirmed by PCR and sequencing.

To generate Ref(2)P-Flag-knock-in fly, gRNA3 (sequence “GCTAACCCAGCTCCTAGAGT”) and gRNA4 (sequence “GCGATACGTTCATTACGTCCA”) were cloned in pU6.2B as described above. pHD-Scareless-DsRed donor vector was used as template vector. Left arm was amplified from fly genome with primer “AGACCTATAGTGTCTTCGGGGCCGAAAAGTGTCCCTAGAACCGAAGATCC” and “GTAATGAACGTATCGCAGAACCGCAAC;” right arm was amplified with primer

“CGCAGACTATCTTTCTAGGGTTAAgATATTCATATTCAATATCGA” and “GAGTTCGTAATTTGCAGTCGCCGGGAAAAGAAGACCATATACGTC.” TEV-3XFlag cDNA were synthesized and cloned into pHD-Scareless-DsRed donor vector. Both gRNA and donor vector were injected into *vas-cas9*/Transposase flies. DsRed was used for initial screening. Once successful insertion was verified by sequencing, DsRed was flipped out with transposase. Homozygous stocks were established by crossed with flies harboring balancer chromosomes.

Knock-in files were screened and confirmed by PCR and sequencing.

### Molecular cloning

#### Plasmids for cell lines

Ref(2)P coding sequences (CDS) was amplified from *Canton-S* fly cDNA with primer 5’- ATGCCGGAGAAGCTGTTG-3’ and 5’-TTAGTTGCGGTTCTGCGATACG-3’. Atg8a CDS was amplified from *Canton-S* fly cDNA with primer 5’-ATGAAGTTCCAATACAAGGAGG-3’ and 5’- TTAATTTTGGCCATGCC-3’. Ref(2)P and Atg8a CDS were cloned into the pEntr/D-Topo vector (Invitrogen, K2400-20) and then into pAWG (#1072), pAWF (#1112) or pAWR (#1132) expression vectors (Drosophila Genomics Resource Center) with the Gateway LR Clonase II kit (Invitrogen, 11791020) to generate plasmids for S2R+ and KC167 cells. sfGFP sequence was further amplified by the primer pair of 5’- GGCGCGCCATGGTGTCCAAGGGCGAGGAGCTGTT-3’ and 5’- GAGCTCCCCTTGTACAGCTCATCCATGCC -3’ and cloned as an insert into the Flag-tagged Ref(2)P plasmid backbone (pAWF-Ref(2)P) with two restriction enzymes, AscI (NEB, R0558S) and SacI-HF (NEB, R3156S) to generate C-terminal sfGFP and Flag tagged Ref(2)P-WT and truncated variant plasmids (pAWGF-Ref(2)P).

### Plasmids for transgenic fly lines

The Ref(2)P-sfGFP-Flag sequence was amplified from the pAWGF-Ref(2)P plasmid using 5’- TGGGAATTCATGCCGGAGAAGCTGTTGAA-3’ and 5’-ACCCTCGAGTCACGTGGACCGGTGCTTGT-3’ primer pairs and cloned into the EcoRI-XhoI site of pUAST-attB plasmid (DGRC, #1419) to generate pUAST-Ref(2)P-sfGFP-Flag plasmid. Domain deletion variants of Ref(2)P were generated by QuikChange II Site-Directed Mutagenesis Kit (Agilent, 200523) with the following primers:

△PB1: 5’-ctacgggcgccagtggcatggtgaagggggcg-3’ and 5’-cgcccccttcaccatgccactggcgcccgtag-3’
△ZZ: 5’-attgttggtcggcatgcgtagggtgaaattgctcggatcgt-3’ and 5’-ctacgcatgccgaccaacaat-3’
△LIR: 5’-ctctggatccaagctgtctgaacg-3’ and 5’- tcagacagcttggatccagaggacaatgcatactctgcaaacaatagtaac-3’
△UBA: 5’-cccacccttgttgcggtagacggggaccgt-3’ and 5’-acggtccccgtctaccgcaacaagggtggg-3’
△IDR1: 5’-actgagtcaagactccggcatggccactgcggcgtcccaat-3’ and 5’-catgccggagtcttgactcagt-3’
△IDR2: 5’-gtcaataagctgccactctggatcaaaaattcccgaatcaccgc-3’ and 5’-gatccagagtggcagcttattgac-3’
△IDR3: 5’-agcctcctcattcatgtgctg and 5’-cagcacatgaatgaggaggctaagcgcacggtccccgtcta-3’

The LucDM is a luciferase variant that harbors mutations R188Q and R261Q ^89^. These mutations make the protein prone to aggregation. LucDM sequence was amplified from the FlucDM-HA- GFP11-N1 plasmid (addgene, #91956) by the primer pair of 5’- ATCTGCGGCCGCGGCTCGAGATGGAAGACGCCAAAAACAT-3’ and 5’- AAGATCCTCTAGAGG- TACCTTATGCGTAGTCGGGCACGTCGTAGGGGTAGCTACCAGCTGCGTAGTCGGGCACGTCGTAGGGG TAGCTACCAGCCACGGCAATCTTTCCGCC-3’ and cloned as an insert into XhoI-KpnI site of pUAST- attB plasmid (DGRC, #1419) to generate pUAST-LucDM-2xHA plasmid.

The pUAST-LucDM-2xHA, pUAST-Ref(2)P were inserted into a specific genomic location using attp-attB and PhiC31 integrase-mediated transgenesis systems (Bestgene). W+ marker (Red eye color) was used to screen transgenic flies.

### Plasmids for bacterial protein expression

9XUbiquitin coding sequence was synthesized from IDT (www.idtdna.com) and cloned into pET- His6-MBP-PreScission vector with primer 5’- TGTCAAGTGACGATCACAGCGATTCGGACCCTGGAACAGCACTTCCAGCTC-3’ and 5’- GGCAATCAAATCTTCCGAATCTACTTAAAAGGGGAAACTCCGAC-3’ by Gibson assembly. Atg8a coding sequence were amplified from pAWR-Atg8a vector and cloned into pET-His6-MBP- PreScission vector with primer 5’- CGAGCTGGAAGTGCTGTTCCAGGGTCCGAATATGAAGTTCCAATACAAGGAGGAGCACGCC-3’ and 5’- GGCAATCAAATCATTCCGAATCTAACCCTTGTTAATTTTGGCCATGCCG-3’ by Gibson assembly. For pET-His6-MBP-Ref(2)P-sfGFP vector, sfGFP was first cloned into pET-His6-MBP-PreScission vector with primer 5’- GTGCTGTTCCAGGGTCCGAATCTCGAGGAATTCGGTAAGCTTGAAAACCTGTATTTTCAGGGTGAGCTT ATGGTGTCCAAGGGCGAG-3’ and 5’-GGCAATCAAATCATTCCGAATCTACTTGTACAGCTCATCCATGCC-3’ by Gibson assembly and then Ref(2)P CDS was inserted with restriction enzymes EcoRI and HindIII. GFP nanobody sequence ^90^ was synthesized by IDT (www.idtdna.com) and cloned into pET26b vector with two restriction enzymes NdeI and BlpI.

### Plasmids for S2 cell expression

pAWG vector from DGRC (DGRC_1072 ) under Act5c promoter.

The following primers were used:

PB1-EGFP: 5’-ggtactcgagATGCCGGAGAAGCTGTTGAA-3’ and 3’-

acggagctcacTGGAGCCACCTGAACGTGCAT-5’

ZZ-EGFP: 5’- ggtactcgagatgATCCACGACGCCGTTGAA-3’ and 5’- acggagctcacCATCAAGTGCTCAGGATGC-3’

IDR2-EGFP: 5’-ggtactcgagatgGATTCGGGAATTTTTGCGCC-3’ and 3’-

acggagctcacTGGCTTAGGATTCAGGGGCTCT-5’. Construct has a Proline to Leucine mutation at CDS a.a. 40 (position 395 in Ref(2)p CDS

UBA-EGFP: 5’-ggtactcgagatgCACACTGATGAAAGCATCAA-3’ and 3’-

acggagctcacGTTGCGGTTCTGCGATACGT-5’ XhoI and SacI cut sites were used to insert CDS.

### Cell culture

S2R+ (#150) cell line was originally obtained from Drosophila Genomics Resource Center and maintained by Tissue Culture Core of Stowers Institute of Medical Research. S2R+ cells were cultured in Schneider’s Drosophila Medium (Thermo Fisher, 21720024) with 5% Fetal Bovine Serum (Thermo Fisher, 16140071) and 1% antibiotics penicillin and streptomycin (Thermo Fisher,15140122) in a 25 °C tissue culture incubator. Expression vectors were transfected into S2R+ cells with Effectene transfection reagent (Qiagen, 301427).

### Fluorescence Recovery After Photobleaching (FRAP)

S2R+ (#150) cell line was transfected with different variants of Ref(2)P-sfGFP plasmids in a confocal glass-bottom dish (ASI, TGC22) one day before FRAP. The Nikon spinning disk system was used for the data acquisition, with a bleach laser (488 nm) setting at 100% AOTF, 10 pre-bleach frames with 500 ms interval and 360 post-bleach frames with 500 ms interval collected. Image analysis was completed in ImageJ. Briefly, after background subtraction, the intensity in the bleached region was plotted and fit with the following equation: I(t) = A(1-exp[-t/tau])+B. Here, tau is the recovery time constant, A is the recovery amplitude, and B is the background intensity. Fraction mobile was calculated using fmobile = A/(IpreFRAP-B), where IpreFRAP is the average intensity in the region of interest prior to its bleaching. All data plotted in each condition was the average recovery percentage of N>=9 puncta.

### Heat-shock assay

Three-day old 20 female flies were collected into empty vial and heat shocked for 30 min in a 38.5 °C water bath. Following heat shock flies were transferred back into vials with standard fly food at 25 °C. Tissues were dissected for immunofluorescence staining and western blot analysis at different time points after heat shock. To measure heat shock resistance, both wildtype and Ref(2)P-mutant flies were heat shocked as mentioned above for three times, every 24-hr and the number of live flies were counted after 24 hr recovery from the last heat shock. Each group had 3 independent vials.

### Starvation assay

Three day old 20 female flies were transferred into a vial with wet paper and housed at 25 °C under 12-hour light/dark cycle and 60 % humidity. Wet paper was changed every 24 hours to ensure hydration. After 48hours food deprivation, flies were dissected for immunostaining and western blot analysis. Each group consists of 3 independent vials.

### Lethality assay

Ten age-matched males and virgin females were crossed for each experimental condition. Genotypes of F1 generation were continuously monitored. To assess fitness, we calculated the ratio of observed hatch rates to that of expected hatch rates for each genotype within a cross. This procedure was replicated across three independent crosses per group for quantification.

### Immunofluorescence and imaging analyses

Fly tissues (brain, muscle, midgut, ovary, and carcass (oenocyte and fat body)) were dissected in ice-cold PBS for whole mount experiments. Or whole flies were embedded into a Tissue-Tek Cryomold (SAKURA, #4566) with O.C.T. compound (SAKURA, #4583) and serial 20-μm fresh- frozen cryostat sections were collected for large-scale staining. Dissected tissues or sectioned slides were fixed in 4% paraformaldehyde in PBS at room temperature for 30 min, permeabilized in PBST buffer (0.1% Triton X-100 in PBS) for 30 min, incubated for 1 h in blocking buffer (5% normal goat serum (Vector labs, S-1000), 0.1% Triton X-100 in PBS) and stained with primary antibodies overnight at 4 °C in the blocking buffer. The primary antibodies and their dilutions were as follows: rabbit anti-ref2P/Ref(2)P (Abcam, 178440; 1:1000), rabbit anti-atg8a/ GABARAP+GABARAPL1+GABARAPL2 (Abcam, 109364; 1:500), rabbit anti-atp5a (Abcam, 151229; 1:1000), mouse anti-atp5a (Abcam, 14748; 1:1000), mouse anti-FK2/ubi (Enzo Life Sciences, BML- PW8810; 1:1000), mouse anti-UBCJ2/ubi (Enzo Life Sciences, ENZ-ABS840; 1:1000), Chicken anti- GFP (Abcam ab13970; 1:3000) and mouse anti-Flag (Sigma, F1804; 1:1000). After incubation with primary antibody, the samples were washed five times with PBST and stained with DAPI (1:1000 dilution) and secondary antibodies at room temperature for 2 hours. The secondary antibodies and their dilutions were as follows: Alexa Fluor 488 anti-rabbit (Invitrogen, A11034; 1:1000), Alexa Fluor 488 anti-mouse (Abcam, 150113; 1:1000), Alexa Fluor 594 anti-mouse (Invitrogen, A11005; 1:1000), Alexa Fluor 594 anti-rabbit (Invitrogen, A11037; 1:1000), Alexa Fluor 647 anti- mouse (Invitrogen, A21235; 1:1000), Alexa Fluor 568 anti-Rabbit (Invitrogen A11011; 1:1000) and Alexa Fluor 633 anti-rabbit (Invitrogen, A21071; 1:1000). Samples were washed five times with PBST and mounted in Vectashield medium (Vector lab, H-1000).

#### Imaging acquisition and visualization

All images of fly tissues and cells shown in this study are confocal images acquired with either Leica SP8 confocal or Nikon spinning disk systems. For Leica SP8, we used several objectives including HC PL APO CS2 10x objective lens N.A. 0.40, HC PL APO 63x objective lens N.A. 1.4 Oil and HC PL APO CS2 100x objective lens N.A. 1.4 Oil. For DAPI we used 405 nm laser excitation (White light laser) on PMT detector with emission detection band 420-494 nm, GFP channel used 488 nm laser excitation (White light laser) on HyD detector with emission detection band 495- 537 nm, red channel used 594 nm laser excitation (White light laser) on HyD detector with emission detection band 600-645 nm and far-red channel used 647 nm laser excitation (White light laser) on HyD detector with emission detection band 655-700 nm. For Nikon spinning disk, images were acquired with an Orca Flash 4.0 sCMOS at full resolution. We used several objectives including Plan Apochromat Lambda 60x oil objective lens, N.A. 1.4, 0.228 um/px and Plan Apochromat Lambda 100x oil objective lens, N.A. 1.49, 0.185 um/px. The lasers used include 405 nm/ 488 nm/ 561 nm/ 640 nm with emission filters 430-480 nm (DAPI), 507-543 nm (GFP), 570- 640 nm (Red) and 662.5-737.5 nm (Far-red) respectively.

Certain images captured using the Leica system underwent deconvolution using Huygens Professional software (version 14.10; Scientific Volume Imaging). The model of theoretically calculated point spread function (PSF) was applied, while all other parameters were maintained at their default values based on the acquisition setting mentioned above. The background was determined from the original raw image, and the signal-to-noise ratio was adjusted within the range of 15 to 20.

#### Ref(2)P aggregates quantification and colocalization analysis

For quantification of Ref(2)P aggregates, Ref(2)P channel from a z-stack series of confocal image was isolated in the biological-image analysis software Fiji and processed as a maximum projection from 15-20 stacks (0.3 μm/ stack) depending on the tissue types. Ref(2)P signals were segmented by applying the ‘Subtract Background’ and ‘Auto Threshold v1.18.10 Method_Yen’ functions and analyzed by the ‘Analyze Particles’ function.

For colocalization analysis of Ref(2)P and ubiquitin with other subcellular marker such as ATP5A or atg8a, an object-based colocalization method was applied. Specifically, two montage images were generated from a z-stack series of confocal images for Ref(2)P/ubi channel and atg8a/ATP5A channel separately. Each channel was segmented by applying the ‘Subtract Background’ and ‘Auto Threshold v1.18.10 Method_Yen (for Ref(2)P, ubi, atg8 signals) and Auto Threshold v1.18.10 Method_Percentile (for atp5a signals)’ functions. Overlapping areas between two segmented channels were analyzed by applying the ‘imageCalculator_AND’ and created as a new overlapping channel. Colocalization percentage was calculated by the ratio of overlapping channel versus Ref(2)P channel from results analyzed by the ‘Analyze Particles’ function.

## Silver enhanced Immuno-EM

Fly tissues (muscle, midgut and carcass(oenocyte)) were dissected in ice-cold PBS, fixed in 4% paraformaldehyde and 0.01% Glutaraldehyde in PBS at room temperature for 30 min, permeabilized in 0.1% saponin (Sigma, S4521) in PBS for 30 min, incubated for 1 h in blocking buffer (5% normal goat serum (Vector labs, S-1000), 0.03% saponin in PBS) and stained with primary antibodies overnight at 4 °C in the blocking buffer. Primary antibodies used in this study and their dilutions were as follows: rabbit anti-ref2P/Ref(2)P (Abcam, 178440; 1:750), mouse anti-Flag (Sigma, F1804; 1:1000). After incubation with primary antibody, the tissues were washed five times with 0.03% saponin in PBS and incubated with Goat-anti-Rabbit IgG H&L Ultra Small (Aurion, #25100; 1:50) or Goat-anti-Mouse IgG+IgM Ultra Small (Aurion, #25160; 1:50) immunogold reagents (Molecular Probes/Invitrogen; 1:1,000) overnight at 4 °C in the blocking buffer. After incubation with secondary antibody, samples were washed five times with 0.03% saponin in PBS and then fixed in 50 mM Sodium Cacodylate buffer (pH 7.4) containing 2.5% Paraformaldehyde and 2% Glutaraldehyde. The nano-gold signal was enlarged using Aurion R- Gent SE-EM kit (Aurion, #500.033). The samples were then post fixed with 0.5% OsO4 for an hour, and 1% Uranyl Acetate overnight. After dehydration with a graded Ethanol series, samples were infiltrated and embedded into Epon resin (EMS, Fort Washington, PA). Ultrathin (100 nm) sections were collected on slot grids, stained with 4% Uranyl Acetate in 70% Methanol and 2% Lead Citrate. Sections were imaged using a FEI transmission electron microscope at 80 kV.

## APEX2-EM and Corelative Light and Electron Microscopy (CLEM)

S2 cells expressing Ref(2)P-APEX2 (for APEX2 see reference ^91^) and Ref(2)P-mEOS4b (for mEOS4b see reference ^92^) were cultured in 35 mm petri-dish with gridded coverslip (P35G-1.5-14-C-GRD, Mattek) and fixed with 4% PFA in PBS. Confocal fluorescent images were taken before EM processing for LM-EM correlation. The samples were further fixed with 2% glutaraldehyde after fluorescence imaging, and then subjected to 3,3′-Diaminobenzidine (DAB) treatment. Following DAB reaction, the samples were washed with cacodylate buffer, fixed again in 2% glutaraldehyde for an hour to stabilize the DAB reaction product. Subsequently, the samples were fixed with 0.5% Osmium for half an hour, washed, stained overnight with 0.5% uranyl acetate. The samples were then serial dehydrated with ethanol and embedded into Epon resin. Ultrathin sections were cut at 100 nm on an ultramicrotome (UC6, Leica) and imaged using a Tecnai Biotwin electron microscopy operated at 80 kV.

## Histology staining

### H&E staining of paraffin-embedded adult thorax

Dissected Drosophila thoraxes were placed in 4% PFA and fixed overnight at 4 °C. The tissue was then rinsed in 1x PBS thoroughly and dehydrated in 30%, 50%, and then 70% ethanol in preparation for paraffin processing. Tissue was then processed for paraffin embedding, and thoraxes were individually embedded in paraffin under a dissection scope allowing the tissue to be cut at a transverse orientation. Tissue blocks were sectioned serially by microtome (Leica Hi1220 AUTOCUT automated rotary microtome) at 5 um section thickness. Sections were checked for artifacts created by the chitin exoskeleton, and block was soaked in 10% formic acid (TBD-2 – Epredia – 6764003) for 10 minutes if surface decalcification was needed. Sections were collected on charged Microscope slides (Avantik SureBond Slides – SL6332-1) and dried in a 37 °C oven overnight. Slides were then heated at 60 °C for an hour to remove excess paraffin wax and promote good tissue adherence to the slide. Slides were stained on the Automated H&E stainer (Dakewe Dp360+CS500 Combo System) with Program 1 and cover slipped with xylene-based mounting media (CytoSeal XYL – VWR – 48212-196).

### Toluidine blue staining of resin-embedded adult thorax

We used a published protocol ^93^. Briefly, dissected Drosophila thoraxes were fixed in 4% PFA and 2% glutaraldehyde, treated by OsO4 and embedded in epoxy resin after dehydration through a graded ethanol series. Samples were sectioned with 1 um thickness in the microtome and stained by toluidine blue solution (1% of toluidine blue and 1% of borax in H2O).

## Extraction of soluble and insoluble fractions

Samples were homogenized in the “soluble” buffer (0.1% Triton X-100 in PBS with protease inhibitors), incubated 30 min at 4 °C and cleared by centrifugation at 18000xg for 10 min at 4 °C. Supernatants were collected as soluble factions. Pellets were washed three times with soluble buffer and collected by centrifugation at 18000g for 3 min at 4 °C after each wash. Washed pellets were then dissolved in the “insoluble” buffer (8 M UREA, 5% SDS and 0.1% Triton X-100 in PBS with protease inhibitors mentioned above), incubated 30 min at 4 °C and denatured with 4X sample loading buffer for 5 min at 95 °C. Supernatants were collected as insoluble factions after a spin of 2000g for 30 s.

## Co-immunoprecipitation

Whole flies or dissected fly muscles were homogenized in lysis buffer (50 mM sodium phosphate buffer (pH 7.5), 150 mM sodium chloride, 0.5% IGPAL CA-630, 1 mM PMSF, 1 mM MgCl_2_, one complete EDTA-free protease inhibitor cocktail tablet, 0.01% Benzonase). The lysates were incubated at 4 °C for 30 minutes, and then centrifuged at 4,000g for 10 minutes. Supernatants were collected and incubated with pre-washed flag beads (EZview™ Red ANTI-FLAG® M2 Affinity Gel, F2426) for one hour at 4 °C. The incubated beads were then washed with a wash buffer (50 mM sodium phosphate buffer (pH 7.5), 150 mM sodium chloride, 0.5% IGPAL CA-630). The washed beads were eluted with 1X SDS sample buffer at 90 °C for 10 minutes. Supernatants were collected for western blot.

## Western blot

Samples were lysed in lysis buffer (RIPA buffer [25 mM Tris-HCl, pH 7.6; 150 mM NaCl; 1% NP-40; 1% sodium deoxycholate and 0.1% SDS] or urea buffer [5% SDS; 8M urea] was used unless noted otherwise) with 1 mM phenylmethanesulfonylfluoride (Sigma, 93482) and appropriate amount of EDTA-free protease inhibitor (Roche, 11873580001; 1 tablet per 10 ml buffer). An additional 50 μM RP-619, inhibitor of the deubiquitylating enzymes (Sigma, 662141) was added when ubiquitinated protein level was measured. Dissected tissues and whole flies were homogenized using Disposable Pellet Mixers and Cordless Motor (Avantor-VWR, 47747-370 and 47747-366). After 30 min incubation at 4 °C, cell/tissue lysates were cleared by centrifugation at 18000xg for 10 min at 4 °C. Supernatants were collected, protein concentration was measured by Pierce BCA Protein Assay Kit (Thermo Fisher, 23227), normalized into equal concentration, denatured with 4X sample loading buffer (200 mM Tris-HCl, pH=6.8; 8% SDS; 40% glycerol; 4% β- mercaptoethanol; 50 mM EDTA; 0.08% bromophenol blue) for 5 min at 95 °C, resolved in NuPAGE 4-12% Bis-Tris Protein Gels (Thermo Fisher, NP0326) and analyzed by immunoblotting. Primary antibodies and their dilutions in the blocking buffer (5% milk in 1× Tris-buffered saline (TBS) containing 0.1% Tween-20) were as follows: rabbit anti-ref2P/Ref(2)P (Abcam, 178440; 1:1500), rabbit anti-atg8a/ GABARAP+GABARAPL1+GABARAPL2 (Abcam, 109364; 1:3000), rabbit anti-GFP (MBL, 598; 1:3000), mouse anti-P4D1/ubi (Cell signaling, 3936; 1:1500), mouse anti-UBCJ2/ubi (Enzo Life Sciences, ENZ-ABS840; 1:1500), mouse anti-atp5a (Abcam, 14748; 1:1500), mouse anti- Flag (Sigma, F1804; 1:1500), mouse anti-actin/C4 (Sigma, MAB1501; 1:10000), Chicken anti-GFP (Abcam ab13970; 1:2000) and mouse anti-HSP70 (Thermo Fisher, 7FB, 1:100) Second antibodies and their dilutions were as follows: goat anti-rabbit IgG, HRP-linked Antibody (Cell Signaling, 7074; 1:3000), anti-chicken IgG, HRP-linked antibody (Thermo Fischer A9792, 1:3000) and horse anti-mouse IgG, HRP-linked Antibody (Cell Signaling, 7076; 1:3000).

## Protein purification

### Recombinant protein expression and purification from *E. coli*

For bacterial protein expression, 6XHis-MBP-3C-sfGFP, 6XHis-MBP-3C-Ref(2)P-WT-sfGFP, 6XHis- MBP-3C-9xUbiquitin, 6XHis-MBP-3C-Atg8a and 6XHis-GFP nanobody proteins were expressed in *E. coli* Rosetta (DE3) cells. Cells were induced with 1 mM IPTG at an OD_600_ of 0.5-1 for 18 hours at 25 °C. Bacteria pellets were collected and frozen in -80 °C until use.

To purify bacterial proteins, bacteria pellets were homogenized in lysis buffer (50 mM sodium phosphate buffer (pH 7.5), 150 mM sodium chloride, 20 mM imidazole, 0.5% IGPAL CA-630, 1 mM PMSF, one complete EDTA-free protease inhibitor cocktail tablet, 0.2% Lysozyme). The lysates were incubated at 4 °C for 30 minutes, and then centrifuged at 18,000g for 30 minutes to remove debris. Supernatants were collected and filtered with 0.45 μm polyether sulfone membrane. Cleaned supernatants were then injected into a column of Ni-NTA resin equilibrated with buffer A (50 mM sodium phosphate buffer (pH 7.5), 500 mM sodium chloride, 20 mM imidazole) in AKTA Pure chromatography system. The columns were washed with buffer A until OD280 reached baseline and then eluted with buffer B (50 mM sodium phosphate buffer (pH 7.5), 500 mM sodium chloride, 300 mM imidazole). OD280 peak fractions were collected and dialyzed overnight with dialysis buffer (50 mM sodium phosphate buffer (pH 7.5), 150 mM sodium chloride). Protein samples were aliquoted and stored in -80 °C until use. For 6XHis-MBP- 3C-Ref(2)P-sfGFP protein, HRV 3C protease was used to remove MBP tag. Cleaved protein samples were then passed through column of Ni-NTA resin equilibrated with buffer A (50 mM sodium phosphate buffer (pH 7.5), 500 mM sodium chloride, 20 mM imidazole). The flowthrough with Ref(2)P-sfGFP proteins were collected and stored at -80 °C until use.

### GFP nanobody-based protein purification from fly cell line and tissue lysates

For Ref(2)P-sfGFP-Flag protein expression in S2 cells, S2 cells were transfected with pAWG- Ref(2)P-sfGFP-Flag plasmid and cultured for two days. The transfected S2 cells were collected and stored at -80 °C until use. For Ref(2)P-sfGFP-Flag protein expression in different fly tissues, Ref(2)P-sfGFP-Flag protein was expressed in specific tissues driven by tissue-specific GAL4s (See genotype table for more details). Millions of adult flies were collected and stored at -80 °C until use.

To purify Ref(2)P-sfGFP-Flag protein from S2 cell and fly tissue, S2 cell pellets or adult flies were homogenized in lysis buffer (50 mM sodium phosphate buffer (pH 7.5), 150 mM sodium chloride, 20 mM imidazole, 0.5% IGPAL CA-630, 1 mM PMSF, one complete EDTA-free protease inhibitor cocktail tablet, 0.01% Benzonase). The lysates were incubated at 4 °C for 30 minutes, and then filtered through Mira cloth, followed by centrifugation at 4,000g, 4 °C for 10 minutes. Supernatants were collected and precleaned through Ni-NTA column equilibrated with buffer A (50 mM sodium phosphate buffer (pH 7.5), 500 mM sodium chloride, 20 mM imidazole) in AKTA Pure chromatography system. Flowthrough lysates were collected and incubated with Histidine-tagged GFP nanobody at 4 °C for one hour. GFP-His nanobody incubated lysates were then injected into a fresh Ni-NTA column equilibrated with buffer A (50 mM sodium phosphate buffer (pH 7.5), 500 mM sodium chloride, 20 mM imidazole). The columns were washed with buffer A until OD280 reached baseline and then eluted with buffer B (50 mM sodium phosphate buffer (pH 7.5), 500 mM sodium chloride, 300 mM imidazole). OD280 peak fractions were collected and dialyzed overnight with size exclusion buffer (50 mM sodium phosphate buffer (pH 7.5), 300 mM sodium chloride).

Size exclusion was then used to further remove contaminants and GFP nanobody. Ni-NTA column purified proteins were loaded into Superose® 6 size exclusion column 10/300 GL (Sigma-Aldrich, GE29-0915-96). OD280 peak fractions were collected and targeted proteins were conformed with western blot and silver staining. Protein samples were then dialyzed overnight with dialysis buffer (50 mM sodium phosphate buffer (pH 7.5), 150 mM sodium chloride). Samples were aliquoted and stored at -80 °C until use.

## Mitochondrial isolation

w1118 or Ref(2)P^-/-^ mutant adult flies were gently homogenized in chilled isolation buffer A (100 mM Tris HCl, 225 mM Sorbitol, 60 mM KCl, 1 mM EDTA, 0.1% BSA, adjust the pH to 7.2 using sodium hydroxide; Filter with 0.22 um pore size). Lysates were then centrifuged at 3,000xg for 5 minutes at 4 °C to remove debris. The pellets were collected and resuspended in isolation buffer Resuspended pellets were centrifuged at 15,000xg for 15 minutes at 4 °C. Pellets were then resuspended in isolation buffer A, loaded on the top of 18%-23%-40% percoll gradient and centrifuged at 60,000g for one hour at 4 °C. Mitochondrial fraction at the interface of 23%-40% layer was extracted carefully and diluted with suspension Buffer C (100 mM Tris HCl, 250 mM Sorbitol, 60 mM KCl, 0.05 mM EDTA). Mitochondrial fractions were then centrifuged at 20,000xg for 15 minutes at 4 °C. Pellets were stored for further use.

### *In vitro* binding assay

#### Ref(2)P and Atg8a binding

Recombinant 6XHis-MBP-3C-9xUbiquitin and 6XHis-MBP-3C-Atg8a proteins were incubated with purified Ref(2)P proteins (Bacteria, S2 cell, fly midgut, fly muscle and fly oenocytes) in binding buffer (50 mM sodium phosphate buffer (pH 7.5), 150 mM sodium chloride, 20 mM imidazole) for one hour at 4 °C. GFP-His nanobody was then added into the solution and incubated for another one hour at 4 °C. The mixture was passed through Ni-NTA column equilibrated with buffer A (50 mM sodium phosphate buffer (pH 7.5), 500 mM sodium chloride, 20 mM imidazole). The columns were washed with buffer A and then eluted with buffer B (50 mM sodium phosphate buffer (pH 7.5), 500 mM sodium chloride, 300 mM imidazole). Eluents were stored for further characterization.

#### Ref(2)P and Mitochondria binding

Isolated mitochondria pellets were resuspended in binding buffer (100 mM Tris HCl, 250 mM Sorbitol, 60 mM KCl, 0.05 mM EDTA) and incubated with purified Ref(2)P proteins (Bacteria, S2 cell, fly midgut, fly muscle and fly oenocytes) for one hour at 4 °C. The mixture was then centrifuged at 20,000Xg for 15 minutes at 4 °C to pellet mitochondria. Mitochondria-bound Ref(2)P was recovered in the pellet. Unbound Ref(2)P stayed in the supernatant. Pellets were stored for further characterization.

## Thermal shift assay

The thermal shift assays were performed following the published protocol ^94^ with some optimization. Purified proteins or whole lysates were heated in a temperature gradient from 35 °C to 65 °C for 3 minutes in a PCR instrument (Eppendorf, Mastercycler proS). Heated samples were transferred to new Eppendorf tubes and centrifuged at 20,000xg for 15 minutes at 4 °C. Supernatants, which contain the soluble proteins, were collected for western blot analysis.

## Limited proteolysis assay

An equal amount of purified proteins was incubated with 0; 0.1 ug/ml; 1 ug/ml; 2 ug/ml; 10 ug/ml; 100 ug/ml of papain enzyme (Worthington PAP2, LK003178) prepared in PBS buffer. After a 10-minute incubation at 37 °C, reactions were stopped by adding 4X sample loading buffer and immediately incubating at 95 °C for 5 min. Samples were then analyzed by western blot.

## Crosslinking and mass spectrometry analysis

### Crosslinking reaction and peptide preparation

Purified proteins were incubated with 5 mM DSSO (Sigma-Aldrich,909602) or 20 mM DMTMM (Sigma-Aldrich,74104) in reaction buffer (50 mM sodium phosphate buffer (pH 7.5), 150 mM sodium chloride) for one hour or indicated time at room temperature. For DMTMM and Dihydrazide sulfoxide (DHSO, Sigma, 909599) crosslinking, 20 mM DMTMM and 20 mM DHSO were added into protein samples and incubated for one hour at room temperature. All reactions were stopped with 8 M urea in 50 mM Tris-HCl (pH 8.0). For in-gel digestion, the samples were loaded and run SDS-PAGE gel. Targeted bands were cut and then processed with in-gel digestion protocol ^95^. Protein samples were incubated with 50 mM Tris(2-carboxyethyl) phosphine hydrochloride (TCEP, Sigma-Aldrich, 51805-45-9) for 30 minutes at room temperature, and in 50 mM Iodoacetamide (Sigma-Aldrich, A3221) for another 30 minutes at room temperature. The samples were then diluted four times with 100 mM Tris-HCl (pH 8.0) and pH was adjusted to 8.0.

Samples were first digested with rLys-C (Promega, V167A) at a ratio of 1:100 (rLys-C to protein amount) for two hours at 37 °C, and then digested with Trypsin (Promega, V5280) at a ratio of 1:100 (Trypsin to protein amount) for 16 hours at 37 °C. The digested peptide samples were desalted with Pierce^TM^ C18 spin tips (ThermoFisher, 84850) following the protocol provided by the manufacturer. Desalted crosslinking peptide samples were dried and stored at -80 °C until use.

### Mass spectrometry acquisition of crosslinking peptides

Cross-linked peptides were resolved for mass spectrometry analysis using a Dionex Ultimate 3000 RSLCnano liquid chromatography system. Peptides were initially loaded from the autosampler onto an Acclaim™ PepMap™ 100 C18 LC Trap Cartridge (0.3 mm inside diameter, 5 mm length) (Thermo Scientific, San Jose, CA) using a loading pump flow rate of 8 nl/minute. Peptides were subsequently resolved for mass spectrometry analysis using an analytical column (75 μm inside diameter, 250 mm length) packed in-house with ReproSil®-Pur C18-aQ 1.9 μm resin (Dr. Masch GmbH, Germany). Chromatography was performed using combinations of buffer A (95% water, 5% acetonitrile, and 0.1% formic acid (v/v/v), pH 2.6), and buffer B (20% water, 80% acetonitrile, and 0.1% formic acid (v/v/v), pH 2.6). The following chromatography steps were performed using a flow rate of 180 nl/minute: (1) 2% B for 25 minutes (column equilibration); (2) a linear gradient from 2% to 10% B over 2 minutes; (3) a linear gradient from 10% to 33% B over 78 minutes; (4) a linear gradient from 33% to 55% B over 12 minutes; (5) a linear gradient from 55% to 95% B over 8 minutes; (6) 95% B for 5 minutes; (7) a linear gradient from 95% B to 2% B over 1 minute; (8) 2% B for 10 minutes.

Eluted peptides were analyzed by mass spectrometry using an Orbitrap Fusion™ Lumos™ mass spectrometer (Thermo Scientific, San Jose, CA). An MS3 based method was used for identification of DHSO/DSSO cross-linked peptides as follows: Full MS scans were performed using the Orbitrap mass analyzer (60,000 m/z resolution, 1.6 m/z isolation window, and 375-1500 m/z scan range); The top 3 peptides identified with charge state 4 to 8 were selected for MS2 fragmentation (20% CID energy) and subsequent detection with the Orbitrap mass analyzer (30,000 m/z resolution and a dynamic exclusion time of 40 s); Pairs of MS2 fragments with a mass difference of 31.9720 (20 ppm mass tolerance) were selected for MS3 fragmentation (HCD energy 35%) and detection using the Linear Ion Trap mass analyzer (rapid scan, 3 m/z isolation window, maximum ion injection time 200 ms); Each MS2 scan was followed by a maximum of 4 MS3 scans. In PRM, targeted mass triggers were applied at MS1 using inclusion list for to each FAIMS setting. Raw data and search results files have been deposited to **MassIVE MSV000097980 (**ftp://MSV000097980@massive-ftp.ucsd.edu)

### Searching for cross-linked amino acids

Proteome Discoverer v3.0 (Thermo Fisher Scientific) was used for PTM analysis, with dynamic modifications including phosphorylation on S, T, and Y (+79.996 Da), acetylation on K (+42.011 Da), and M oxidation (+15.995 Da). Crosslinking analysis using DSSO and DMTMM was performed with Proteome Discoverer v3.0 integrated with XlinkX v3.0 ^55^. The detailed searching includes peptide mass (500–10,000 Da); minimum peptide length (5 residues), precursor mass tolerance (±10 ppm), product-ion mass tolerance for Orbitrap data (±20 ppm), mass tolerance for ion trap data (±0.5 Da), modifications (oxidation (M), carbamidomethyl (C)), and 1% FDR control for CSMs. For DSSO cross-links, lysine (K), Serine (S), Threonine (T) and Tyrosine (Y) residues were used for cross-links with a mass shift 158.0038 Da, cleavage alkene fragment mass of 54.0106 Da and thiol fragment mass of 85.9826 Da. For DHSO /DMTMM cross-links, DHSO cross-links were searched with asparanin acid (D) and Glutamine acid (E) for a mass shift 186.0575 Da, cleavage alkene fragment mass of 68.0375 Da and thiol fragment mass of 100.0095 Da. As DMTMM can promote the crosslink reaction between K and D/E ^96^, DMTMM non-cleavable cross-links were searched with a mass shift -18.0105 Da. The data were searched against both whole database (Uniprot *Drosophila Melanogaster* reference proteome plus contaminant sequences and Ref(2)P-sfGFP- Flag sequence) and small database that were generated by ID characterization of purified protein samples.

## Tandem Mass Tag (TMT) labeling for whole proteomics

Immediately before use, the vials containing frozen TMT 16plex reagents were allowed to warm up at room temperature. To each of the 0.5 mg vials, 20 μl of anhydrous acetonitrile were added. The TMT reagents were let to dissolve for 5 min with occasional vortexing, then the tubes were briefly centrifuged to gather the solution. 100 μg of peptides were measured out based on the peptide assay, and the volumes were adjusted to 100 μl with 100 mM TEAB, then added to the TMT vials. The labeling reaction was allowed to proceed for 1 hour at room temperature. To test labeling efficiency, 1 μl of each sample was analyzed by LC/MSMS over a 2-hour C18 reverse- phase (RP) gradient on an Orbitrap Eclipse Tribrid Mass Spectrometer (Thermo Scientific) with a FAIMS Pro interface, equipped with a Nanospray Flex Ion Source, and coupled to a Dionex UltiMate 3000 RSCLnano System. The TMT labeling levels were >99% for all detected peptides (data not shown). The differentially labeled samples (10 μl each) were combined in a new tube and the volume was reduced to less than 10 μl (about 1 hour) using a SpeedVac concentrator (Savant).

## High pH reverse phase fractionation and Mass spectrometry data acquisition

The dried TMT-labeled peptide mixture was resuspended in 300 μl of 0.1% trifluoroacetic acid (TFA). One high pH fractionation cartridge (Pierce, cat. # 84868) was placed on a new 2.0 ml sample tube and 300 μl of the TMT-labeled peptide mixture were loaded onto the column. After centrifuging at 3000×g for 2 minutes, the eluate was collected as the “flow-through” fraction. The loaded cartridge was placed on a new 2.0 ml sample tube, washed with 300 μl of ddH20, and the eluate collected as “wash” fraction. An additional round of washing was performed using 300 μl of 5% acetonitrile in 0.1% TFA to remove unreacted TMT reagent. A total of 9 HpH RP fractions were collected by sequential elution in new sample tubes using 300 μl of 10%, 12.5%, 15%, 17.5%, 20%, 22.5%, 25%, 50%, and 80% acetonitrile in 0.1% TFA. The solvents evaporated to dryness using vacuum centrifugation. Dried samples were resolubilized in 44 μl of buffer A (5% acetonitrile in 0.1% formic acid, FA) before LC-MS analysis.

TMT-labeled peptides were analyzed on Orbitrap Eclipse Tribrid Mass Spectrometer (Thermo Scientific) with a FAIMS Pro interface, equipped with a Nanospray Flex Ion Source, and coupled to a Dionex UltiMate 3000 RSCLnano System. Peptides (22 μl for each HpH RP fraction) were loaded on an Acclaim PepMap 100 C18 trap cartridge (0.3 mm inner diameter (i.d.), 5 mm length; Thermo Fisher Scientific) with the U3000 loading pump at 2 μL/minute via the autosampler.

A 75 μm i.d. analytical microcapillary column was packed in-house with 250 mm of 1.9 μm ReproSil-Pur C18-AQ resin (Dr. Masch). AgileSLEEVE (Analytical Sales & Products) was used to maintain column temperature at 40 °C. The organic solvent solutions were water:acetonitrile:formic acid at 95:5:0.1 (volume ratio) for buffer A (pH 2.6) and 20:80:0.1 (volume ratio) for buffer B. The chromatography gradient was a 30 min column equilibration step in 2% B; a 5 min ramp to reach 10% B; 90 min from 10% to 40% B; 10 min to reach 90% B; a 5 min wash at 90% B; 0.1 min to 2% B; followed by a 15 min column re-equilibration step in 2% B. The nano pump flow rate was set to 0.180 μl/min. For whole proteome TMT labeled high pH reversed fractioned samples, the Orbitrap Eclipse was set up to run the TMT-SPS-MS3 method with three FAIMS compensation voltages (CV) at -40 V, -55 V, and -70 V and a cycle time of 1 sec. Briefly, peptides were scanned from 400-1600 m/z in the Orbitrap at 120,000 resolving power before MS2 fragmentation by CID at 35% NCE and detection in the ion trap set to turbo detection. Dynamic exclusion was enabled for 45 s. Real time search (RTS) was performed during the ddMS2 IT CID step against a non-redundant Drosophila melanogaster sequence database downloaded from NCBI 2020-04-23 and complemented with common contaminants. Carbamidomethyl (57.0215 Da on Cys) and TMTpro16plex (304.2071 Da on Kn) were searched statically, while methionine oxidation (15.9949 Da) was searched as a variable modification. Synchronous precursor scanning (SPS) selected the top 10 MS2 peptides for TMT reporter ion detection in the Orbitrap using HCD fragmentation at 65% NCE at 50,000 resolving power.

## RNA-Seq and Analysis

Total RNA was extracted from fly muscle and gut tissues with Maxwell® RSC simplyRNA Tissue Kit (Promega, AS1340), and library was generated by the TruSeq Stranded mRNA Preparation Kit (Illumina), followed by sequencing on an Illumina HiSeq2000 for 100 bp single reads. Raw reads were demultiplexed into Fastq format allowing up to one mismatch using Illumina bclconvert 3.10.5. Reads were aligned to UCSC genome dm6 with STAR aligner (version 2.7.3a), using Ensembl 106 gene models. TPM values were generated using RSEM (version v1.3.0). Three biological replicates were sequenced for each experimental condition.

## Structural modeling

### Alphafold2 and Alphafold3 prediction and local refinements with crosslinking

Ref(2)P protein structural models were generated by Alphafold2 ^97^ implemented in GPU clusters of Stower Computation Center or on-line Alphafold3 ^98^. Alphafold2 structures were predicted using ColabFold notebook ^99^ with alphafold_multimer_v3 model, full MSA, 20 recycles, seed (0). Unrelaxed structures were used for analysis. For domain-domain interaction, well-folded domains and IDRs of Ref(2)P were first identified according to predicted aligned error (PAE) with boundary PAE<10 Å and then subjected to Alphafold2 or Alphafold3 for interaction prediction. The interaction matrix was generated according to pTM or ipTM score. Crosslinking information was then used to locate the regions for structural refinement. For intra-chain crosslinking of “N- C” form, the interaction prediction of PB1-sfGFP or PB1-UBA were used. For inter-chain homo- crosslinking on IDR2, the IDR2 sequence was split into three fragments – “SANQ” motif, its N- terminal and its C-terminal, and then interaction prediction was done with every two of the three fragments.

### AlphaLink modeling

AlphaLink ^57^ was used to incorporate crosslinking restraints into Alphafold2 prediction. *--flag* settings were as followed:

*N_eff_ size* (10); *checkpoint_path* (finetuning_model_5_ptm_CACA_10A.pt); and *FDR* (0.01). We compared predicted structures using both AlphaLink training models: *finetuning_model_5_ptm_CACA_10A.pt*

*finetuning_model_5_ptm_distogram.pt*

In --*distogram* model, the distance distribution was calculated with *FDR* (0.01) using script:

*preprocessing_distributions.py*

*--flag* settings were as followed:

#### N_eff_ size (10); checkpoint_path (finetuning_model_5_ptm_distogram.pt)

However, the --*distogram* model generated structures like Alphafold2 did, implying the model did not take the crosslinking data as constraints in prediction. So the *CACA_10A* model was used to generate AlphaLink structures for the following analysis.

### Rosetta comparative modeling with cross-link restraints

Rosetta-commons (released version 3.37) was installed in GPU clusters of Stower Computation Center. Rosetta comparative modeling was used to further incorporate crosslinking data in prediction following the published procedure ^56^ with some modifications. AlphaLink predicted structures and RoseTTAFold predicted structures were used as templates in the *<MOVER>* module of Rosetta *.xml* script. 200 structures were generated for each of three template sets with the three AlphaLink models for midgut- muscle- and bacteria- derived Ref(2)P protein. To generate crosslinking constraints, the *BOUNDED* function was used:

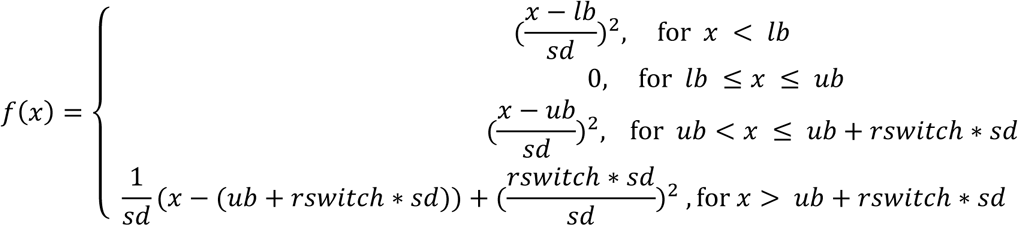

Settings of *BOUNDED* function were as follows: lb (10), ub (30), and rswitch (0.5).

Ref(2)P 3-mer and 9-mer fragments were generated with Fragment Server provided by David Baker’s lab (http://old.robetta.org/fragmentsubmit.jsp).

Rosetta module *rosetta_scripts.static.linuxgccrelease* was used for prediction and the flag *- constraints:cst_file* was used to incorporate crosslinking data. Three stages of weight were introduced for *SCOREFXNS* module of the Rosetta *.xml* script: Stage1.wts (env 1.0, pair 1.0, cbeta 1.0, cenpack 1.0, hs_pair 2.0, ss_pair 2.0, rsigma 2.0, sheet 2.0, vdw 0.2, rg 2.0, rama 0.3, linear_chainbreak 2.0, atom_pair_constraint 1.0); Stage2.wts (hbond_sr_bb 2.0, hbond_lr_bb 2.0, rama 0.2, omega 0.2, rg 2.0, vdw 1.0, cen_env_smooth 2.0, cen_pair_smooth 1.0, cbeta_smooth 1.0, cenpack_smooth 1.0, cart_bonded 0.05, atom_pair_constraint 0.5); and Stage2.wts (fa_atr 0.8, fa_rep 0.44, fa_sol 0.65, fa_intra_rep 0.004, fa_pair 0.49, fa_dun 0.56, ref 1, hbond_lr_bb 1.17, hbond_sr_bb 0.585, hbond_bb_sc 1.17, hbond_sc 1.1, p_aa_pp 0.32, dslf_ss_dst 0.5, dslf_cs_ang 2, dslf_ss_dih 5, dslf_ca_dih 5, pro_close 1.0, rama 0.2, omega 0.5, atom_pair_constraint 0.5, coordinate_constraint 0.0, cart_bonded 0.5)

Rosetta module *cst_info.static.linuxgccrelease* was used to score the structures for crosslinking data. The top 10 structures from each of the three predictions were further evaluated with ^100^ to measure Solvent Accessible Surface Distance. The best model was then used for analysis.

#### Structural visualization

*xiView* (https://xiview.org/xiNET_website/index.php) was used to calculate Cα-Cα Euclidean distance of crosslinking pairs and to generate *.pb* file for cross-link visualization. *ChimeraX* (https://www.rbvi.ucsf.edu/chimerax/) was used for structural visualization and crosslinking mapping.

## Negative staining of electronic microscopy

For negative staining, 4 µL of the sample was applied to glow-discharged carbon-coated 300- mesh copper grids (Electron Microscopy Sciences, PA, USA) and incubated for 2 minutes. Excess liquid was removed by blotting. The grids were then washed once and stained with 4 µL of 2% (w/v) uranyl acetate for 1 minute. The Imaging was performed using a Talos F200C transmission electron microscope (Thermo Fisher Scientific) operated at 200 kV. Micrographs were acquired with EPU software using a 4k × 4k Ceta 16M CMOS camera.

Micrographs were imported in CryoSPARC v4.7.0, following their specific negative stain data protocol (https://guide.cryosparc.com/processing-data/tutorials-and-case-studies/negative-stain-data). Particles were then picked using CryoSPARC’s blob-pick implementation, inspected and submitted for two rounds of 2D classification.

## Circular dichroism

Ref(2)P proteins, IDR2-sfGFP and control BSA or GFP were dialyzed with CD buffer (20 mM sodium phosphate plus 100 mM sodium chloride). The salt was applied to prevent Ref(2)P protein aggregation. Far-UV CD spectra of Ref(2)P proteins at a concentration of 0.1 μM, were recorded using a JASCO-J1500 spectropolarimeter (JASCO Inc.) equipped with a Peltier temperature control unit and using quartz cuvettes of 0.1-mmcell-path length. BSA and lysozyme, two proteins with significant content in *helical structures*, were used as controls. The secondary structure content was quantified by CD spectra deconvolution using the server BeStSel ^101^.

## Code availability

All the code for protein structure modeling was deposited to Github (https://github.com/Yan-Yan-2020/p62-protein-pleomorphism).

---

**Extended Data Fig. 1.**
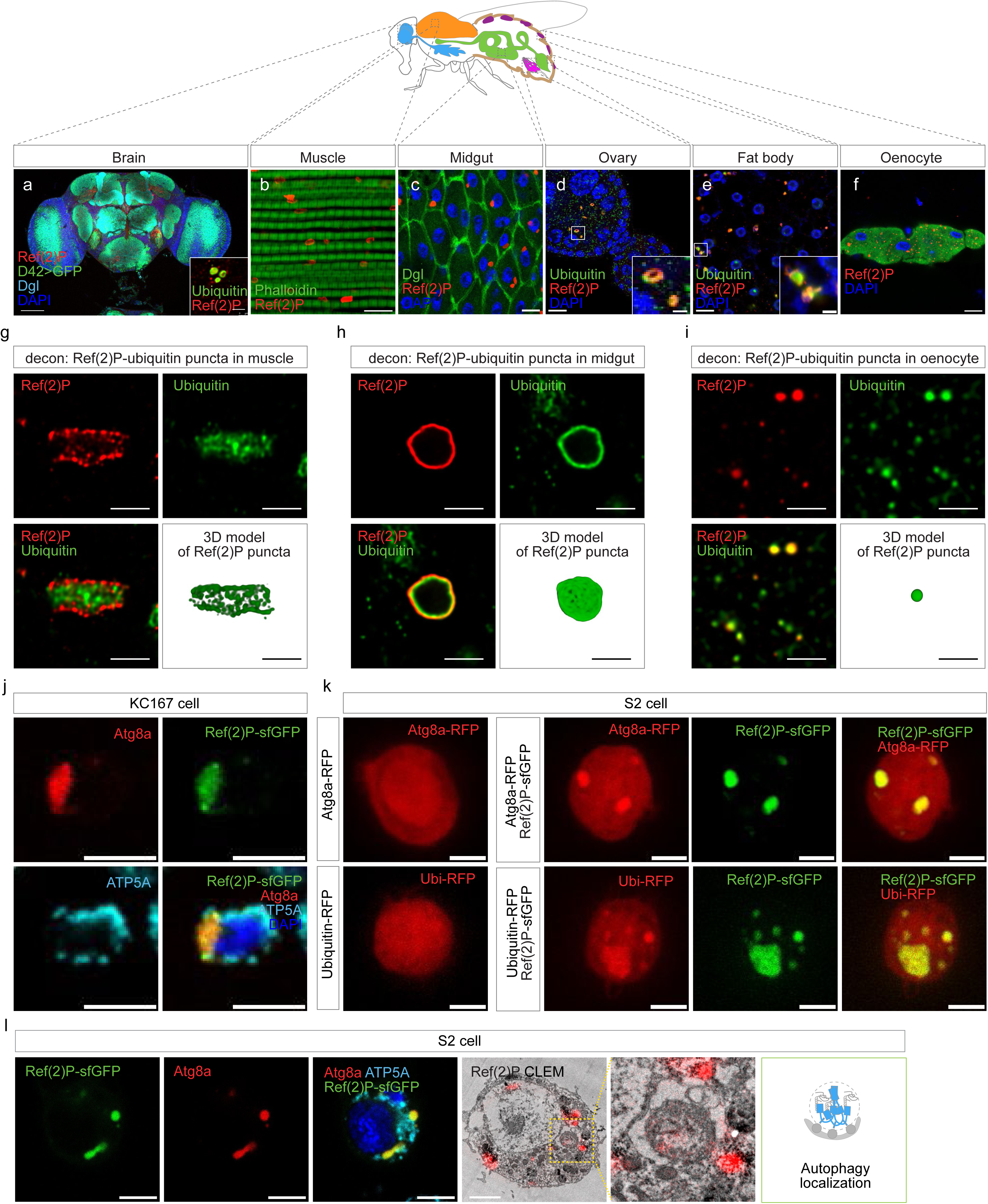
: Ref(2)P colocalizes with ubiquitin and occupies distinct compartments across tissues and cell lines. **a-f,** Immunofluorescence of Ref(2)P and ubiquitin in the indicated adult tissues. Tissue-specific GAL4-driven GFP marks neurons in brain (D42-GAL4) and oenocytes (Desat1-GAL4); Dlg, phalloidin and DAPI mark tissue architecture or nuclei as indicated. Insets in ovary and fat body show the boxed regions. Scale bar, 100 µm in brain, 10 µm in the other overview images and 2 µm in the insets. **g-i,** High-resolution deconvolved images and three-dimensional reconstructions of Ref(2)P/ubiquitin puncta in muscle (g), midgut (h) and oenocytes (i). Scale bars, 2 µm. **j,** KC167 cells expressing Ref(2)P-sfGFP and stained for Atg8a (red), ATP5A (cyan) and DAPI (blue). Scale bars, 5 µm. **k,** S2 cells expressing Atg8a-RFP or ubiquitin-RFP alone, or co-expressing each reporter with Ref(2)P-sfGFP. Scale bars, 5 µm. **l,** Fluorescence and correlative light-electron microscopy of Ref(2)P in S2 cells. Atg8a labels autophagic compartments, ATP5A labels mitochondria and the electron-dense peroxidase product from the Ref(2)P-mEos4b-APEX2 reporter is pseudocoloured red. The summary schematic assigns Ref(2)P to autophagic compartments. Scale bars 2 µm in both fluorescent images and EM images.

**Extended Data Fig. 2.**
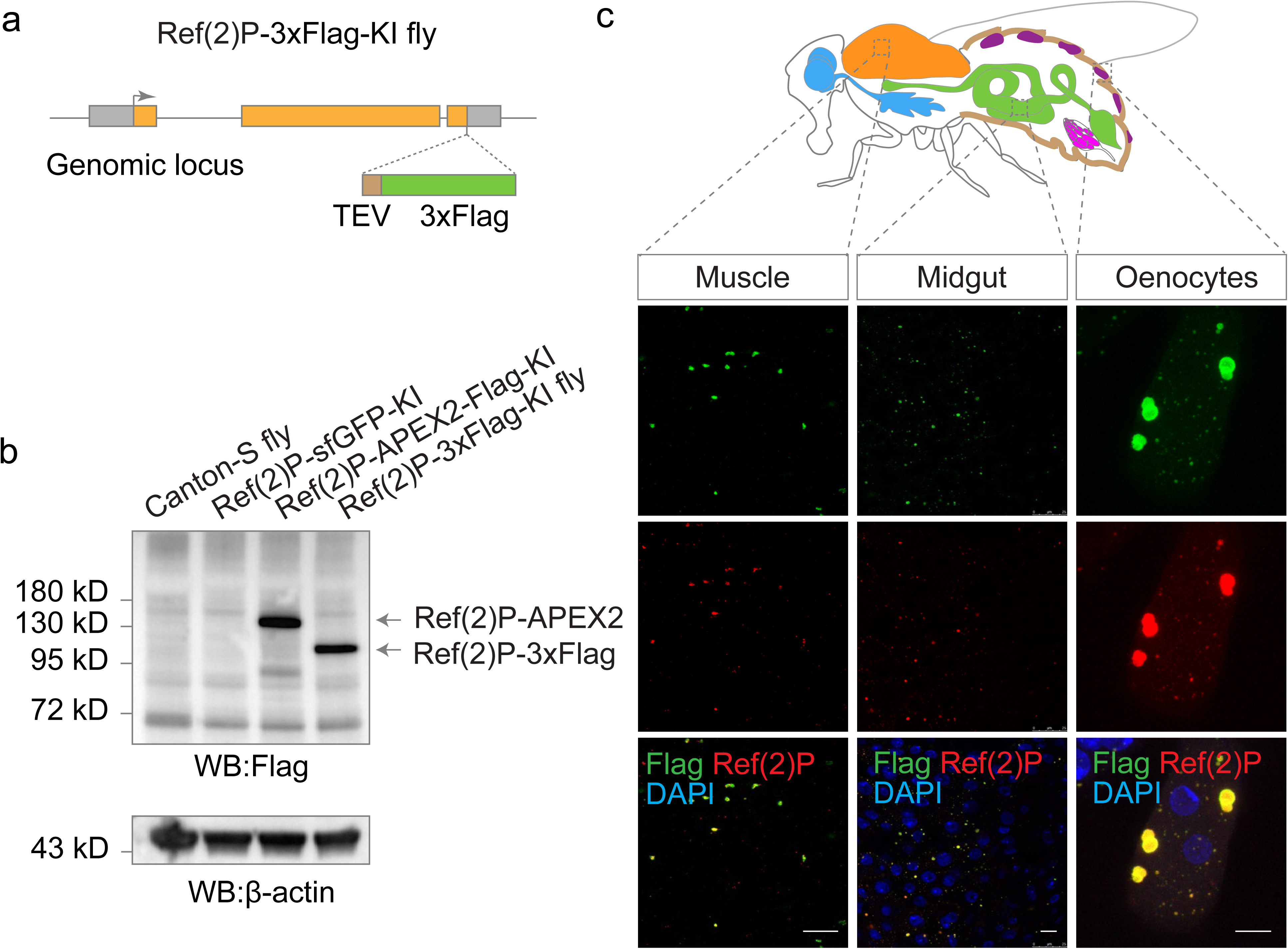
: Characterization of the Ref(2)P-3xFlag knock-in allele. **a,** Design of the endogenous Ref(2)P knock-in allele carrying a C-terminal TEV-3xFlag cassette. **b,** Western blot of control, Ref(2)P-sfGFP knock-in, Ref(2)P-APEX2-Flag knock-in and Ref(2)P-3xFlag knock-in flies. Anti-Flag detects the tagged alleles; beta-actin is the loading control. **c,** Immunofluorescence of muscle, midgut and oenocytes from Ref(2)P-3xFlag knock-in flies stained with anti-Flag (green), anti-Ref(2)P (red) and DAPI (blue). Scale bars, 10 µm.

**Extended Data Fig. 3.**
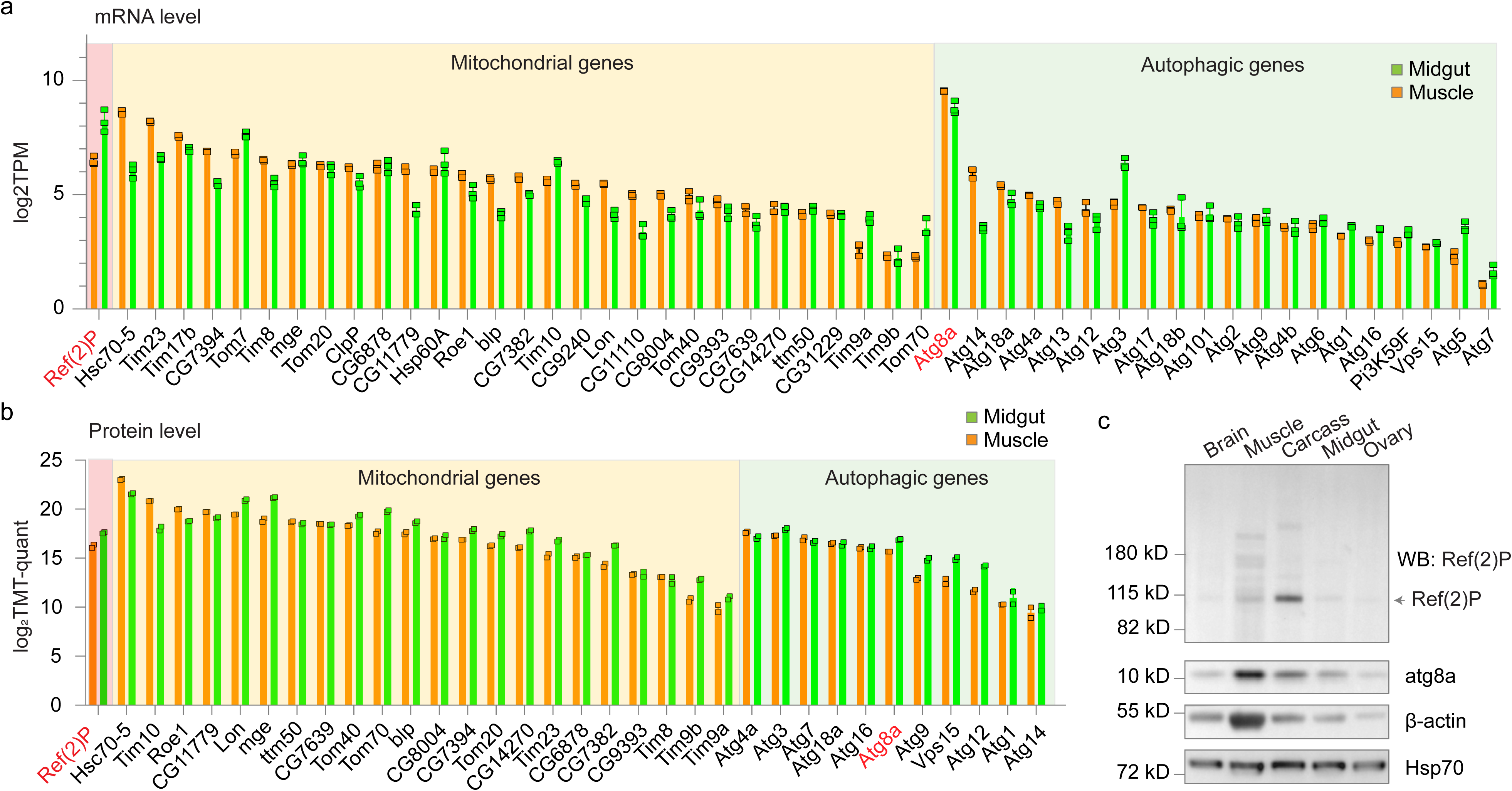
: Autophagy and mitochondrial-quality-control factors are expressed in both midgut and muscle. **a,** RNA abundance of mitochondrial and autophagy genes in midgut (green) and muscle (orange), shown as log2 transcripts per million (TPM). Points denote the three biological replicates. **b,** Quantitative abundance of the corresponding proteins measured by TMT proteomics and shown as log2 TMT quantity. Points denote biological replicates. **c,** Western blot of Ref(2)P and Atg8a across the indicated adult tissues. Beta-actin and Hsp70 are loading controls.

**Extended Data Fig. 4.**
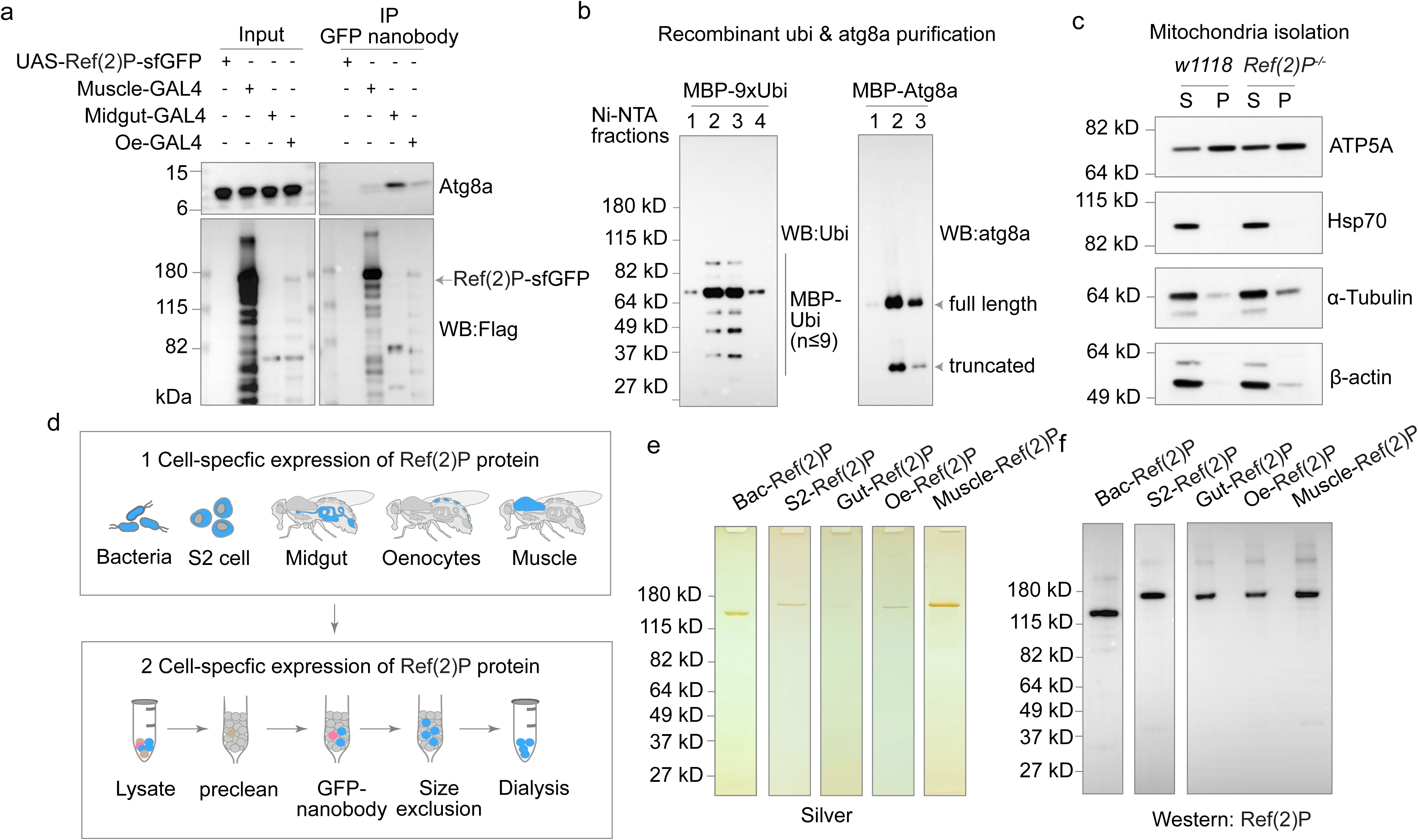
: Purification controls and in vivo validation of tissue-specific Ref(2)P binding. **a,** Co-immunoprecipitation of tissue-expressed Ref(2)P-sfGFP-Flag with endogenous Atg8a. GFP nanobody was used for immunoprecipitation; anti-Atg8a and anti-Flag were used for western blotting. Input, 2%. **b,** Ni-NTA purification fractions of recombinant MBP-9xUbi and MBP-Atg8a analyzed with anti-ubiquitin and anti-Atg8a. Full-length and truncated products are indicated. **c,** Mitochondrial fractionation from control (w1118) and Ref(2)P-null (Ref(2)P^-/-^) flies. ATP5A marks mitochondria; Hsp70, alpha-tubulin and beta-actin monitor cytosolic contamination. S, supernatant; P, mitochondrial pellet. **d,** Workflow for source-specific expression and purification of Ref(2)P-sfGFP-Flag from E. coli, S2 cells, midgut, oenocytes and muscle. **e, f,** Purified Ref(2)P preparations analyzed by silver staining (e) and western blotting with anti-Ref(2)P (f).

**Extended Data Fig. 5.**
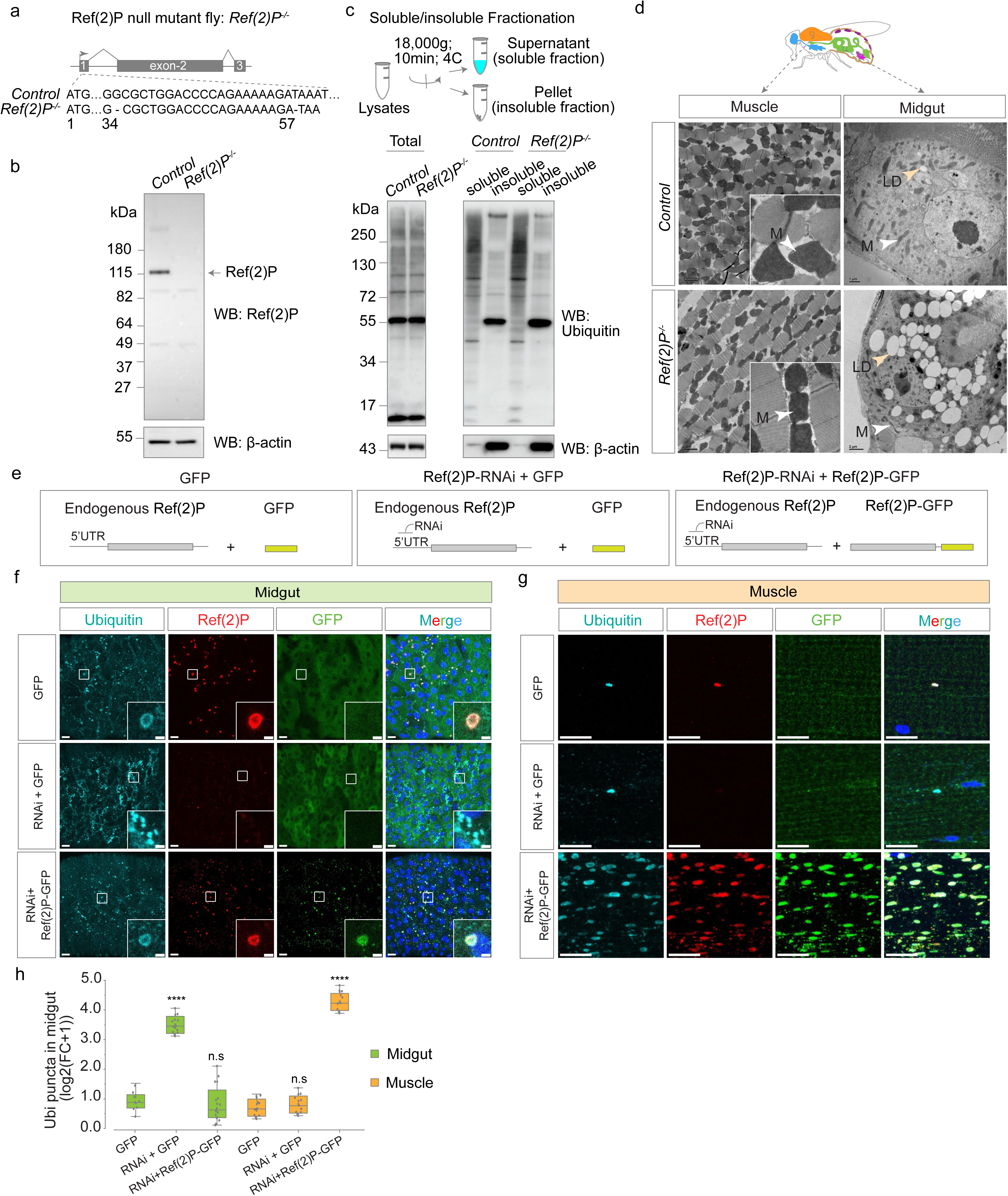
: Validation of Ref(2)P loss of function and tissue-specific rescue. **a,** Genomic lesion in the Ref(2)P^-/-^ null allele. The frameshift introduces a premature stop codon. **b,** Western blot of control and Ref(2)P^-/-^ whole-fly lysates with anti-Ref(2)P; beta-actin is the loading control. **c,** Detergent-soluble and detergent-insoluble fractionation of control and Ref(2)P-null lysates, analyzed with anti-ubiquitin; beta-actin is the loading control. **d,** Electron microscopy of muscle and midgut from control and Ref(2)P-null flies. M, mitochondrion; LD, lipid droplet. Arrowheads mark the indicated organelles. Scale bars, 1 or 2 µm as indicated. **e,** Design of the tissue-specific depletion and rescue experiment. The RNAi targets the 5-prime UTR of endogenous Ref(2)P and therefore does not target the UAS-Ref(2)P-sfGFP rescue transgene. **f, g,** Immunofluorescence of midgut (f) and muscle (g) expressing control GFP, Ref(2)P RNAi, or Ref(2)P RNAi together with Ref(2)P-sfGFP. Ubiquitin (cyan), endogenous Ref(2)P (red), transgenic GFP or Ref(2)P-sfGFP (green) and DAPI (blue). Insets show boxed regions. Scale bars 10 µm; Insets, magnified views of the boxed regions, Scale bars 2 µm. **h,** Quantification of ubiquitin puncta in f and g, expressed as log2(fold change + 1). Each point represents one fly; two-tailed unpaired t-test; ****P < 0.0001; n.s., not significant.

**Extended Data Fig. 6:**
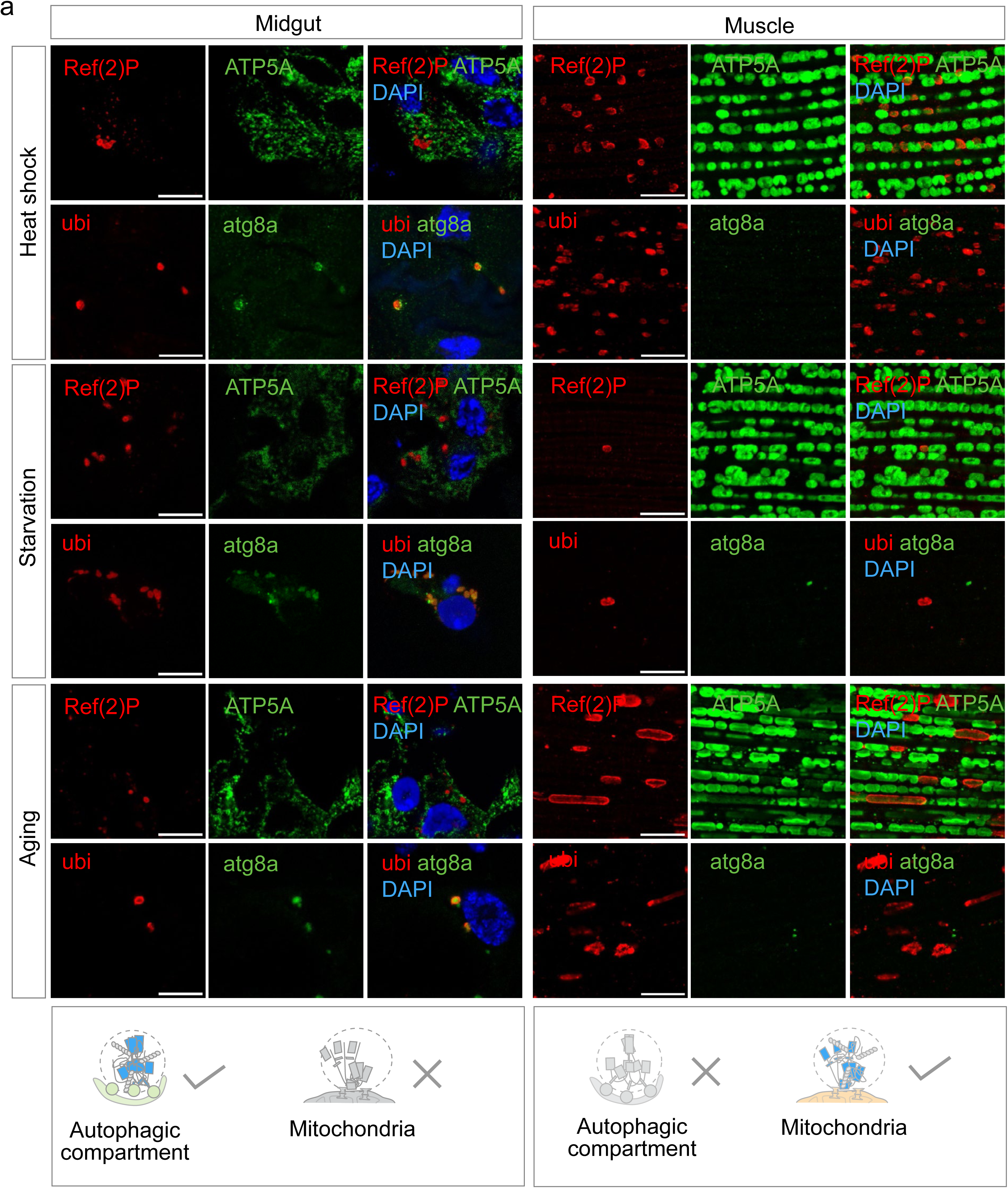
Tissue-specific Ref(2)P localization persists during heat shock, starvation and ageing. **a,** Immunofluorescence of midgut and muscle after heat shock, after 48 h starvation and in 6- week-old flies. The upper row for each tissue shows Ref(2)P (red) with the mitochondrial marker ATP5A (green); the lower row shows ubiquitin (red) with the autophagy marker Atg8a (green). DAPI is blue. The summary schematics indicate that Ref(2)P remains associated with autophagic compartments in midgut and mitochondria in muscle. Scale bars 10 µm; Insets, magnified views of the boxed regions, Scale bars 2 µm.

**Extended Data Fig. 7.**
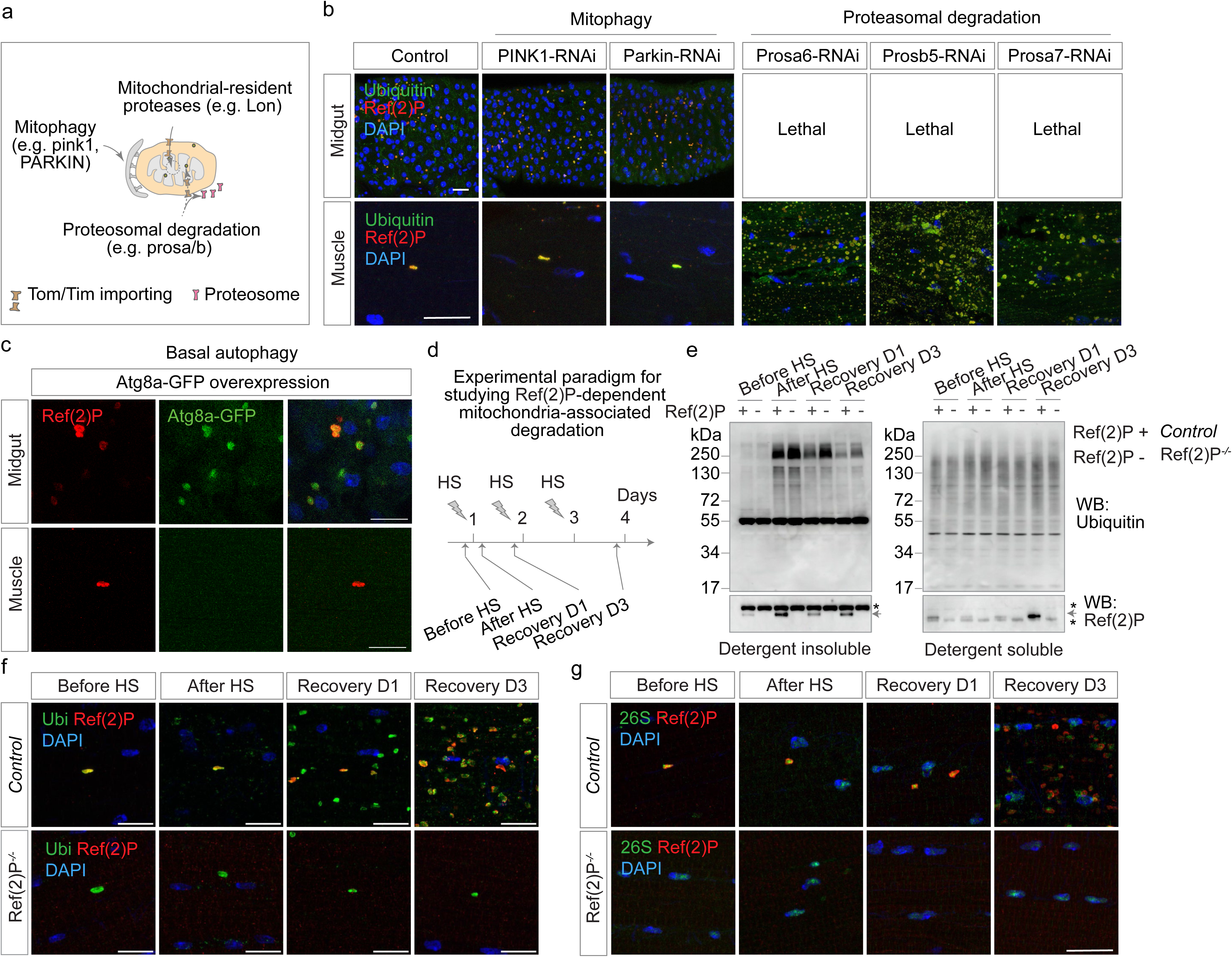
: Genetic and heat-shock tests of mitochondria-associated protein degradation. **a,** The three mitochondrial degradation routes tested: PINK1/Parkin-dependent mitophagy, the mitochondrial-resident protease Lon and proteasomal degradation at the mitochondrial surface. **b,** Immunofluorescence of midgut and muscle after tissue-specific knockdown of PINK1, Parkin, Lon or the indicated proteasome subunits. Ubiquitin (green), Ref(2)P (red) and DAPI (blue). Knockdown of Prosalpha6, Prosbeta5 or Prosalpha7 in midgut was lethal. Scale bars 10 µm.. **c,** Basal Atg8a-GFP signal in midgut and muscle expressing Atg8a-GFP; Ref(2)P is red and DAPI blue. Scale bars 10 µm. **d,** Repeated heat-shock paradigm and sample-collection time points: before heat shock, immediately after heat shock, and after 1 or 3 days of recovery. **e,** Detergent- insoluble and detergent-soluble ubiquitinated proteins from control and Ref(2)P-null muscle across the heat-shock time course. Anti-Ref(2)P identifies Ref(2)P and nonspecific bands marked by asterisks. **f, g,** Immunofluorescence of control and Ref(2)P-null muscle across the heat-shock time course, stained for ubiquitin (f) or the 26S proteasome (g), together with Ref(2)P and DAPI. Scale bars 10 µm.

**Extended Data Fig. 8.**
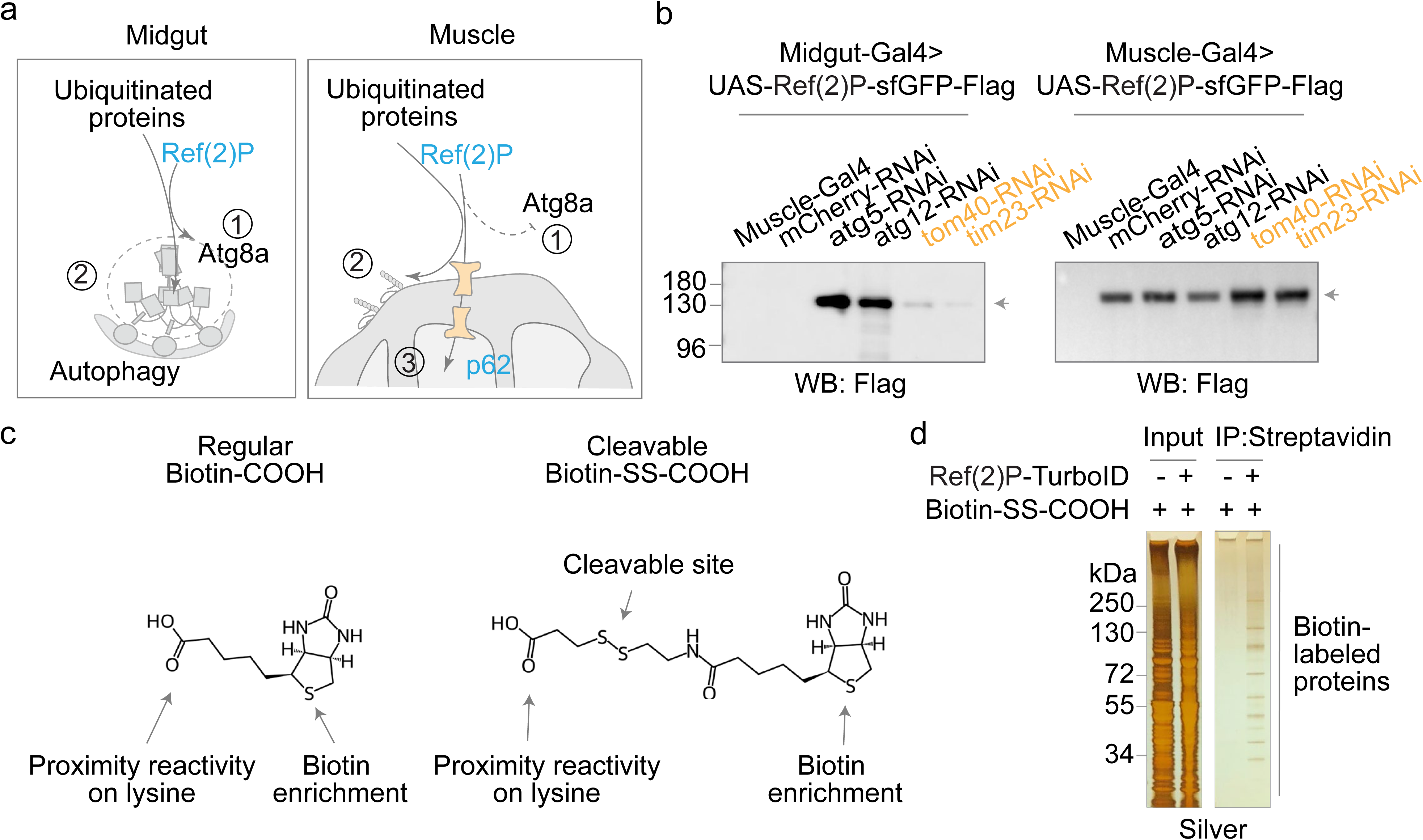
: Mitochondrial-import perturbation and Ref(2)P-TurboID controls. **a,** Working model for tissue-specific routing of ubiquitinated cargo: Atg8a-dependent autophagy in midgut and mitochondrial-import-associated degradation in muscle. **b,** Western blot of UAS- Ref(2)P-sfGFP-Flag in midgut and muscle upon knockdown of the indicated autophagy or mitochondrial-import genes. Anti-Flag detects transgenic Ref(2)P. **c,** Structures and functional groups of conventional biotin-COOHand the cleavable biotin-SS-COOH reagent used for proximity labelling and enrichment. **d,** Silver staining of input and streptavidin-captured proteins from control and Ref(2)P-TurboID muscle supplied with biotin-SS-COOH. The Ref(2)P bait and the labelled protein population are indicated.

**Extended Data Fig. 9.**
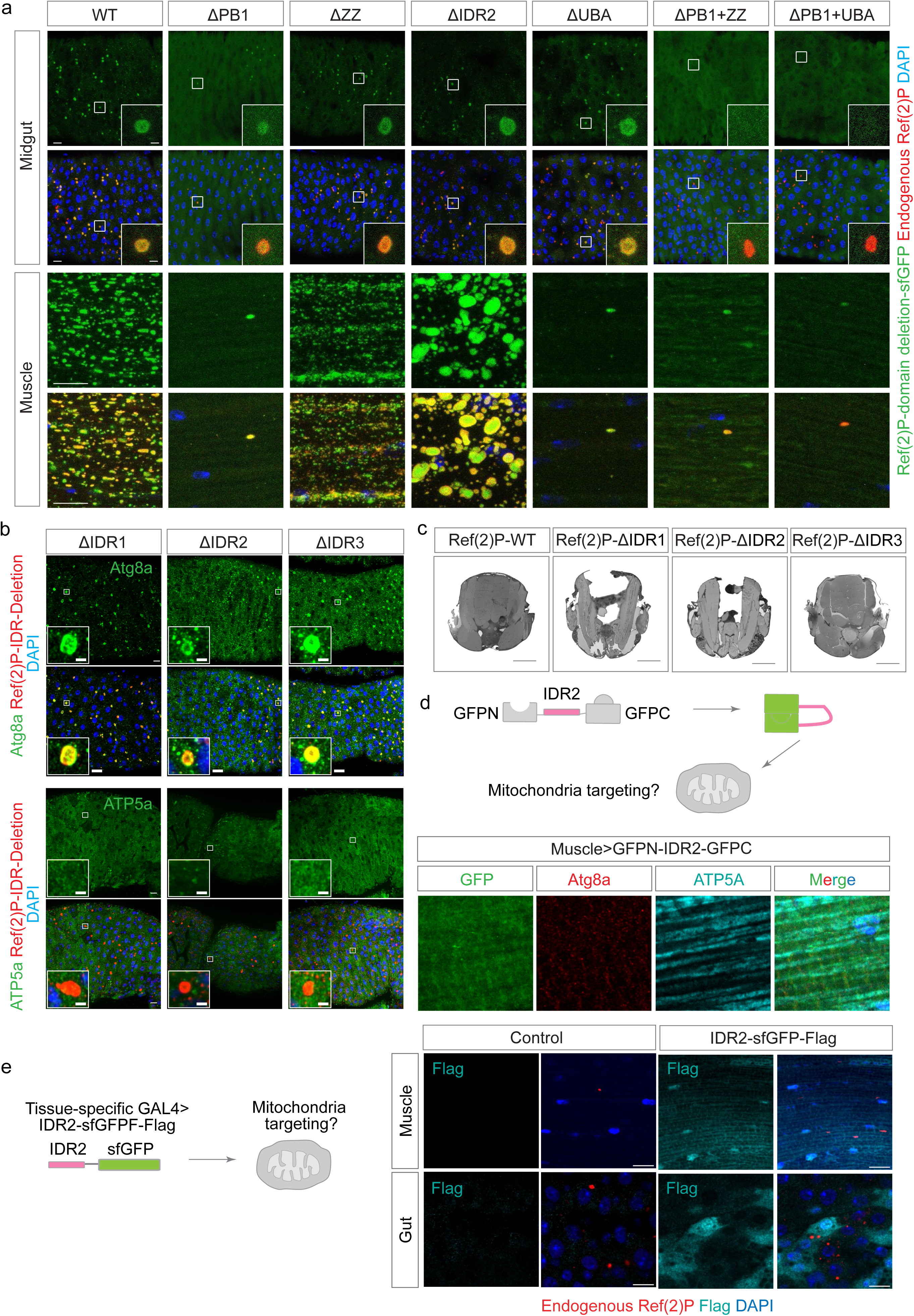
: Domain and intrinsically disordered-region requirements for tissue- specific Ref(2)P localization. **a,** Immunofluorescence of midgut and muscle expressing full-length or the indicated domain- deleted Ref(2)P-sfGFP proteins. Transgenic Ref(2)P-sfGFP is green, endogenous and transgenic Ref(2)P detected with anti-Ref(2)P is red, and DAPI is blue. Insets show boxed regions. Scale bars 10 µm. **b,** Midgut expressing Ref(2)P-ΔIDR1, Ref(2)P-ΔIDR2 or Ref(2)P-ΔIDR3 and stained for Atg8a or ATP5A, transgenic Ref(2)P and DAPI. Scale bars 10 µm. **c,** Sections of 1-week-old flight muscle expressing Ref(2)P-WT or the indicated IDR deletion. Scale bars, 200 µm. **d,** Split-GFP design and fluorescence images of an IDR2-only construct flanked by GFP fragments in muscle. The diffuse signal does not overlap the mitochondrial marker ATP5A. Scale bars 10 µm. **e,** Tissue- specific expression of IDR2-sfGFP-Flag in muscle and gut. Flag is cyan, endogenous Ref(2)P is red and DAPI is blue. Scale bars 10 µm.

**Extended Data Fig. 10.**
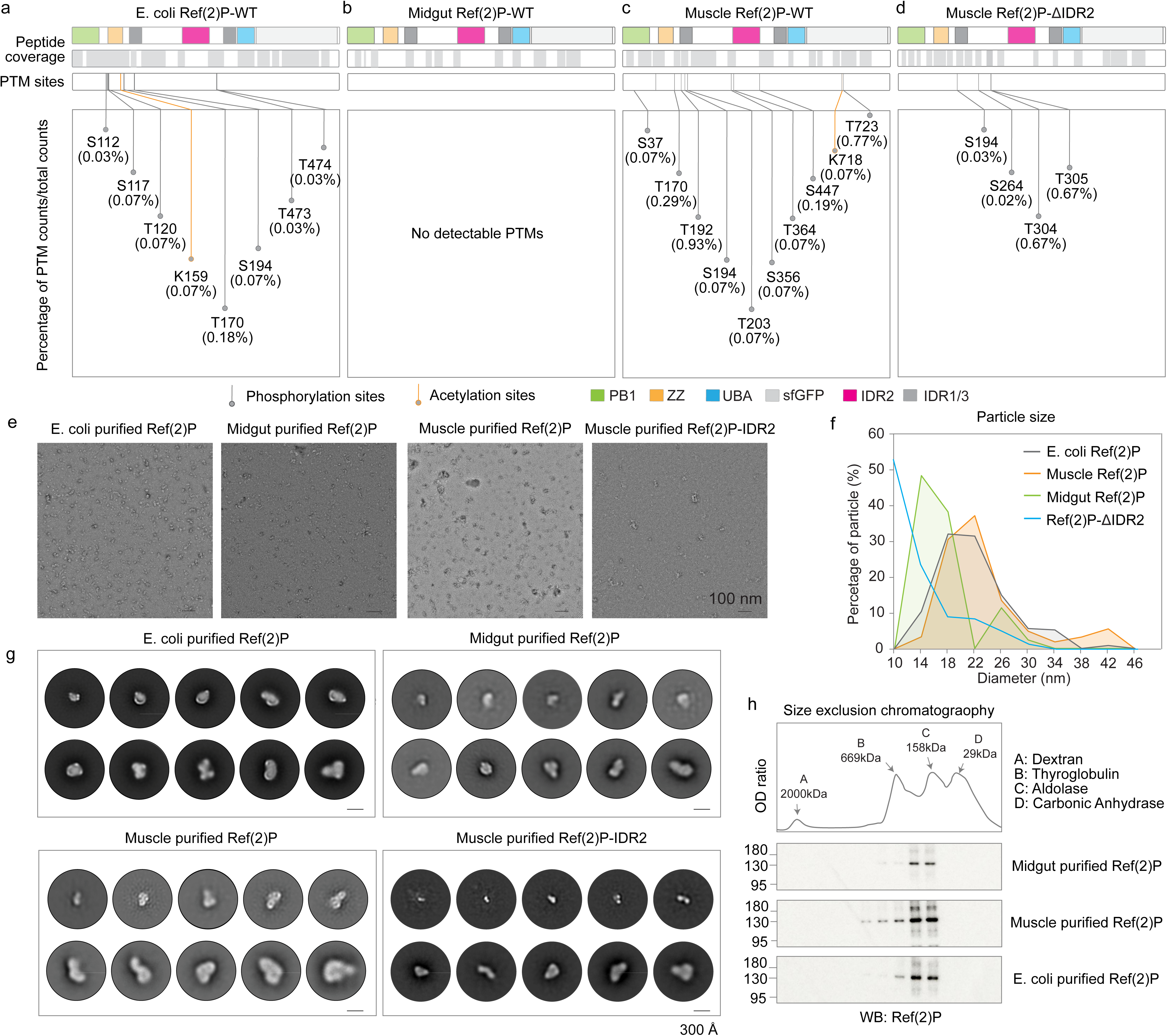
: Post-translational modifications and particle properties of Ref(2)P purified from different sources. **a-d,** Peptide coverage and detected phosphorylation (grey) or acetylation (orange) sites in Ref(2)P purified from E. coli (a), midgut (b), muscle (c) and muscle expressing Ref(2)P-ΔIDR2 (d). Values are modification counts as a percentage of total peptide counts. **e,** Negative-stain electron micrographs of the indicated purified Ref(2)P preparations. Scale bars, 100 nm. **f,** Particle- diameter distributions measured from e. **g,** Representative two-dimensional classes of particles from the indicated preparations. Scale bars, 300 Angstrom. **h,** Size-exclusion chromatography of Ref(2)P purified from midgut, muscle and E. coli. Elution standards are indicated above the chromatogram; fractions were analysed by western blotting with anti-Ref(2)P.

**Extended Data Fig. 11.**
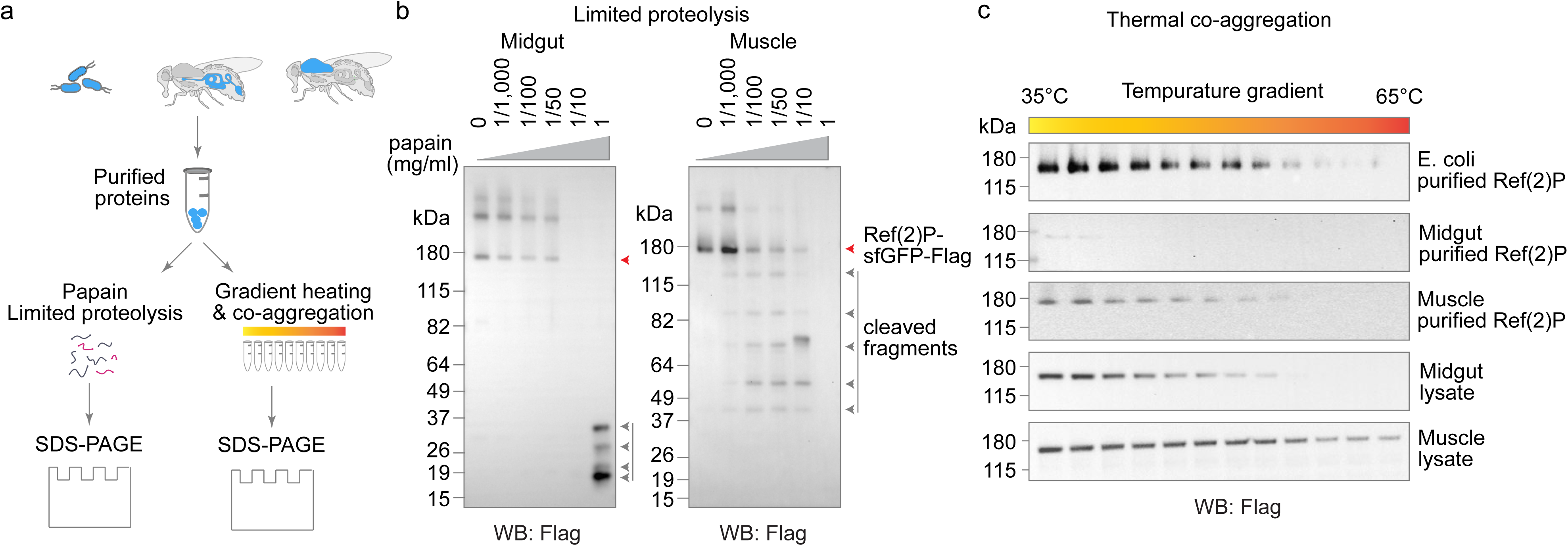
: Limited proteolysis and thermal co-aggregation distinguish midgut and muscle Ref(2)P. **a,** Workflow for limited proteolysis and thermal co-aggregation of source-purified Ref(2)P. **b,** Limited papain digestion of Ref(2)P-sfGFP-Flag purified from midgut and muscle. The C-terminal Flag tag detects full-length protein and C-terminal cleavage products. **c,** Thermal co-aggregation of Ref(2)P purified from E. coli, midgut or muscle, and of Ref(2)P in midgut or muscle lysates, across a 35-65 degrees C gradient. Anti-Flag detects Ref(2)P-sfGFP-Flag.

**Extended Data Fig. 12.**
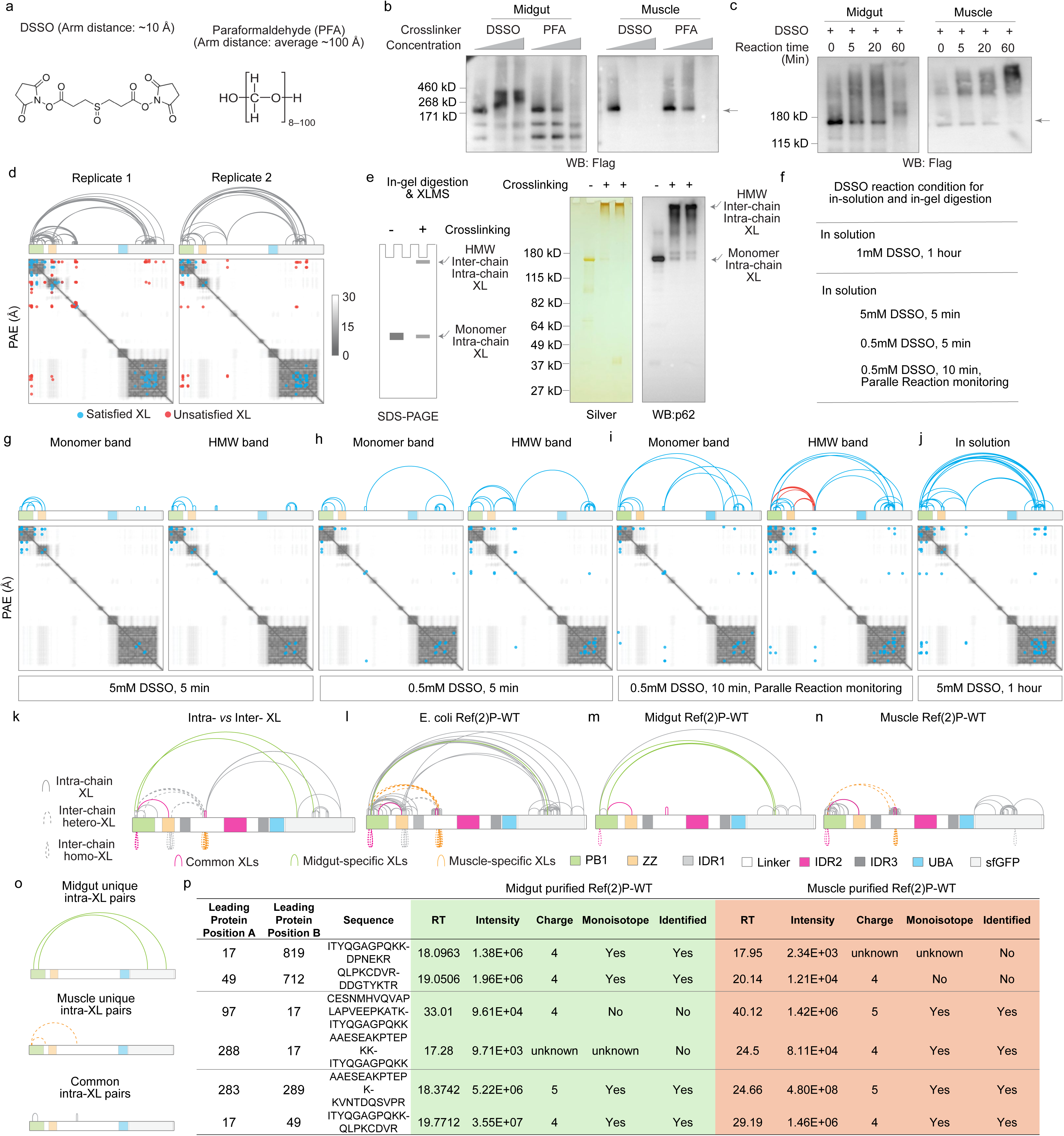
: Crosslinking controls, reproducibility and assignment of intra-chain and inter-chain Ref(2)P crosslinks. **a,** Chemical structures and approximate spacer lengths of DSSO and paraformaldehyde (PFA). **b,** Western blot of midgut- and muscle-derived Ref(2)P after increasing DSSO or PFA concentrations. Anti-Flag detects monomeric and high-molecular-weight crosslinked species. **c,** DSSO time course for midgut- and muscle-derived Ref(2)P. **d,** Crosslink arc diagrams and PAE maps from two independent mass-spectrometry acquisitions of the same crosslinked sample. Satisfied and unsatisfied crosslinks are blue and red, respectively. **e,** In-gel workflow used to distinguish crosslinks recovered from the monomer band from those recovered in high-molecular-weight material, with silver stain and anti-Ref(2)P western blot of the excised regions. **f,** DSSO reaction and acquisition conditions used for in-solution and in-gel analyses. **g-j,** Crosslink maps for monomer and high-molecular-weight bands under the indicated reaction and acquisition conditions (g-i), and for the in-solution digest (j). Orange arcs mark pairs unique to high- molecular-weight material. **k,** Classification of intra-chain, inter-chain heterotypic and inter-chain homotypic crosslinks. **l-n,** Crosslink maps for Ref(2)P purified from E. coli (l), midgut (m) and muscle (n). Magenta, common; green, midgut-specific; orange, muscle-specific. **o,** Representative midgut-specific, muscle-specific and common intra-chain pairs. **p,** Precursor features for representative common and tissue-specific crosslinked peptides in the midgut and muscle datasets.

**Extended Data Fig. 13.**
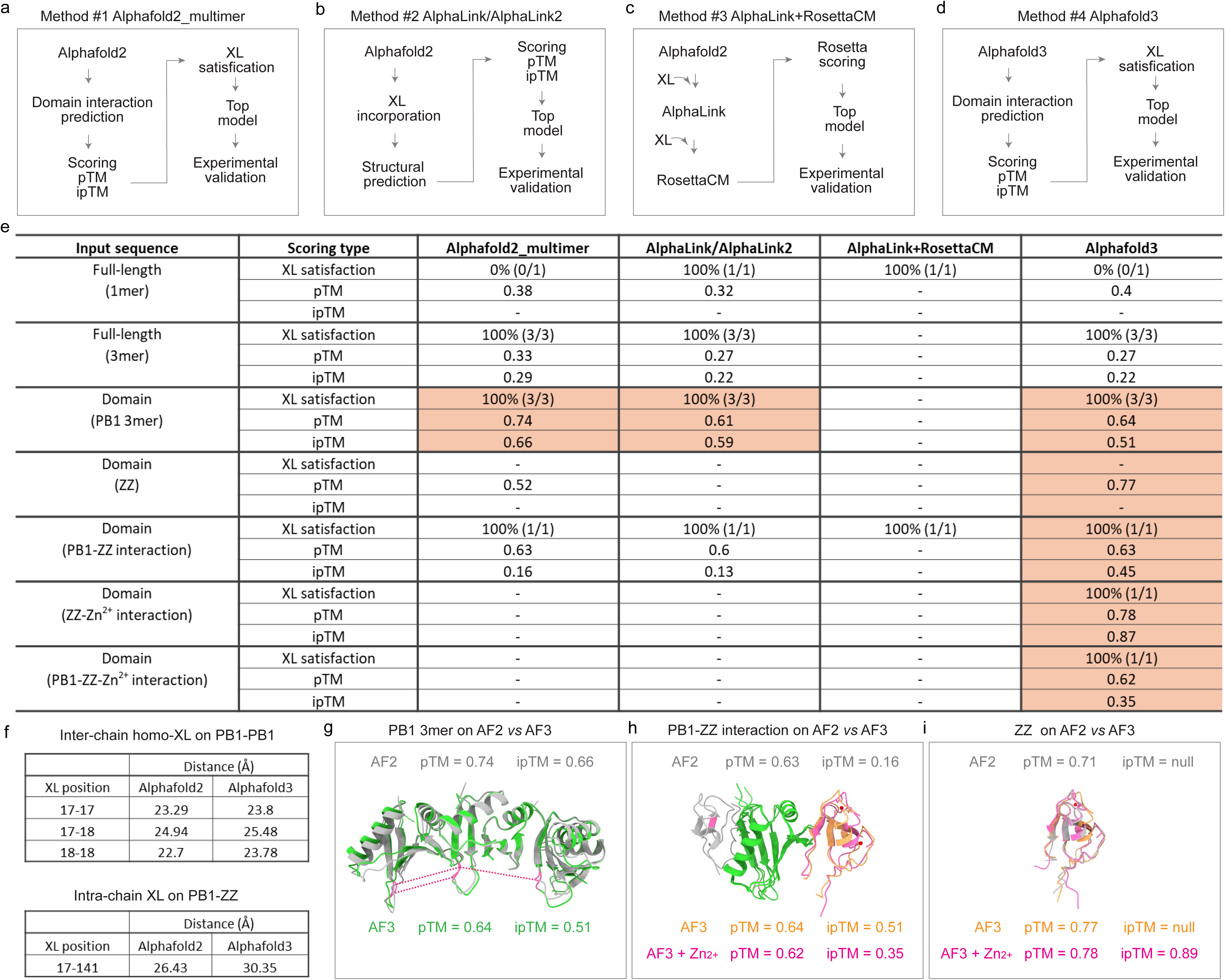
: Comparison of crosslink-guided structural-modelling strategies. **a-d,** Workflows based on AlphaFold2 multimer (a), AlphaLink or AlphaLink2 (b), AlphaLink plus RosettaCM (c) and AlphaFold3 (d). **e,** Comparison of crosslink satisfaction, pTM and ipTM for the four methods and the indicated sequence inputs. A dash denotes a metric that was not available. **f,** Distances of inter-chain PB1-PB1 and intra-chain PB1-ZZ crosslinks in AlphaFold2 and AlphaFold3 models. **g-i,** Comparison of AlphaFold2 and AlphaFold3 models of the PB1 trimer (g), the PB1-ZZ interaction (h) and the ZZ domain with or without zinc (i).

**Extended Data Fig. 14.**
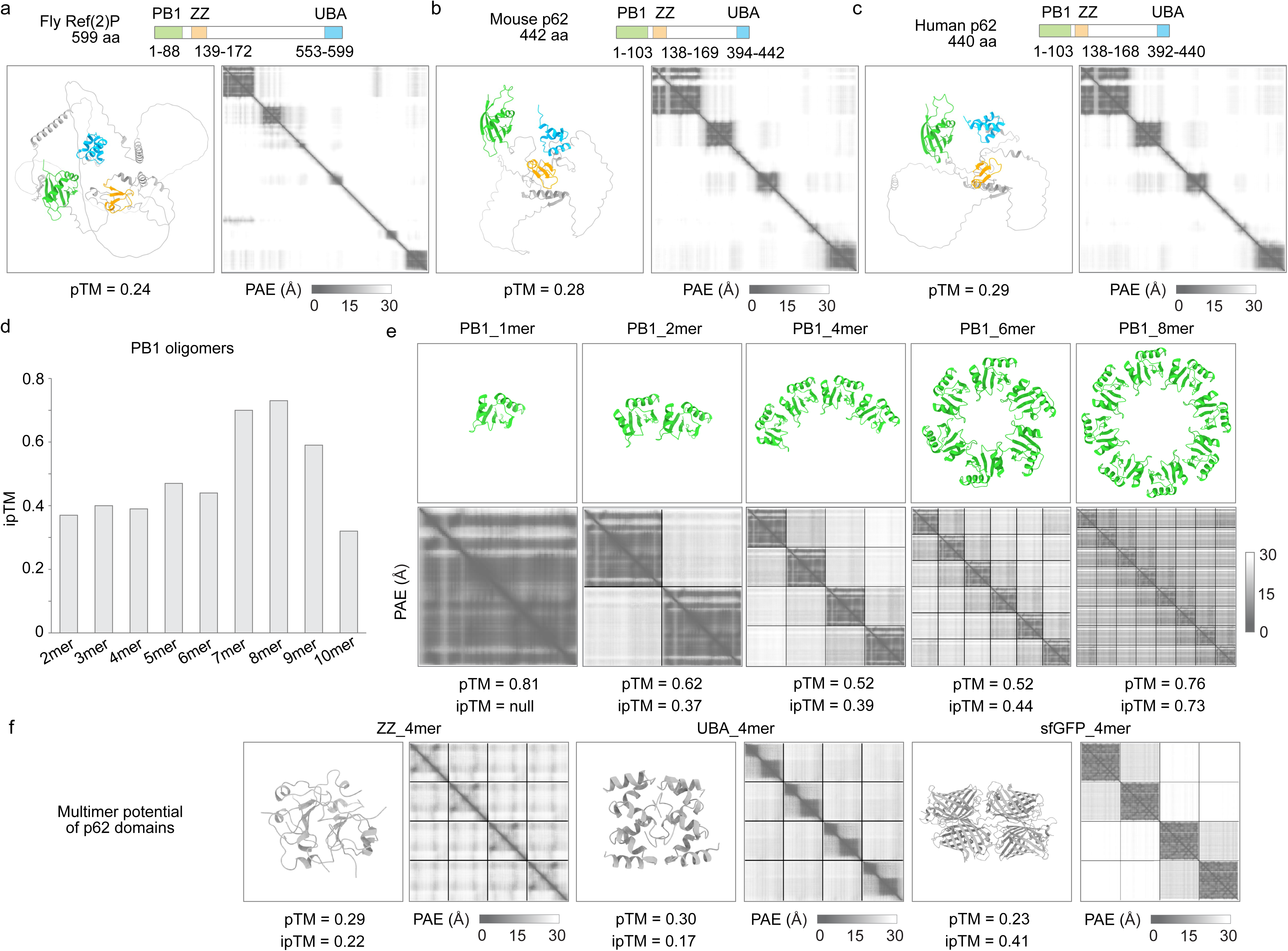
: AlphaFold3 predictions of full-length Ref(2)P and isolated domains. **a-c,** AlphaFold3 full-length models and PAE maps for Drosophila (a), mouse (b) and human (c) p62. Domain boundaries and pTM scores are indicated. **d,** ipTM scores for predicted Drosophila PB1 oligomers containing two to ten subunits. **e,** Structural models and PAE maps of PB1 monomer, dimer, tetramer, hexamer and octamer, with pTM and ipTM values. **f,** Predicted tetramers and PAE maps for the ZZ domain, UBA domain and sfGFP.

**Extended Data Fig. 15.**
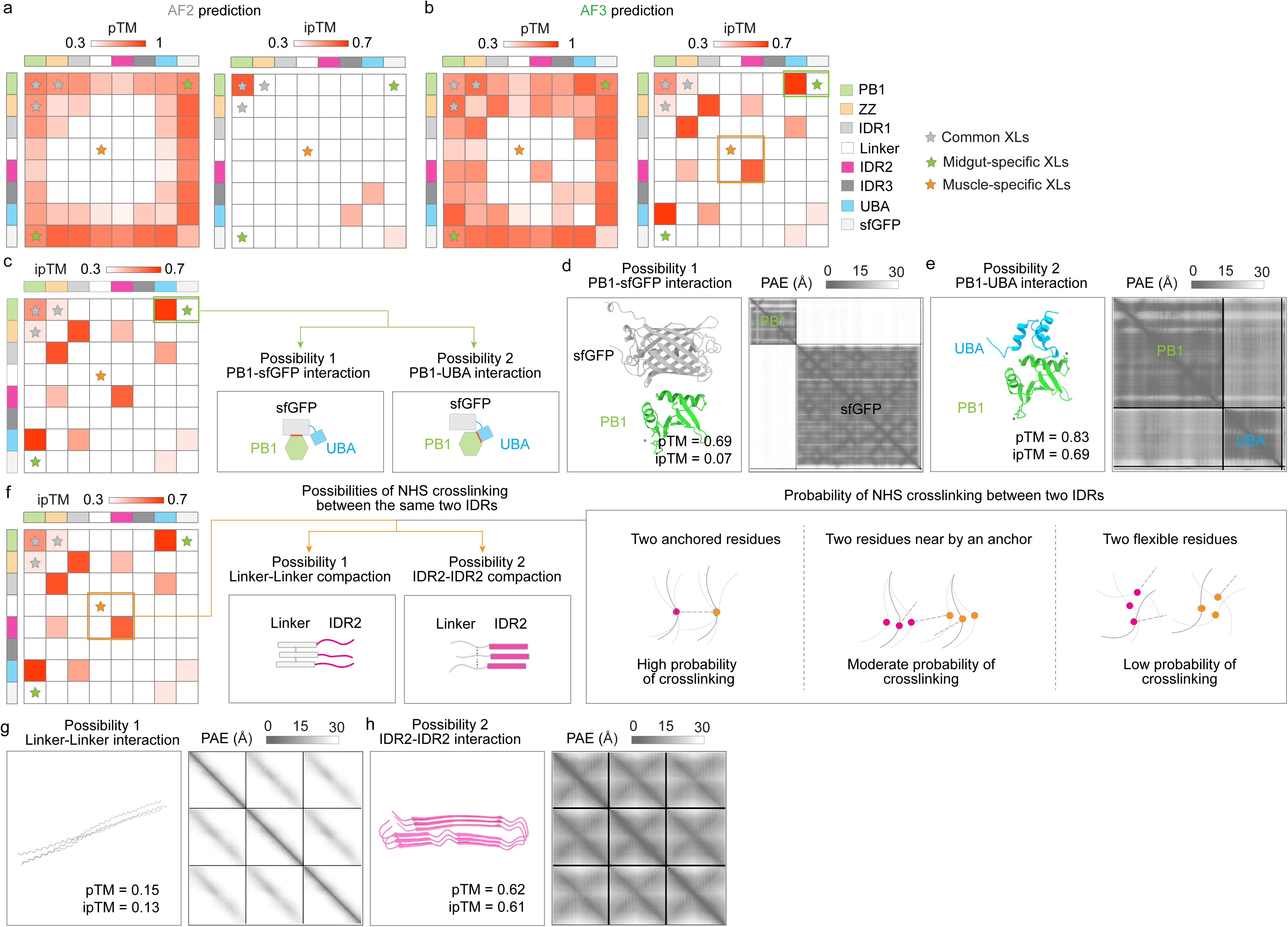
: Crosslink-guided selection of midgut- and muscle-specific structural models. **a, b,** AlphaFold2 (a) and AlphaFold3 (b) pTM and ipTM domain-interaction matrices. Stars mark common (grey), midgut-specific (green) and muscle-specific (orange) crosslinks. **c,** Two explanations for the midgut-specific PB1-to-C-terminal crosslink: PB1-sfGFP or PB1-UBA interaction. **d, e,** AlphaFold3 models and PAE maps for PB1-sfGFP (d) and PB1-UBA (e). **f,** Two explanations for the muscle-specific crosslinks between equivalent linker regions: linker-linker compaction or interaction between the adjacent IDR2 regions. The right-hand schematic summarizes how residue constraint affects the probability of NHS crosslinking. **g, h,** AlphaFold3 models and PAE maps for linker-linker (g) and IDR2-IDR2 (h) interactions.

**Extended Data Fig. 16.**
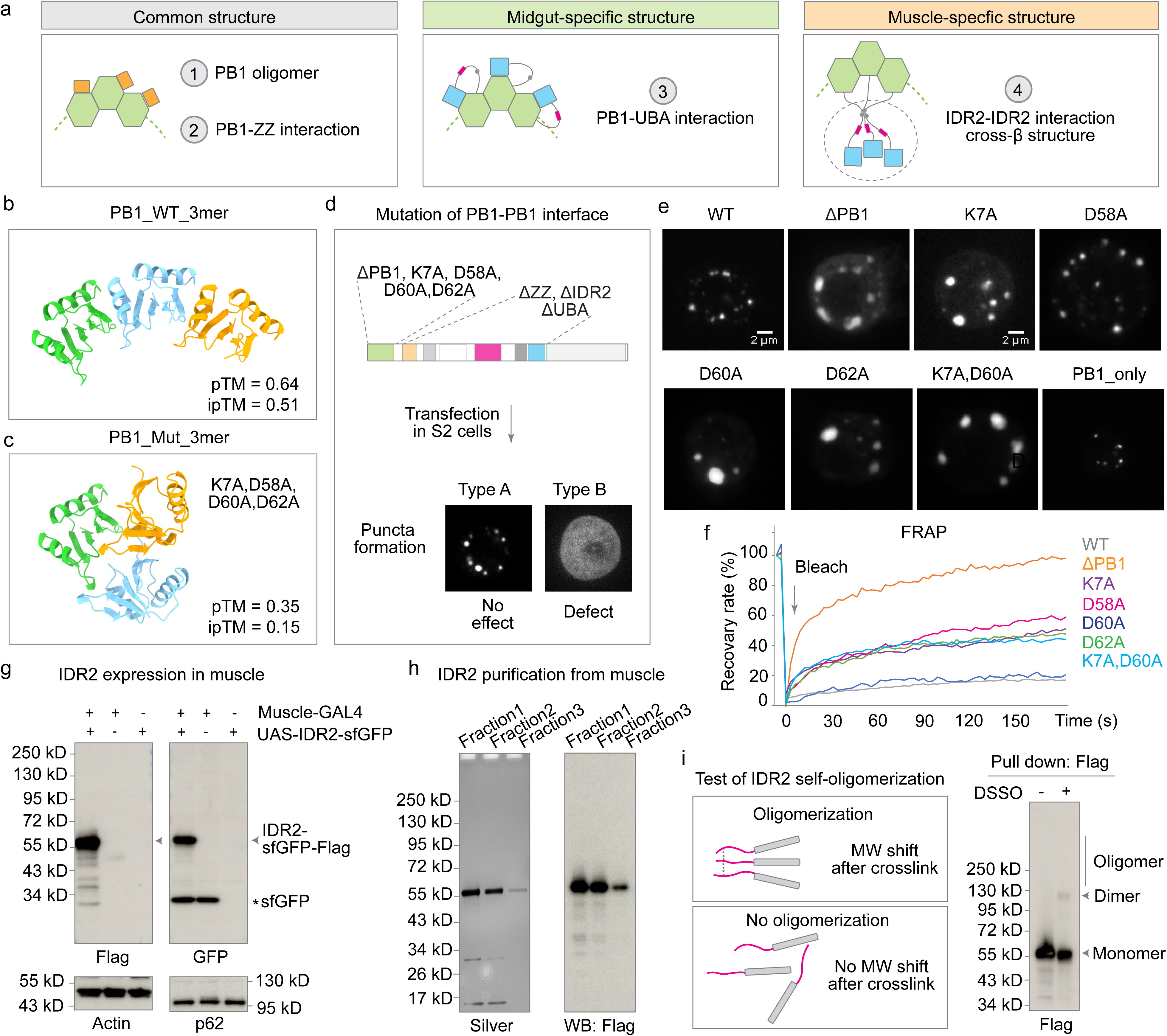
: Experimental validation of common and muscle-specific Ref(2)P structural features. **a,** Summary of the structural features tested: common PB1 oligomerization and PB1-ZZ interaction, the midgut-specific PB1-UBA interaction and the muscle-specific IDR2-IDR2 cross- beta interaction. **b, c,** AlphaFold3 models of wild-type PB1 trimer (b) and a PB1 trimer carrying K7A, D58A, D60A and D62A (c). **d,** Design of the S2-cell assay used to score punctate versus diffuse Ref(2)P and to test mutations in the predicted PB1-PB1 interface. **e, Fluorescent images of** representative S2 cells expressing wild-type Ref(2)P, Ref(2)P-ΔPB1, the indicated interface mutants or the PB1 domain alone. Scale bars, 2 µm. **f,** Fluorescence recovery after photobleaching of wild-type and mutant Ref(2)P-sfGFP puncta. The arrow marks bleaching. **g,** Expression of IDR2-sfGFP-Flag in muscle analysed with anti-Flag and anti-GFP; actin is the loading control and endogenous Ref(2)P is shown below. **h,** Silver stain and anti-Flag western blot of fractions from purification of muscle-derived IDR2-sfGFP-Flag. **i,** DSSO crosslinking test for IDR2 self-association. Anti-Flag detects monomeric, dimeric and higher-order species.

**Extended Data Fig. 17.**
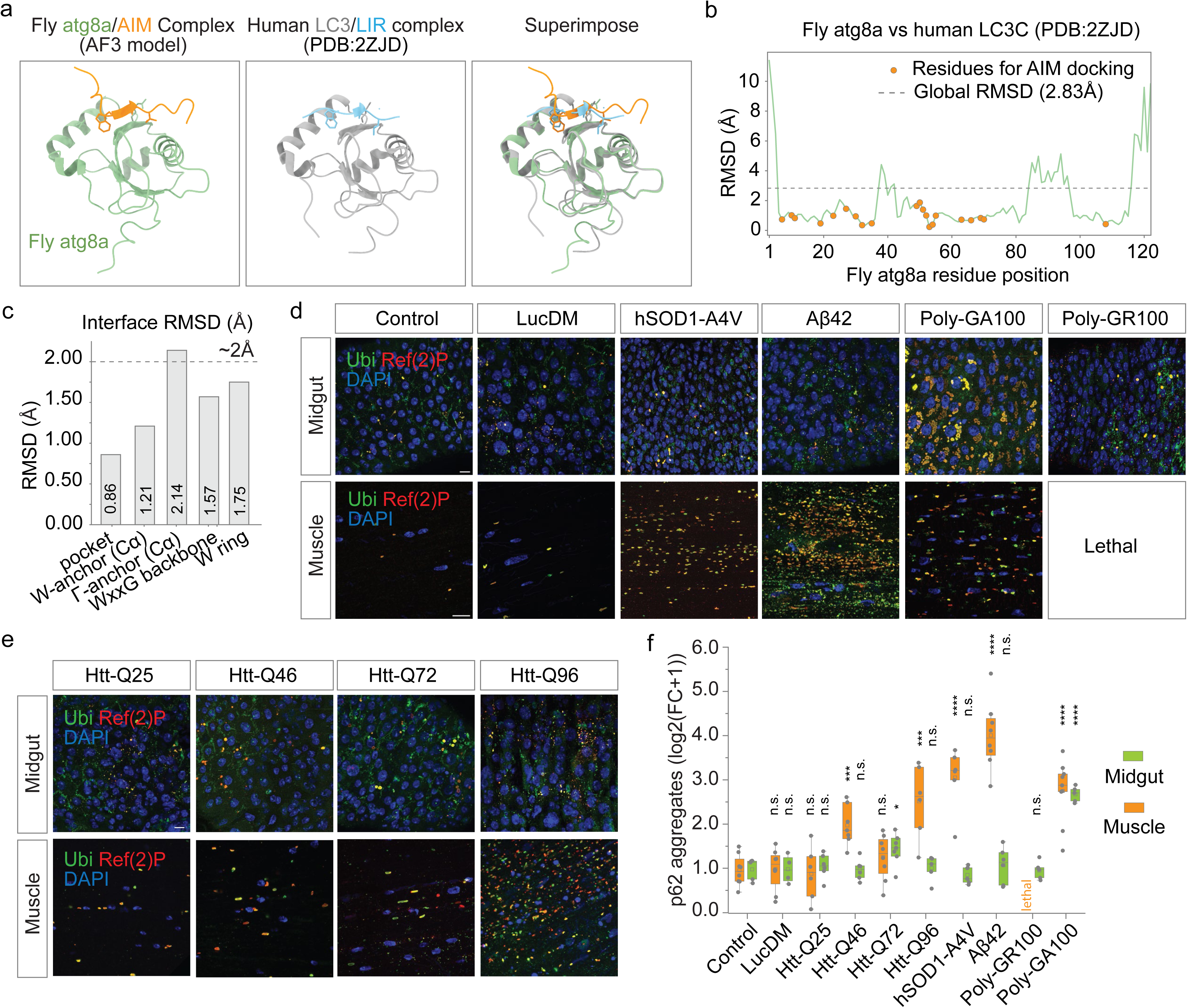
: Conservation of the Atg8a-binding interface and tissue-specific responses to aggregation-prone proteins. **a,** AlphaFold3 model of the Drosophila Atg8a-AIM complex, the human LC3-LIR structure (PDB 2ZJD) and their superposition. **b,** Per-residue RMSD between fly Atg8a and human LC3C. Orange points mark residues that contact the AIM; the dashed line marks the global RMSD of 2.83 Angstrom. **c,** RMSD values for structural elements of the AIM/LIR-binding interface. **d,** Immunofluorescence of midgut and muscle expressing the indicated aggregation-prone proteins. Ubiquitin is green, Ref(2)P red and DAPI blue. Muscle expression of poly-GR100 was lethal. Scale bars, 10 µm. **e,** Immunofluorescence of midgut and muscle expressing the indicated Htt polyglutamine-length variants. Ubiquitin is green, Ref(2)P red and DAPI blue. Scale bars, 10 µm. **f,** Quantification of Ref(2)P aggregates in d and e, expressed as log2(fold change + 1). Each point represents one fly; two-tailed unpaired t-test; **P < 0.01, ****P < 0.0001; n.s., not significant.

## References

1 Baralle, F. E. & Giudice, J. Alternative splicing as a regulator of development and tissue identity. Nature reviews. Molecular cell biology 18, 437–451 (2017).

2 Dishman, A. F. & Volkman, B. F. Metamorphic protein folding as evolutionary adaptation. Trends in biochemical sciences 48, 665–672 (2023).

3 Kim, A. K. & Porter, L. L. Functional and Regulatory Roles of Fold-Switching Proteins. Structure 29, 6–14 (2021).

4 Porter, L. L. & Looger, L. L. Extant fold-switching proteins are widespread. Proceedings of the National Academy of Sciences of the United States of America 115, 5968–5973 (2018).

5 Jarosz, D. F. & Khurana, V. Specification of Physiologic and Disease States by Distinct Proteins and Protein Conformations. Cell 171, 1001–1014 (2017).

6 Ruan, L. et al. Mitochondria-Associated Proteostasis. Annual review of biophysics 49, 41–67 (2020).

7 Hipp, M. S., Kasturi, P. & Hartl, F. U. The proteostasis network and its decline in ageing. Nature reviews. Molecular cell biology 20, 421–435 (2019).

8 Glickman, M. H. & Ciechanover, A. The ubiquitin-proteasome proteolytic pathway: destruction for the sake of construction. Physiological reviews 82, 373–428 (2002).

9 Shaid, S., Brandts, C. H., Serve, H. & Dikic, I. Ubiquitination and selective autophagy. Cell death and differentiation 20, 21–30 (2013).

10 Dikic, I., Wakatsuki, S. & Walters, K. J. Ubiquitin-binding domains - from structures to functions. Nature reviews. Molecular cell biology 10, 659–671 (2009).

11 Vargas, J. N. S., Hamasaki, M., Kawabata, T., Youle, R. J. & Yoshimori, T. The mechanisms and roles of selective autophagy in mammals. Nature reviews. Molecular cell biology 24, 167–185 (2023).

12 Lamark, T. & Johansen, T. Mechanisms of Selective Autophagy. Annual review of cell and developmental biology 37, 143–169 (2021).

13 Lamark, T. et al. Interaction codes within the family of mammalian Phox and Bem1p domain- containing proteins. The Journal of biological chemistry 278, 34568–34581 (2003).

14 Horos, R. et al. The Small Non-coding Vault RNA1-1 Acts as a Riboregulator of Autophagy. Cell 176, 1054–1067 e1012 (2019).

15 Birgisdottir, A. B., Lamark, T. & Johansen, T. The LIR motif - crucial for selective autophagy. Journal of cell science 126, 3237–3247 (2013).

16 Isogai, S. et al. Crystal structure of the ubiquitin-associated (UBA) domain of p62 and its interaction with ubiquitin. The Journal of biological chemistry 286, 31864–31874 (2011).

17 Pankiv, S. et al. p62/SQSTM1 binds directly to Atg8/LC3 to facilitate degradation of ubiquitinated protein aggregates by autophagy. The Journal of biological chemistry 282, 24131–24145 (2007).

18 Bjorkoy, G. et al. p62/SQSTM1 forms protein aggregates degraded by autophagy and has a protective effect on huntingtin-induced cell death. The Journal of cell biology 171, 603–614 (2005).

19 Ciuffa, R. et al. The selective autophagy receptor p62 forms a flexible filamentous helical scaffold. Cell reports 11, 748–758 (2015).

20 Zaffagnini, G. et al. p62 filaments capture and present ubiquitinated cargos for autophagy. The EMBO journal 37, e98308 (2018).

21 Dezelee, S. et al. Molecular analysis of ref(2)P, a Drosophila gene implicated in sigma rhabdovirus multiplication and necessary for male fertility. The EMBO journal 8, 3437–3446 (1989).

22 Nezis, I. P. et al. Ref(2)P, the Drosophila melanogaster homologue of mammalian p62, is required for the formation of protein aggregates in adult brain. The Journal of cell biology 180, 1065–1071 (2008).

23 Bartlett, B. J. et al. p62, Ref(2)P and ubiquitinated proteins are conserved markers of neuronal aging, aggregate formation and progressive autophagic defects. Autophagy 7, 572–583 (2011).

24 Nagy, P. et al. Atg17/FIP200 localizes to perilysosomal Ref(2)P aggregates and promotes autophagy by activation of Atg1 in Drosophila. Autophagy 10, 453–467 (2014).

25 DeVorkin, L. & Gorski, S. M. Monitoring autophagic flux using Ref(2)P, the Drosophila p62 ortholog. Cold Spring Harbor protocols 2014, 959–966 (2014).

26 Denton, D. et al. Autophagy, not apoptosis, is essential for midgut cell death in Drosophila. Current biology 19, 1741–1746 (2009).

27 Komatsu, M. et al. The selective autophagy substrate p62 activates the stress responsive transcription factor Nrf2 through inactivation of Keap1. Nature cell biology 12, 213–223 (2010).

28 Duran, A. et al. p62 is a key regulator of nutrient sensing in the mTORC1 pathway. Molecular cell 44, 134–146 (2011).

29 Singh, R. et al. Autophagy regulates lipid metabolism. Nature 458, 1131–1135 (2009).

30 Scott, R. C., Schuldiner, O. & Neufeld, T. P. Role and regulation of starvation-induced autophagy in the Drosophila fat body. Developmental cell 7, 167–178 (2004).

31 Kuma, A. et al. The role of autophagy during the early neonatal starvation period. Nature 432, 1032–1036 (2004).

32 Kauppila, T. E. S., Kauppila, J. H. K. & Larsson, N. G. Mammalian Mitochondria and Aging: An Update. Cell metabolism 25, 57–71 (2017).

33 Ruan, L. et al. Cytosolic proteostasis through importing of misfolded proteins into mitochondria. Nature 543, 443–446 (2017).

34 Vives-Bauza, C. et al. PINK1-dependent recruitment of Parkin to mitochondria in mitophagy. Proceedings of the National Academy of Sciences of the United States of America 107, 378–383 (2010).

35 Narendra, D., Tanaka, A., Suen, D. F. & Youle, R. J. Parkin is recruited selectively to impaired mitochondria and promotes their autophagy. The Journal of cell biology 183, 795–803 (2008).

36 Youle, R. J. & Narendra, D. P. Mechanisms of mitophagy. *Nature reviews.* *Molecular cell biology* **12**, 9–14 (2011).

37 Quiros, P. M., Langer, T. & Lopez-Otin, C. New roles for mitochondrial proteases in health, ageing and disease. Nature reviews. Molecular cell biology 16, 345–359 (2015).

38 Ng, M. Y. W., Wai, T. & Simonsen, A. Quality control of the mitochondrion. Developmental cell 56, 881–905 (2021).

39 Komatsu, M. et al. Homeostatic levels of p62 control cytoplasmic inclusion body formation in autophagy-deficient mice. Cell 131, 1149–1163 (2007).

40 Branon, T. C. et al. Efficient proximity labeling in living cells and organisms with TurboID. Nature biotechnology 36, 880–887 (2018).

41 Itakura, E. & Mizushima, N. p62 Targeting to the autophagosome formation site requires self- oligomerization but not LC3 binding. The Journal of cell biology 192, 17–27 (2011).

42 Wurzer, B. et al. Oligomerization of p62 allows for selection of ubiquitinated cargo and isolation membrane during selective autophagy. eLife 4, e08941 (2015).

43 Wright, P. E. & Dyson, H. J. Intrinsically disordered proteins in cellular signalling and regulation. Nature reviews. Molecular cell biology 16, 18–29 (2015).

44 van der Lee, R. et al. Classification of intrinsically disordered regions and proteins. Chemical reviews 114, 6589–6631 (2014).

45 Babu, M. M. The contribution of intrinsically disordered regions to protein function, cellular complexity, and human disease. Biochemical Society transactions 44, 1185–1200 (2016).

46 Jaffe, E. K. Morpheeins--a new structural paradigm for allosteric regulation. Trends in biochemical sciences 30, 490–497 (2005).

47 Fontana, A., Polverino de Laureto, P., De Filippis, V., Scaramella, E. & Zambonin, M. Probing the partly folded states of proteins by limited proteolysis. Folding & design 2, R17–26 (1997).

48 Fontana, A. et al. Probing protein structure by limited proteolysis. Acta biochimica Polonica 51, 299–321 (2004).

49 Tan, C. S. H. et al. Thermal proximity coaggregation for system-wide profiling of protein complex dynamics in cells. Science 359, 1170–1177 (2018).

50 Berkamp, S. et al. Structural organization of p62 filaments and the cellular ultrastructure of calcium-rich p62-enwrapped lipid droplet cargo. Nature communications 16, 10810 (2025).

51 Jakobi, A. J. et al. Structural basis of p62/SQSTM1 helical filaments and their role in cellular cargo uptake. Nature communications 11, 440 (2020).

52 O’Reilly, F. J. & Rappsilber, J. Cross-linking mass spectrometry: methods and applications in structural, molecular and systems biology. Nature structural & molecular biology 25, 1000–1008 (2018).

53 Piersimoni, L., Kastritis, P. L., Arlt, C. & Sinz, A. Cross-Linking Mass Spectrometry for Investigating Protein Conformations and Protein-Protein Interactions horizontal line A Method for All Seasons. Chemical reviews 122, 7500–7531 (2022).

54 Sinz, A. Chemical cross-linking and mass spectrometry for mapping three-dimensional structures of proteins and protein complexes. Journal of mass spectrometry : JMS 38, 1225–1237 (2003).

55 Liu, F., Lossl, P., Scheltema, R., Viner, R. & Heck, A. J. R. Optimized fragmentation schemes and data analysis strategies for proteome-wide cross-link identification. Nature communications 8, 15473 (2017).

56 Kahraman, A. et al. Cross-link guided molecular modeling with ROSETTA. PloS one 8, e73411 (2013).

57 Stahl, K., Graziadei, A., Dau, T., Brock, O. & Rappsilber, J. Protein structure prediction with in-cell photo-crosslinking mass spectrometry and deep learning. Nature biotechnology 41, 1810–1819 (2023).

58 Manalastas-Cantos, K. et al. Modeling Flexible Protein Structure With AlphaFold2 and Crosslinking Mass Spectrometry. Molecular & cellular proteomics : MCP 23, 100724 (2024).

59 Kao, A. et al. Development of a novel cross-linking strategy for fast and accurate identification of cross-linked peptides of protein complexes. Molecular & cellular proteomics : MCP 10, M110 002212 (2011).

60 Leitner, A. et al. Chemical cross-linking/mass spectrometry targeting acidic residues in proteins and protein complexes. Proceedings of the National Academy of Sciences of the United States of America 111, 9455–9460 (2014).

61 Ichimura, Y. et al. Structural basis for sorting mechanism of p62 in selective autophagy. The Journal of biological chemistry 283, 22847–22857 (2008).

62 Schlagowski, A. M. et al. Increased levels of mitochondrial import factor Mia40 prevent the aggregation of polyQ proteins in the cytosol. The EMBO journal 40, e107913 (2021).

63 Lu, K., Psakhye, I. & Jentsch, S. Autophagic clearance of polyQ proteins mediated by ubiquitin-Atg8 adaptors of the conserved CUET protein family. Cell 158, 549–563 (2014).

64 Mathew, R. et al. Functional role of autophagy-mediated proteome remodeling in cell survival signaling and innate immunity. Molecular cell 55, 916–930 (2014).

65 Lopes da Fonseca, T., Villar-Pique, A. & Outeiro, T. F. The Interplay between Alpha-Synuclein Clearance and Spreading. Biomolecules 5, 435–471 (2015).

66 Ingolia, N. T. Ribosome profiling: new views of translation, from single codons to genome scale. Nature reviews. Genetics 15, 205–213 (2014).

67 Dunn, J. G., Foo, C. K., Belletier, N. G., Gavis, E. R. & Weissman, J. S. Ribosome profiling reveals pervasive and regulated stop codon readthrough in Drosophila melanogaster. eLife 2, e01179 (2013).

68 Jungreis, I. et al. Evidence of abundant stop codon readthrough in Drosophila and other metazoa. Genome research 21, 2096–2113 (2011).

69 Dishman, A. F. & Volkman, B. F. Unfolding the Mysteries of Protein Metamorphosis. ACS chemical biology 13, 1438–1446 (2018).

70 Forman-Kay, J. D. & Mittag, T. From sequence and forces to structure, function, and evolution of intrinsically disordered proteins. Structure 21, 1492–1499 (2013).

71 Burmann, B. M. et al. An alpha helix to beta barrel domain switch transforms the transcription factor RfaH into a translation factor. Cell 150, 291–303 (2012).

72 Tseng, R. et al. Structural basis of the day-night transition in a bacterial circadian clock. Science 355, 1174–1180 (2017).

73 Tuinstra, R. L. et al. Interconversion between two unrelated protein folds in the lymphotactin native state. Proceedings of the National Academy of Sciences of the United States of America 105, 5057–5062 (2008).

74 Domnauer, M. et al. Proteome plasticity in response to persistent environmental change. Molecular cell 81, 3294–3309 e3212 (2021).

75 Luo, X. & Yu, H. Protein metamorphosis: the two-state behavior of Mad2. Structure 16, 1616–1625 (2008).

76 Luo, X. et al. The Mad2 spindle checkpoint protein has two distinct natively folded states. Nature structural & molecular biology 11, 338–345 (2004).

77 Murzin, A. G. Biochemistry. Metamorphic proteins. Science 320, 1725–1726 (2008).

78 Dishman, A. F. et al. Evolution of fold switching in a metamorphic protein. Science 371, 86–90 (2021).

79 Chapple, C. E. et al. Extreme multifunctional proteins identified from a human protein interaction network. Nature communications 6, 7412 (2015).

80 Jeffery, C. J. Enzymes, pseudoenzymes, and moonlighting proteins: diversity of function in protein superfamilies. The FEBS journal 287, 4141–4149 (2020).

81 Jeffery, C. J. Moonlighting proteins. Trends in biochemical sciences 24, 8–11 (1999).

82 Schafer, J. W. & Porter, L. L. Evolutionary selection of proteins with two folds. Nature communications 14, 5478 (2023).

83 Wayment-Steele, H. K. et al. Predicting multiple conformations via sequence clustering and AlphaFold2. Nature 625, 832–839 (2024).

84 Chang, Y. G. et al. Circadian rhythms. A protein fold switch joins the circadian oscillator to clock output in cyanobacteria. Science 349, 324–328 (2015).

85 Sala, A. J., Bott, L. C. & Morimoto, R. I. Shaping proteostasis at the cellular, tissue, and organismal level. The Journal of cell biology 216, 1231–1241 (2017).

86 Kniepert, A. & Groettrup, M. The unique functions of tissue-specific proteasomes. Trends in biochemical sciences 39, 17–24 (2014).

87 Lopez, A., Fleming, A. & Rubinsztein, D. C. Seeing is believing: methods to monitor vertebrate autophagy in vivo. Open biology 8, 180106 (2018).

88 Mizushima, N., Yamamoto, A., Matsui, M., Yoshimori, T. & Ohsumi, Y. In vivo analysis of autophagy in response to nutrient starvation using transgenic mice expressing a fluorescent autophagosome marker. Molecular biology of the cell 15, 1101–1111 (2004).

## References

89 Gupta, R. et al. Firefly luciferase mutants as sensors of proteome stress. Nature methods 8, 879–884 (2011).

90 Kubala, M. H., Kovtun, O., Alexandrov, K. & Collins, B. M. Structural and thermodynamic analysis of the GFP:GFP-nanobody complex. Protein science : a publication of the Protein Society 19, 2389–2401 (2010).

91 Lam, S. S. et al. Directed evolution of APEX2 for electron microscopy and proximity labeling. Nature methods 12, 51–54 (2015).

92 Paez-Segala, M. G. et al. Fixation-resistant photoactivatable fluorescent proteins for CLEM. Nature methods 12, 215–218, 214 p following 218 (2015).

93 Selma-Soriano, E., Artero, R. & Llamusi, B. Optical Cross-Sectional Muscle Area Determination of Drosophila Melanogaster Adult Indirect Flight Muscles. Journal of visualized experiments : JoVE 134, e56179 (2018).

94 Huynh, K. & Partch, C. L. Analysis of protein stability and ligand interactions by thermal shift assay. Current protocols in protein science 79, 28 29 21-28 29 14 (2015).

95 Shevchenko, A., Tomas, H., Havlis, J., Olsen, J. V. & Mann, M. In-gel digestion for mass spectrometric characterization of proteins and proteomes. Nature protocols 1, 2856–2860 (2006).

96 Cammarata, M. B., Macias, L. A., Rosenberg, J., Bolufer, A. & Brodbelt, J. S. Expanding the Scope of Cross-Link Identifications by Incorporating Collisional Activated Dissociation and Ultraviolet Photodissociation Methods. Analytical chemistry 90, 6385–6389 (2018).

97 Jumper, J. et al. Highly accurate protein structure prediction with AlphaFold. Nature 596, 583–589 (2021).

98 Abramson, J. et al. Accurate structure prediction of biomolecular interactions with AlphaFold 3. Nature 630, 493–500 (2024).

99 Mirdita, M. et al. ColabFold: making protein folding accessible to all. Nature methods 19, 679–682 (2022).

100 Kahraman, A., Malmstrom, L. & Aebersold, R. Xwalk: computing and visualizing distances in cross- linking experiments. Bioinformatics 27, 2163–2164 (2011).

101 Micsonai, A. et al. BeStSel: a web server for accurate protein secondary structure prediction and fold recognition from the circular dichroism spectra. Nucleic acids research 46, W315–W322 (2018).

